# Contrasted gene decay in subterranean vertebrates: insights from cavefishes and fossorial mammals

**DOI:** 10.1101/2020.03.05.978213

**Authors:** Maxime Policarpo, Julien Fumey, Philippe Lafargeas, Delphine Naquin, Claude Thermes, Magali Naville, Corentin Dechaud, Jean-Nicolas Volff, Cedric Cabau, Christophe Klopp, Peter Rask Møller, Louis Bernatchez, Erik García-Machado, Sylvie Rétaux, Didier Casane

## Abstract

Evolution sometimes proceeds by loss, especially when structures and genes become dispensable after an environmental shift relaxing functional constraints. Gene decay can serve as a read-out of this evolutionary process. Animals living in the dark are outstanding models, in particular cavefishes as hundreds of species evolved independently during very different periods of time in absence of light. Here, we sought to understand some general principals on the extent and tempo of decay of several gene sets in cavefishes. The analysis of the genomes of two Cuban species belonging to the genus *Lucifuga* provides evidence for the most massive loss of eye genes reported so far in cavefishes. Comparisons with a recently-evolved cave population of *Astyanax mexicanus* and three species belonging to the tetraploid Chinese genus *Sinocyclocheilus* revealed the combined effects of the level of eye regression, time and genome ploidy on the number of eye pseudogenes. In sharp contrast, most circadian clock and pigmentation genes appeared under strong selection. In cavefishes for which complete genomes are available, the limited extent of eye gene decay and the very small number of loss of function (LoF) mutations per pseudogene suggest that eye degeneration is never very ancient, ranging from early to late Pleistocene. This is in sharp contrast with the identification of several eye pseudogenes carrying many LoF mutations in ancient fossorial mammals. Our analyses support the hypothesis that blind fishes cannot thrive more than a few millions of years in cave ecosystems.

## Introduction

The evolution of organisms confronted to drastic environmental shifts results in sometimes profound phenotypic changes. Constructive evolution involved in adaptation to new environments, and relying on novelties at phenotypic and genetic levels, has drawn much interest. Nevertheless, it becomes evident that regressive evolution, which is often non adaptive and which occurs by loss of structures and functions and corresponding genes, accounts for a non-negligible part of the evolutionary process (Lahti, et al. 2009; Albalat and Cañestro 2016). Here, we sought to better understand the modalities, extent, tempo and limits of molecular decay of several light-related genetic systems in subterranean vertebrates. It has been shown that several independent lineages of obligate fossorial mammals with degenerated eyes have lost many genes involved in visual perception (Kim, et al. 2011; Emerling and Springer 2014; Fang, Nevo, et al. 2014; Fang, Seim, et al. 2014; Emerling 2018). Cave vertebrates which are essentially cavefishes are other outstanding models to tackle these issues (Culver and Pipan 2009). However, the molecular decay of genes has not been surveyed at a genome-wide scale in relevant cavefish species. On the one hand, in the reference genome of *A. mexicanus* cavefish, no or only a couple of pseudogenes have been found among sets of genes which are eye specific, involved in the circadian clock, or else related to pigmentation (Protas, et al. 2006; Beale, et al. 2013; McGaugh, et al. 2014). Such maintenance of a very high proportion of functional genes most likely results from a very recent origin, no earlier than in the late Pleistocene, of cave populations (Fumey, et al. 2018). On the other hand, in the genomes of three fishes belonging to the genus *Sinocyclocheilus* (*S. grahami* which is a surface fish with large eyes, *S. anshuiensis* which is a blind cavefish and *S. rhinocerous* which is a small-eyed cavefish) many LoF mutations were found (Yang, et al. 2016), but their tetraploid genomes hampered the identification of those mutations that fixed in relation to the surface to cave shift. Indeed, after a whole-genome duplication (WGD), the pair of paralogs resulting from this process (ohnologs) are most often redundant and one of them can be pseudogenized without reducing fitness. Accordingly, *S. grahami* carries eye peudogenes like the eyeless *S. anshuiensis* and the small-eyed *S. rhinocerous,* but no thorough analysis of differential gene losses in relation to the level of eye degeneration has been performed (Yang, et al. 2016).

In order to examine the long term effect of life in caves on the molecular decay of large sets of genes involved in various light-dependent biological processes, genomes of fishes evolving in caves for a very long time and which did not undergo a recent WGD are required. Two clades of cavefishes (cave brotulas from Bahamas and Cuba) were previously identified in the genus *Lucifuga*, one comprising only blind cavefish species and the other only small-eyed cavefish species (García-Machado, et al. 2011). As no close surface relative has been identified up to now and large genetic distances were found between some species, within and between clades, this genus of cavefishes is likely relatively ancient, and the last common ancestor of extant species was probably a cave-adapted fish. We sequenced the genomes of two Cuban cave brotulas: one specimen, belonging to *L. dentata*, was blind and depigmented, the other one, belonging to *L. gibarensis*, had small eyes and was pigmented. The latter species is a new species first identified as *Lucifuga* sp. 4 (García-Machado, et al. 2011) that will be formally named and described in a forthcoming publication.

We searched for likely LoF mutations (*i.e.* STOP codon gains, losses of START and STOP codons, losses of intron splice sites and small indels leading to frameshifts) and for several signatures of relaxed selection on nonsynonymous mutations in genes: 1) uniquely expressed in the eyes or coding for non-visual opsins, 2) involved in the circadian clock, 3) involved in pigmentation. The contrasted patterns of pseudogenization found for the three categories of genes indicate that eye genes are much less constrained than circadian clock and pigmentation genes in caves. In *A. mexicanus* cavefish, despite only one eye gene carrying a LoF mutation was found, using machine learning-based estimations of the deleterious impact of nonsynonymous mutations implemented in MutPred2 (Pejaver, et al. 2017), we obtained evidence that most if not all eye genes are under relaxed selection, but for a too short period of time to allow the fixation of more than a few LoF mutations. In other cavefishes, more eye pseudogenes were found and the level of gene decay depended on several factors such as the time fishes have spent in the subterranean environment, their level of troglomorphy and the level of ploidy of their genomes. Nevertheless, no eye genes with many LoF mutations were found, in sharp contrast to highly degenerated eye genes identified in some fossorial mammals, suggesting that eye degeneration in cavefishes is much more recent.

## Results

### Assembly of the draft genomes of two Cuban cave brotulas

First, the genome of a specimen of *Lucifuga dentata* (Bythitidae, Ophidiiformes) was sequenced (see a photo in **supplementary fig. S1, Supplementary Material** online). Assembly resulted in 52,944 scaffolds whose size sum up to 634 Mb, N50 = 119.6 kb (for scaffold size distribution, see **supplementary fig. S2, Supplementary Material** online). This genome size is consistent with those of three other genomes available (Malmstrøm, et al. 2017) and estimations of the genome size of other Ophidiiformes (Gregory 2019). To assess the quality of the assembly, raw sequences were realigned to the assembly: 95% of the reads realigned correctly resulting in a mean coverage of 134x. Then, the completeness of the assembly was assessed using BUSCO with the Actinopterygii gene database (Kriventseva, et al. 2015). Among 4,584 genes, 4,249 (92.7%) were found complete, 194 (4.2%) were incomplete and 141 (3.1%) were missing. Using BUSCO with three other Ophidiiformes genomes currently available (*Brotula barbata*, *Carapus acus* and *Lamprogrammus exutus*), the genome of *Lucifuga dentata* appeared as the most complete (see **supplementary fig. S3, Supplementary Material** online). Then, the genome of a specimen belonging to the small-eyed *Lucifuga gibarensis* was sequenced (see a photo in **supplementary fig. S1, Supplementary Material** online). As nuclear DNA sequence divergence is about 1% between the two *Lucifuga* species, short reads of *L. gibarensis* were mapped on *L. dentata* genome. The mean coverage was 84x, with 86% of the reads mapping on the genome. Heterozygosity was estimated on raw Illumina reads using GenomeScope (Vurture, et al. 2017). The heterozygosity of *L. dentata* (0.1%) was lower than that of *L. gibarensis* (0.26%).

### Assembly of a transcriptome of *L. dentata* and genome annotation

Based on mRNA extracted from the gonads, gills, heart and brain of *L. dentata*, a *de novo* transcriptome assembly was obtained using Trinity (Grabherr, et al. 2011). Quality and completeness assessment of this transcriptome were performed following Trinity guide. Among 4,584 genes corresponding to the Actinopterygii gene database of BUSCO, 82.8% were found complete (**supplementary fig. S3, Supplementary Material** online) and 92.31 % of the reads were mapped back to the assembly with 84 % as proper pairs, which indicate an overall good quality transcriptome. More on quality check can be found in **supplementary fig. S4, Supplementary Material** online.

A combination of *de novo* predictions, RNA-seq evidence and protein alignments was used to annotate the genome of *L. dentata* (see workflow in **supplementary fig. S5, Supplementary Material** online). This resulted in 30,001 gene models with an average gene length of 9,693 bp and an average protein length of 435 amino acids. Among predicted genes, 23,524 had a functional annotation with BLAST to the SwissProt/UniProt database and 21,558 genes were detected with a functional domain by Interproscan. Annotation completeness was assessed using BUSCO in protein mode; among 4,584 corresponding to the Actinopterygii gene database of BUSCO, 87.4% were found complete, 6.5% incomplete and 6% missing (**supplementary fig. S3, Supplementary Material** online). A homemade pipeline was used to describe the repeat landscape of the genome of *L. dentata*. We found 16.3% of repeated elements, among which 2.4% of LINEs and 0.4% of SINEs (**supplementary fig. S6, Supplementary Material** online).

### Delimitation and retrieving of eye, circadian clock and pigmentation genes

In zebrafish, *Danio rerio,* we identified 95 genes expressed only in the eyes or coding non-visual opsins expressed in other organs (fig. 1A**, supplementary fig. S7** and **Data Supp 1, Supplementary Material** online, and see Methods). In addition, we retrieved a list of 42 circadian clock genes (Li, et al. 2013) and 257 genes involved in pigmentation (Lorin, et al. 2018) (fig. 1B and fig. 1C**, supplementary Data_Supp1, Supplementary Material** online). Using the program exonerate, homologs were retrieved from other fish genomes, that is five cavefishes (*A. mexicanus* from Pachón cave, *L. dentata* and *L. gibarensis*, *S. anshuiensis* and *S. rhinocerous*), close surface relatives (*A. mexicanus* and *Pygocentrus nattereri*, *Brotula barbata*, *Carapus acus* and *Lamprogrammus exutus, S. grahami* and *C. carpio)* and a distantly-related outgroup (*Lepisosteus oculatus*). Their phylogenetic relationships are shown in fig. 2. Noteworthy, some genes have been duplicated in the terminal lineage leading to zebrafish (used as a reference to establish the gene lists) and thus only one copy was expected to be found in other fishes. On the other hand, gene duplications, gene deletions as well as WGDs occurred in other lineages. Therefore, the number of genes retrieved is highly variable among genomes (fig. 3).

**Fig. 1.**
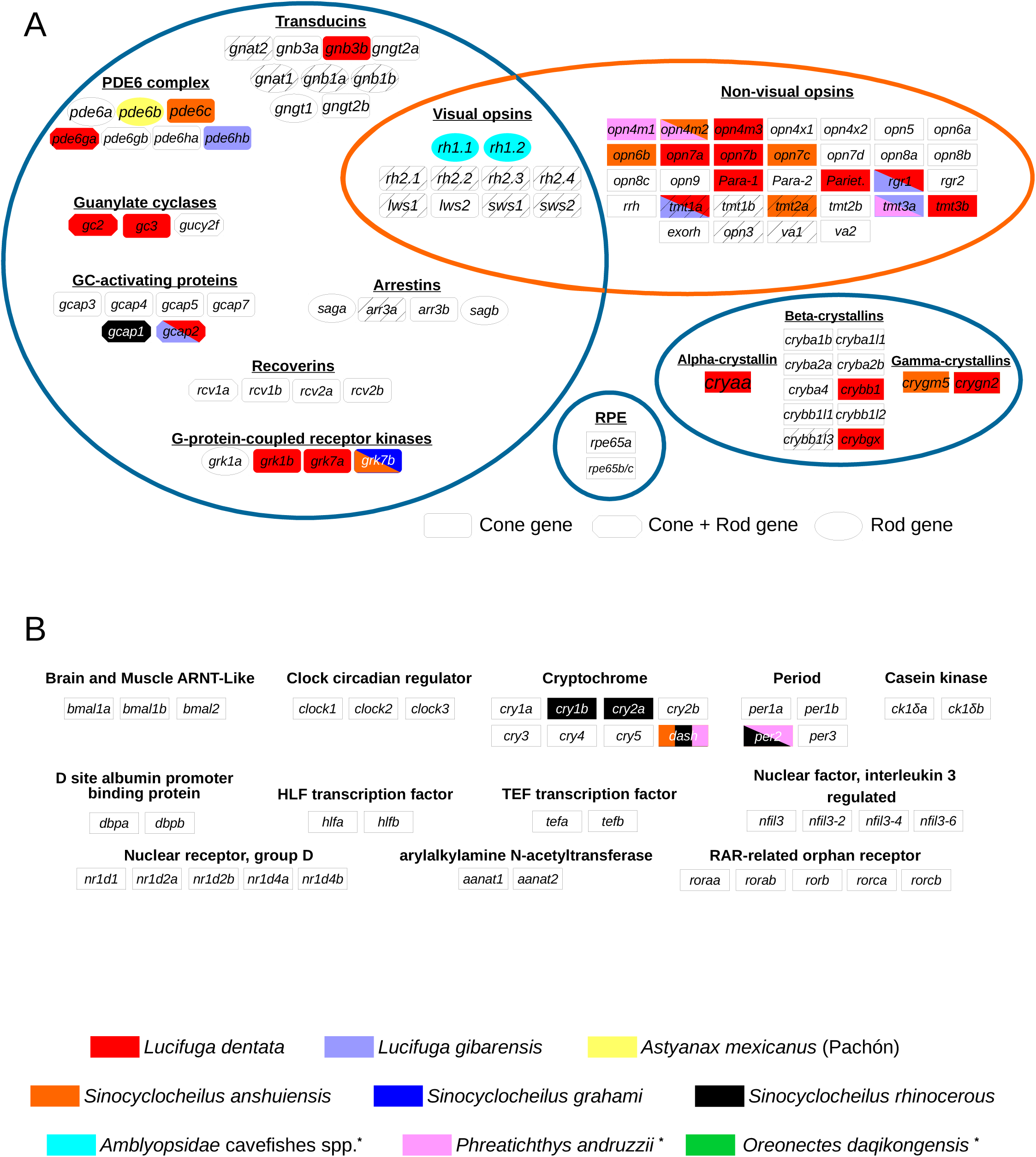
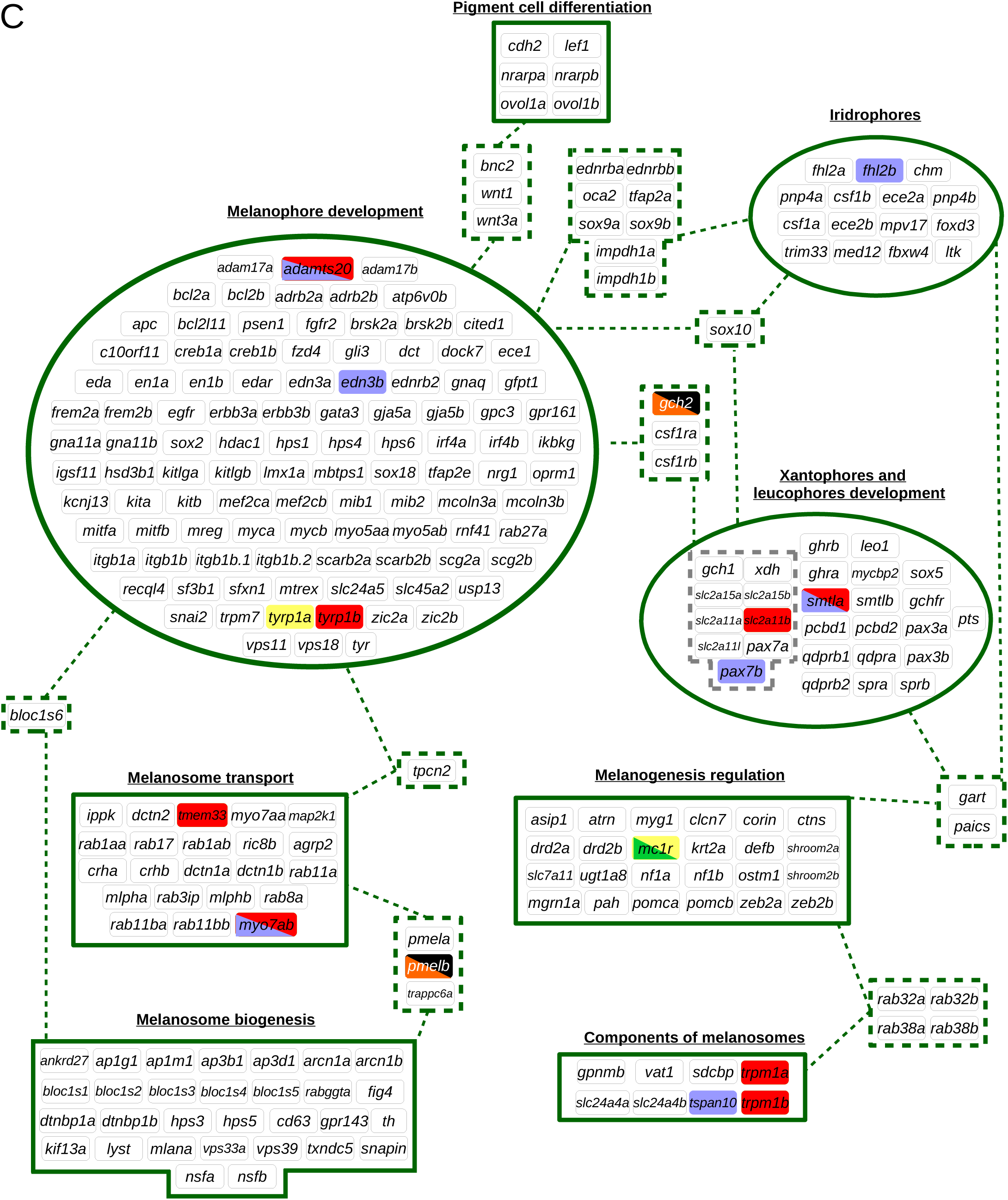
Gene sets. (A) Eye genes. (B) Circadian clock genes. (C) Pigmentation genes. Genes were colored according to the species in which LoF mutations were found (species name followed by * indicate that no genome was available but pseudogenes were identified). Genes with a hatched background are under/not expressed in at least one cavefish species. Genes expressed only in the eyes are surrounded by blue lines and opsins by an orange line. Pigmentation genes were clustered according to Lorin et al. 2018. Genes belonging to several subsets are surrounded by dotted lines with links between the different subsets.

**Fig. 2.**
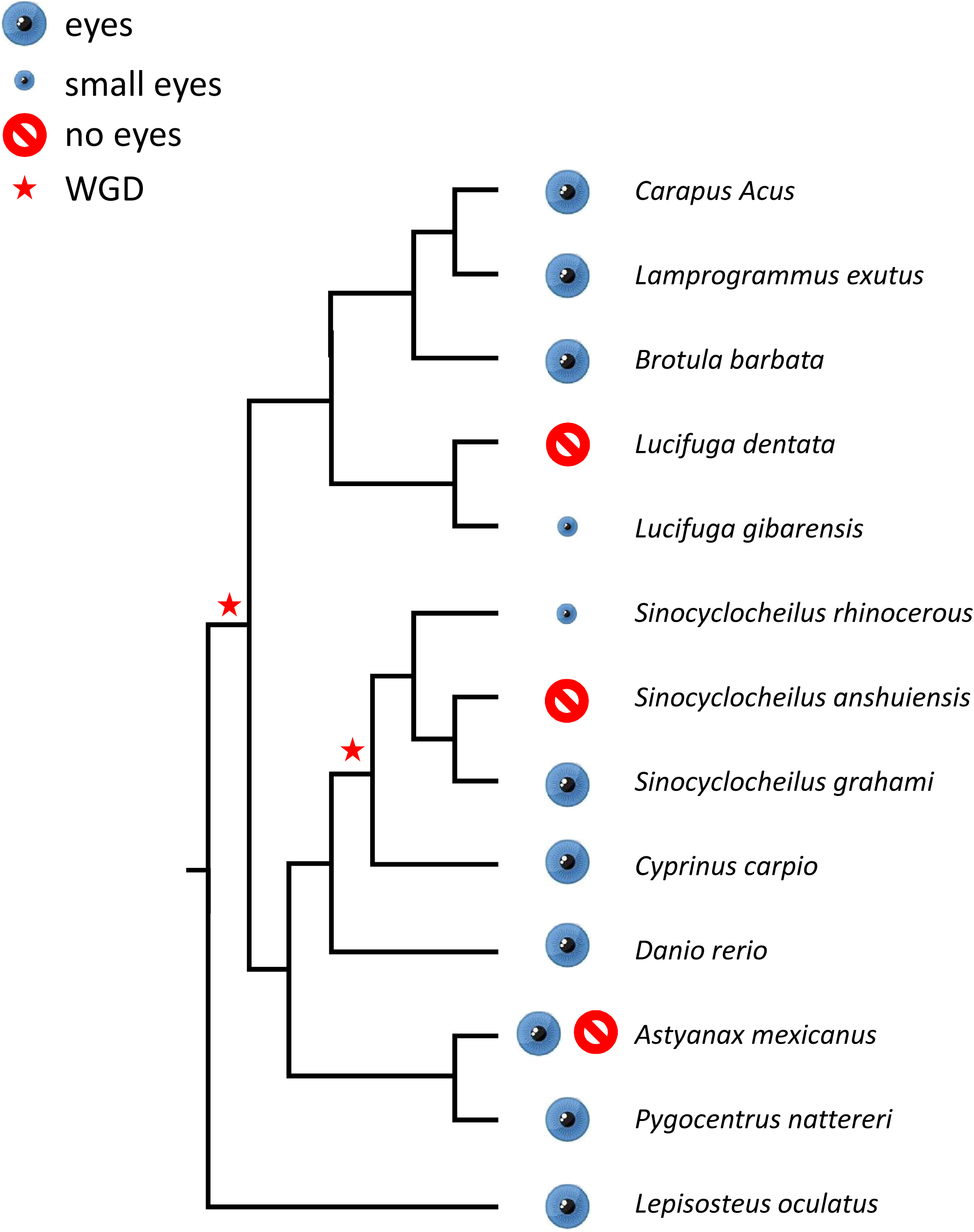
Phylogeny of the cavefishes and some close relatives.

**Fig. 3.**
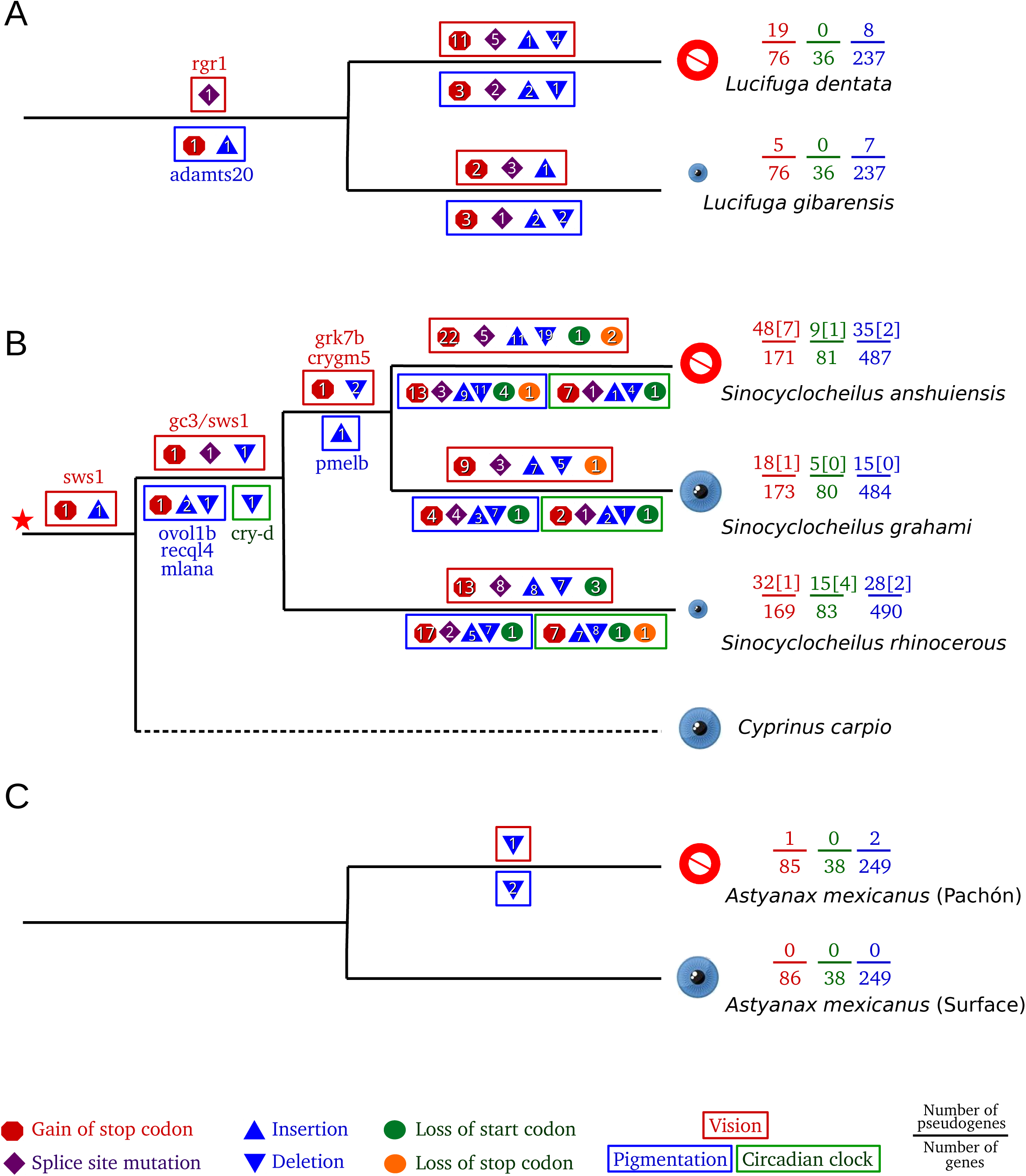
Mapping of LoF mutations. For *Sinocyclocheilus* species, the number of genes for which both ohnologs are pseudogenized is given between brackets.

### Identification of LoF mutations

Genes sequences were classified as functional if found complete with no LoF mutation, as pseudogene if complete and carrying at least one LoF mutation, and as truncated if incomplete (the sequences can be found in **supplementary Data_Supp2, Supplementary Material** online). Only the following LoF mutations were analyzed: gain of an internal STOP codon, loss of the initiation codon, loss of the STOP codon, indel leading to a frameshift, mutations at intron donor and acceptor sites. In the present study, incomplete genes were discarded as it was difficult to know if they corresponded to sequencing gaps, assembly artefacts or true large deletions. Using PCR to amplify missing exons, we estimated that 85% of the large deletions in the *A. mexicanus* cavefish genome are artefacts (data not shown) - although some large deletions such as in the gene *Oca2* (a pigmentation gene) are real (Protas, et al. 2006). Other mutations in non-coding and coding sequences that could lead to a non-functional gene were not searched for as they cannot be readily identified. For example, several in-frame indel mutations were found in *A. mexicanus* but their functional consequences remained elusive (Berning, et al. 2019). The numbers of pseudogenes reported hereafter are thus underestimates of the true numbers of non-functional genes, but they nevertheless allowed comparative analyses.

<u>Eye pseudogenes</u>: among the list of 95 zebrafish eye genes, 76 genes were retrieved from *Lucifuga* genomes, 75 from *B. barbata*, 72 from *C. acus* and 73 from *L. exutus* (fig. 3, **Supplementary fig. S7** and **Data_Supp1, Supplementary Material** online). Interestingly, all these ophidiiforms seem to have lost long-wave sensitive (LWS) opsins. This loss is most likely due to a gene deletion in their common ancestor living in deep ocean (**supplementary fig. S8, Supplementary Material** online), in accordance with a report on the reduction of the number of LWS genes in fishes living below 50 m (Lin, et al. 2017). While no eye pseudogene was found in *B. barbata* or *C. acus* and only one in *L. exutus* (*gcap1*), 5 pseudogenes were identified in *L. gibarensis* and 19 pseudogenes in *L. dentata*. The non-visual opsin *rgr1* was pseudogenized in the common ancestor of the two *Lucifuga* species, as the same mutation (at a splice site of intron 4) was found in both genomes (fig. 3 **and supplementary Data_Supp2, Supplementary Material** online). Examination of the read coverage of LoF mutations indicated that the specimen of *L. gibarensis* sequenced was heterozygous for LoF mutations found at two different sites in the *gcap2* gene (**supplementary table S1**, **Supplementary Material** online). In the transcriptome of *L. dentata*, transcripts corresponding to 9 pseudogenes were found (3 non-visual opsins, 3 crystallins and 3 genes involved in the phototransduction pathway), while no transcripts were found for 10 other pseudogenes (**supplementary table S1**, **Supplementary Material** online). In those transcripts, all the LoF mutations identified at the genome level were present. In agreement with a recent WGD, two copies (ohnologs) of most eye genes were retrieved from the genomes of *Sinocyclocheilus* species (fig. 3, **supplementary fig. S7, Supplementary Material** online). In the large-eyed *S. grahami*, about 10% of retrieved eye genes were pseudogenized (18 / 173 genes carried at least a LoF mutation), to be compared to 19% (32 / 169) in the small-eyed *S. rhinocerous* and 28% (48 / 171) in the eyeless cavefish *S. anshuiensis*. Only one pair of ohnologs were concomitantly pseudogenized in the eyed *S. grahami* and the small-eyed *S. rhinocerous,* while seven pairs of ohnologs were concomitantly pseudogenized in the blind *S. anshuiensis* (fig.1, fig.3**, supplementary fig. S7, Data_Supp1, Supplementary Material** online). A STOP codon and a frameshift in *sws1* were shared by the three *Sinocyclocheilus* species and *Cyprinus carpio*. A new STOP codon and a new frameshift in this gene were shared by *Sinocyclocheilus* species, as well as a mutation at the donor site of the third intron of *gc3*; *S. anshuiensis* and *S. grahami* shared a frameshift in *crygm5* and a frameshift and a new STOP codon in *grk7b* (fig. 3). In *A. mexicanus*, 86 genes were retrieved from the surface fish genome while 85 were retrieved from the genome of the Pachón cavefish. Only one pseudogene was found in the Pachón cavefish genome, which is due to a deletion of 11 bp in *pde6b* (fig. 1 and fig. 3). The examination of the automatic annotation of the gene allowed the identification of an erroneous 1 bp intron (ENSAMXG00000000290, Ensembl 91), restoring the coding frame. Noteworthy, we also confirmed using PCR that two large deletions occurred in the Pachón cavefish, one removing *opn8b* and the last 3 exons of *opn8a,* the other eliminating 2 exons of *rgr2*. However, these genes were not included in the list of pseudogenes, according to the restrictive definition of an identifiable pseudogene used in the present study. In summary, while no or very few eye genes are pseudogenized in surface fishes and *A. mexicanus* cavefish, more eye pseudogenes were found in other cavefishes, up to 25% in *L. dentata*.

<u>Circadian clock pseudogenes</u>: based on a literature survey, 42 genes involved in the circadian clock in *Danio rerio* were identified (Li, et al. 2013) and retrieved from other fish genomes. On the one hand, no pseudogene was found among 36 genes retrieved from *Lucifuga* genomes and 38 genes identified from *Astyanax* genomes. On the other hand, 5, 15 and 9 pseudogenes were identified among 80, 83 and 81 genes retrieved from the genomes of *S. grahami* (eyed), *S. rhinocerous* (small-eyed) and *S. anshuiensis* (blind), respectively. Both ohnologs of *cry-dash* were independently pseudogenized in *S. rhinocerous* and *S. anshuiensis*, a gene also pseudogenized in the Somalian cavefish *P. andruzzii*, Three other pair of ohnologs (*cry1b*, *cry2a* and *per2*) carried LoF mutations in *S. rhinocerous*, *per2* being also non-functional in *P. andruzzii*. These data suggest that the circadian clock has most likely been lost in *S. rhinocerous* whereas this loss is less strongly supported in the case of *S. anshuiensis* (fig. 1 and fig. 3).

<u>Pigmentation genes</u>: based on a literature survey, 257 genes involved in pigmentation in *Danio rerio* were identified (Lorin, et al. 2018) and retrieved from other fish genomes. Very few pseudogenes were found among 237 pigmentation genes in *Lucifuga* genomes, that is 8 LoF mutations in *L. dentata* and 7 in *L. gibarensis*. While *smtla* and *myo7ab* seem to have been lost independently in the two lineages, a STOP codon and an insertion is shared in *adamts20*. The number of pseudogenes in these cavefishes does not seem to depart from those found in some surface relatives, as 6 pseudogenes were identified among 230 pigmentation genes in *Lamprogrammus exutus* (**supplementary Data_Supp1, Supplementary Material** online). Among *Sinocyclocheilus* species, only 3% (15/484) of pseudogenes were found in *S. grahami* while 6% (28/490) were found in *S. rhinocerous* and 7% (35/487) in *S. anshuiensis* (fig. 3). Thus, after the WGD, the retention of pigmentation genes seems to have been much higher than among eye genes in the two cavefishes but also in the surface fish (compare to 10%, 19% and 28% of eye pseudogenes, respectively). Such high percentage of retention of pigmentation genes has been found also after the Salmonid-specific WGD (Lorin, et al. 2018). Strikingly, while no pair of ohnologs was found pseudogenized in *S. grahami*, the same two pairs of ohnologs (*gch2* and *pmelb*) were independently pseudogenized in *S. anshuiensis* and *S. rhinocerous*. The very small number of pseudogenes and the independent pseudogenization of the same genes in these two species suggest that only a limited subset of genes involved in pigmentation can be lost in these cavefishes.

In *A. mexicanus*, 2 pseudogenes were found among 249 pigmentation genes: *mc1r* which has already been reported in the literature (Gross, et al. 2009) and which is also pseudogenized in the Chinese cavefish *Oreonectes daqikongensis* (Liu, et al. 2019), and *tyrp1a* that is mis-annotated in ensembl (ENSAMXG00000021619, Ensembl 91).

### Types of LOF mutations and their distribution along pseudogenes

A total of 118 mutations to a STOP codon, 148 frameshifts (84 deletions and 64 insertions), 5 STOP codon losses, 13 START codon losses and 40 intron splice site losses were identified in our dataset (fig. 4) (see **supplementary Data_Supp1**, **Supplementary Material** online for a detailed description of the number of LoF mutations in each gene set). Most frameshifts were the results of very small deletions or insertions (1 or 2 bp) while a few of them were indels involving a larger number of nucleotides (fig. 4C**, supplementary fig. S10**, **Supplementary Material** online). The largest deletion was 83 bp long in *zic2b* (pigmentation gene) of *S. rhinocerous* and the largest insertion was 20 bp long in *opn7b* (eye gene) of *S. anshuiensis*. In order to test if LoF mutations were distributed randomly along the genes, that is they were not clustered at the 3’ end of the genes where their deleterious effect could be low, we computed the effective segment size generated by LoF mutations and compared this value with simulations of random distributions of mutations along genes. We found that premature STOP codons and frameshifts are distributed randomly along coding sequences (for more details, see Materials and Methods and **supplementary fig. S9**, **Supplementary Material** online). We next tested if the relative frequencies of the different types of LoF mutations (*i.e.* STOP codon gains, losses of START and STOP codons, losses of intron splice sites and small indels leading to frameshifts) were those expected under the neutral model, that is if their relative frequencies were proportional to their probabilities of occurrence. Taking into account a nucleotide mutation rate (μ), observed transition/transversion ratio and codon frequencies, the rate of mutation to a new STOP codon was μ_stop_ = 0.036μ. Based on the ratio of frameshift/STOP mutations, we estimated the rate of indels leading to frameshifts (μ_frameshift_) as 148/118 x μ_stop_ = 0.05μ. The rate of mutations in splice acceptor or donor sites was estimated as: 4 x (number of introns) / Σ (CDS length) x μ, *i.e.* 4 x 34175 / 6154965 x μ = 0.022μ (where 34175 is the number of introns and 6154965 is the number of bases identified in the 3625 genes retrieved from the genomes of two *Lucifuga* species, three *Sinocyclocheilus* species and two genomes of *Astyanax mexicanus*). The rate of START codon loss was estimated as: 3 x (number of genes) / Σ (CDS length) x μ, *i.e.* 3 x 3625 / 6154965 x μ = 0.0018μ. The rate of STOP codon loss was estimated as: 3 x (number of genes) / Σ (CDS length) x 0.85 x μ, *i.e.* 3 x 3625 / 6154965 x 0.85 x μ = 0.0015μ (where 0.85 is the probability that a mutation in a STOP codon leads to a sense codon). The observed distribution of LoF mutations fitted well with those expected, either taking the three datasets together (fig. 4B**, supplementary fig. S11** and **Data_Supp1**, **Supplementary Material** online) or each dataset individually. It was similar to the distribution found in eye pseudogenes of subterranean mammals (**supplementary fig. S11 and supplementary Data_Supp1**, **Supplementary Material** online). These results suggested that the number LoF mutations of each type is proportional to its probability of occurrence.

**Fig. 4.**
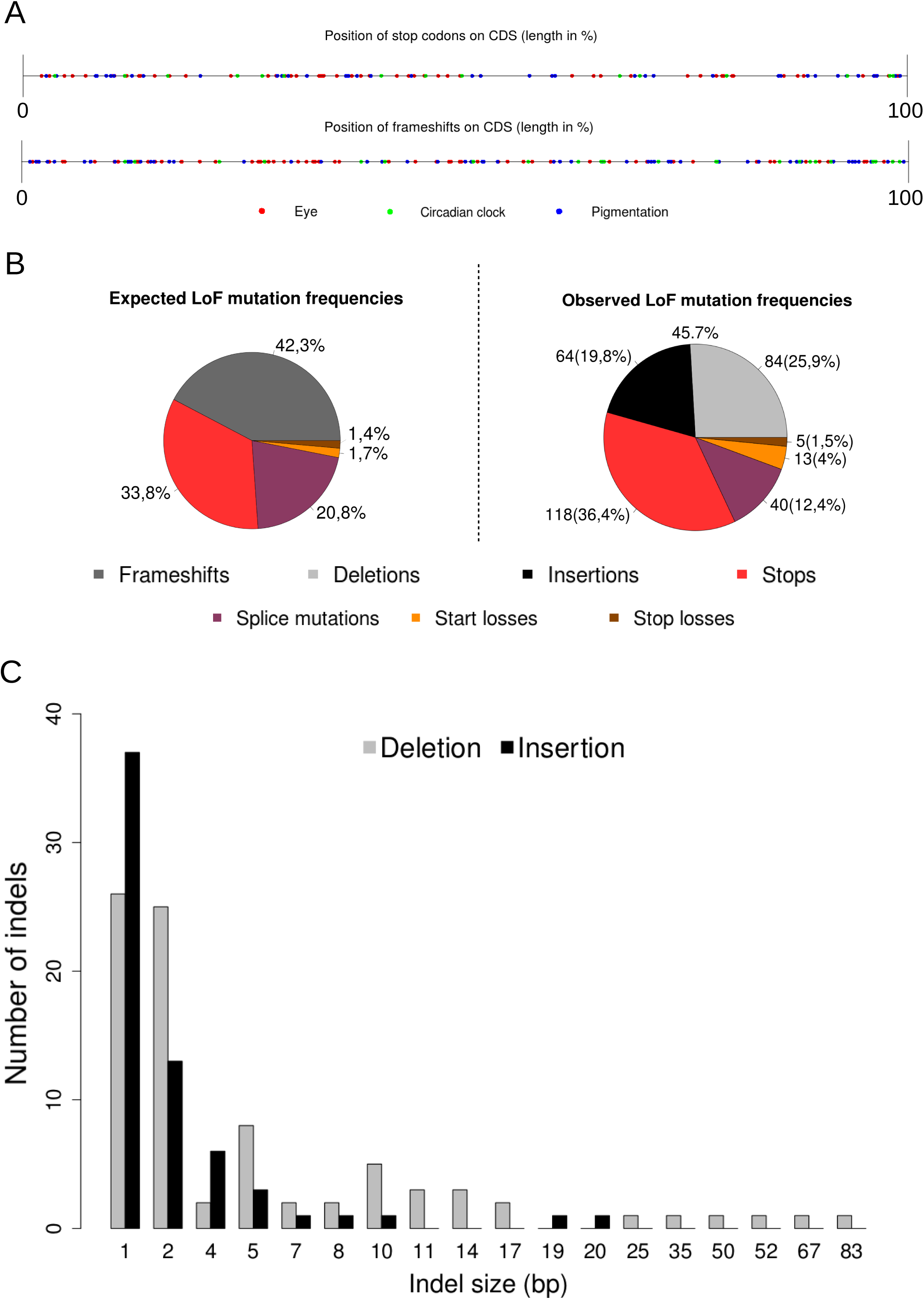
Distribution of different categories of LoF mutations. (A) Position of internal stop codons and frameshifts along coding sequences. (B) Observed and expected frequencies. (C) Distribution of indel size.

### Estimation of the number of effectively neutral eye genes based on the distribution of LoF mutations per pseudogene in cave brotulas

Among pseudogenes, some accumulated more than one LoF mutation, but in most of the cases only one LoF mutation was found (**supplementary fig. S7 and Data_Supp1**, **Supplementary Material** online). In order to test if the whole set, or only a subset, of eye genes is free to accumulate LoF mutations, we compared the distribution of the number of LoF mutations per pseudogene with those expected under these different hypotheses.

Expected distributions were obtained using either a simple analytical model assuming that all genes have the same probability to fix a LoF mutation, or a more complex model that takes into account that different genes do not have the same probability to fix a LoF mutation because they have different length and they do not contain the same number of introns. In the latter case, the computation of expected distribution was based on simulations. Very similar expected distributions were obtained with both approaches. This analysis could be performed only with *Lucifuga* species, as only one LoF mutation was found in *Astyanax mexicanus* and a WGD allowed LoF mutations to reach fixation in a *Sinocyclocheilus* species with large functional eyes such as *S. grahami*.

In *L. dentata*, 22 LoF mutations were distributed among 19 eye pseudogenes. More precisely, among the 76 genes retrieved, there were 57 genes without LoF mutation, 16 with 1 mutation, and 3 with 2 mutations (fig. 3 and **supplementary fig. S7**, **Supplementary Material** online). This distribution was compared with expected distributions obtained for different numbers of neutral genes ranging from 19 to 76 (fig. 5A). The best fit between the observed and expected distribution was found when at least 60 genes are evolving as neutral sequences (fig. 5A). Using the same approach, in *L*. *gibarensis*, we analysed the observed distribution of the number of LoF mutations per pseudogene (71 genes without LoF mutation, 3 with 1 mutation, 2 with 2 mutations), considering a number of neutral genes within a range of 5 to 76 (fig. 5B and **supplementary fig. S7**, **Supplementary Material** online). In this case, the best fit was obtained when about 15 eye genes are free to accumulate LoF mutations (fig. 5B). These results suggested that most genes, if not all, are dispensable in the blind *L. dentata* whereas only a small subset can be lost in the small-eyed *L. gibarensis*.

**Fig. 5.**
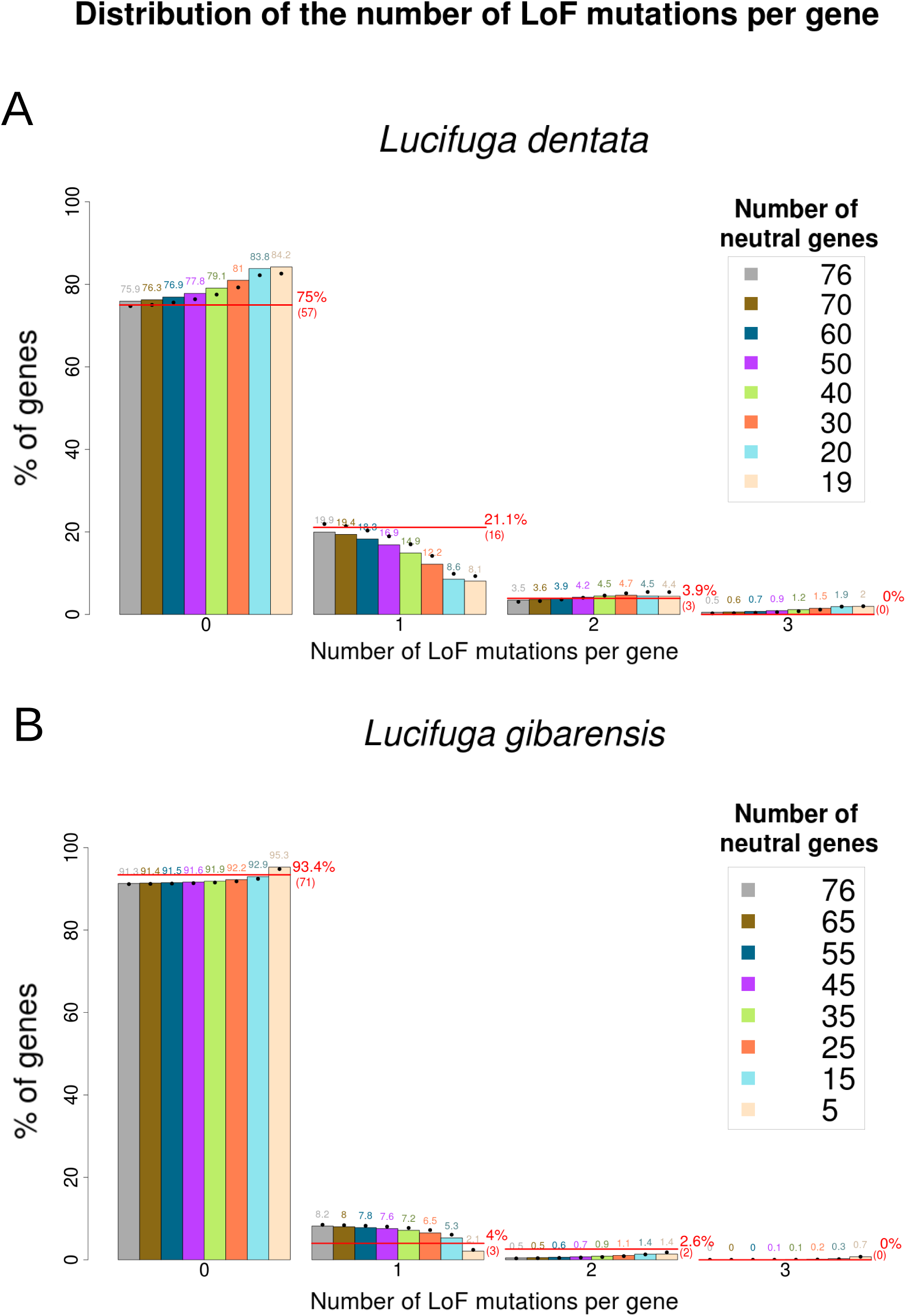
Observed and expected distributions of LoF mutations per gene. (A) *L. dentate*. (B) *L. gibarensis*. Red line: observed distribution. The expected distributions were obtained using an analytical model (dots) and 10,000 simulations (histograms).

### Evidence of relaxed selection on non-synonymous mutations in cavefish eye genes

To reinforce the evidence brought by the above analyses on LoF mutations, we looked for other signatures of relaxed selection using methods based on changes in ω (the ratio of the mean number of nonsynonymous substitutions per nonsynonymous site to the mean number of synonymous substitutions per synonymous site, also known as dn/ds). It is expected to be lower than one under purifying selection, equal to one under neutral evolution, and larger than one under adaptive selection. As gene divergence between *Lucifuga dentata* and *Lucifuga gibarensis* was lower than 0.9% and lower than 0.2% between the two *Astyanax mexicanus* morphs (for more details, see **supplementary folder divergence_values**, **Supplementary Material** online), the number of nucleotide differences per gene was very low and often no sequence change was observed between a cave species (or population) and the closest surface species (or population) (**supplementary fig. S12**, **Supplementary Material** online). Therefore, we used three sets of concatenated gene sequences (eye, circadian clock and pigmentation genes) to compute ω.

With the phylogenetic analysis using maximum likelihood (PAML) package Version 4.9h (Yang 2007), allowing a different ω along each branch, *Lucifuga dentata* had the highest ω (0.409) for eye genes. For circadian clock genes, both *Astyanax mexicanus* cavefish and *Lucifuga dentata* had the highest ω (0.29). For pigmentation genes, ω was similar in cave and surface fishes (fig. 6ABC, **supplementary fig. S13**, **Supplementary Material** online). Independently, we computed ω for the same sets of genes in *Sinocyclocheilus* species. For each species, ohnologs were concatenated into two series of gene sequences. With eye genes, ω was higher in the blind *S. anshuiensis* (0.36) than in the small eyed *S. rhinocerous* (0.32) and the eyed *S. grahami* (0.23). With circadian clock genes, ω has higher in the blind *S. anshuiensis* (0.38) and the small eyed *S. rhinocerous* (0.37) than in the eyed *S. grahami* (0.25). With pigmentation genes ω was higher in the small eyed *S. rhinocerous* (0.32) and the blind *S. anshuiensis* (0.29) than in the eyed *S. grahami* (0.25) (**supplementary fig. S13**, **Supplementary Material** online). Thus ω was consistently higher in cavefishes than in surface fishes, the shift being larger for eye genes than for circadian and pigmentation genes. In order to further examine if the shift of ω in some cavefishes for some sets of genes revealed a relaxed selection, we used another approach implemented in RELAX which computes the values and distribution of three ω using a branch-site model, the convergence of the three ω towards one in a lineage being a signature of relaxed selection (Wertheim, et al. 2015). The magnitude of convergence depends on a parameter, k, which tends to zero as selection tends to complete relaxation. RELAX detected relaxed selection on *Lucifuga dentata* eye genes with an important shift toward ω = 1 (k = 0.2), and this was also true in a lesser extent in *A. mexicanus* cavefish (k = 0.5). For pigmentation genes, the largest shift was also observed in *Lucifuga dentata* (k = 0.48). No shift of distribution was observed with cavefish circadian clock genes, suggesting that these genes are under strong purifying selection (**supplementary fig. S14, fig. S15** and **fig. S16 Supplementary Material** online).

**Fig. 6.**
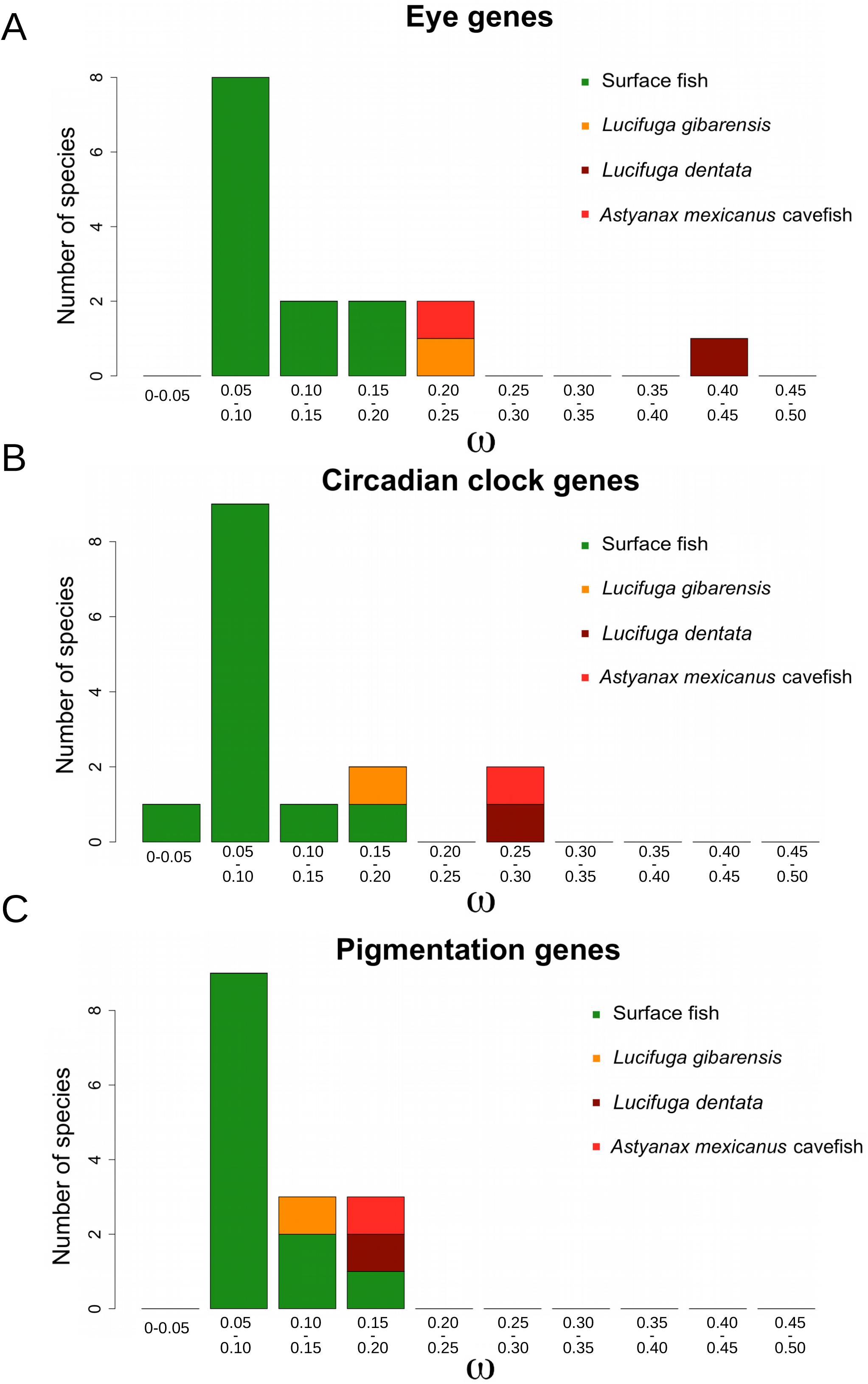
Distribution of ω in surface and cave fishes. (A) Eye genes. (B) Circadian clock genes. (C) Pigmentation genes.

Finally, with the aim of finding independent evidence of relaxed selection in cavefishes, in particular on *A. mexicanus* eye genes for which the number of mutations is low and thus the estimation of ω was not accurate, a novel approach was developed. First, nonsynonymous mutations in different lineages were inferred using the aaml program from the PAML package. Then, the deleterious impact of these mutations, a score which ranges between 0 (not deleterious) and 1 (very deleterious), was estimated using a machine learning method implemented in MutPred2 (Pejaver, et al. 2017). The kernel density estimation (KDE) of the distributions of the scores in eye, circadian clock and pigmentation genes were obtained for each terminal lineages leading to surface fishes and cavefishes, as well as for computer simulations of substitutions in the same sets of genes under a neutral model. Whatever the set of genes, in all surface fishes, the KDE was similarly right-skewed (fig. 7ABC), suggesting that most mutations which reached fixation have a low impact on the fitness. This was confirmed by the shape of the distribution of the scores in simulations of substitutions without selection (equivalent to the distribution before selection) which was very different to those of surface fishes, that is almost uniform and suggesting that the most deleterious mutations had been removed by selection in surface fishes. Before selection, the score distribution was slightly different for the different sets of genes, probably reflecting different selective constraints on the sequences belonging to these gene sets (fig. 7ABC, grey and black curves). Noteworthy, the Transitions/Transversions (Ts/Tv) ratio used in simulations of substitutions under a neutral model had no impact on the distribution of the scores (**supplementary fig. S19, Supplementary Material** online). In cavefishes, the score distribution was very variable, depending on the cavefish species and the set of genes (fig. 7ABC). In order to refine the analysis of the score distribution in cavefishes, admixtures of different proportions of substitutions picked up from two distributions, one under neutral evolution (from the simulations) and the other with selection (in the lineage leading to zebrafish) were also obtained to make comparisons with cavefish distributions. Pairwise comparisons of empirical cumulative distributions (ECDF) were performed using the nonparametric Kolmogorov-Smirnov (KS) test. The same approach was attempted using Grantham’s distances (Grantham 1974) instead of MutPred2 scores but the contrast between the distributions of the distances with and without selection was much less discriminant and not analyzed further (**supplementary fig. S18, Supplementary Material** online).

<u>With eye genes</u>, for *A. mexicanus* cavefish (red curve, fig. 7), the distribution was not statistically different from that expected if all substitutions where neutral in this lineage (KS test, p = 0.2; **supplementary fig. S17, Supplementary Material** online), yet the best fit was with a mixture distribution with 24% of substitutions from the distribution under selection (**supplementary fig. S20, Supplementary Material** online). For *L. dentata* (brown curve, fig. 7) and *L. gibarensis* (orange curve, fig. 7), distributions departed from the neutral distribution (KS test, p = 1.4 x 10^-5^ and p = 4 x 10^-6^ respectively) (fig. 7A and **supplementary fig. S17, Supplementary Material** online) and the best fit was obtained with respectively 34% and 60% of the substitutions from the distribution under selection (**supplementary fig. S20, Supplementary Material** online). For all *Sinocyclocheilus* species, the score distribution was different from those of surface fishes, even for the eyed *S. grahami*, most likely because after the WGD purifying selection on nonsynonymous mutations was partially relaxed on one or both ohnologs, but the ECDF of *S. rhinocerous* and *S. anshuiensis* were more shifted toward the neutral distribution than the ECDF of *S. grahami*, suggesting that the two cavefishes experienced a more neutral regime than the surface fish (**supplementary fig. S21, Supplementary Material** online).

**Fig. 7.**
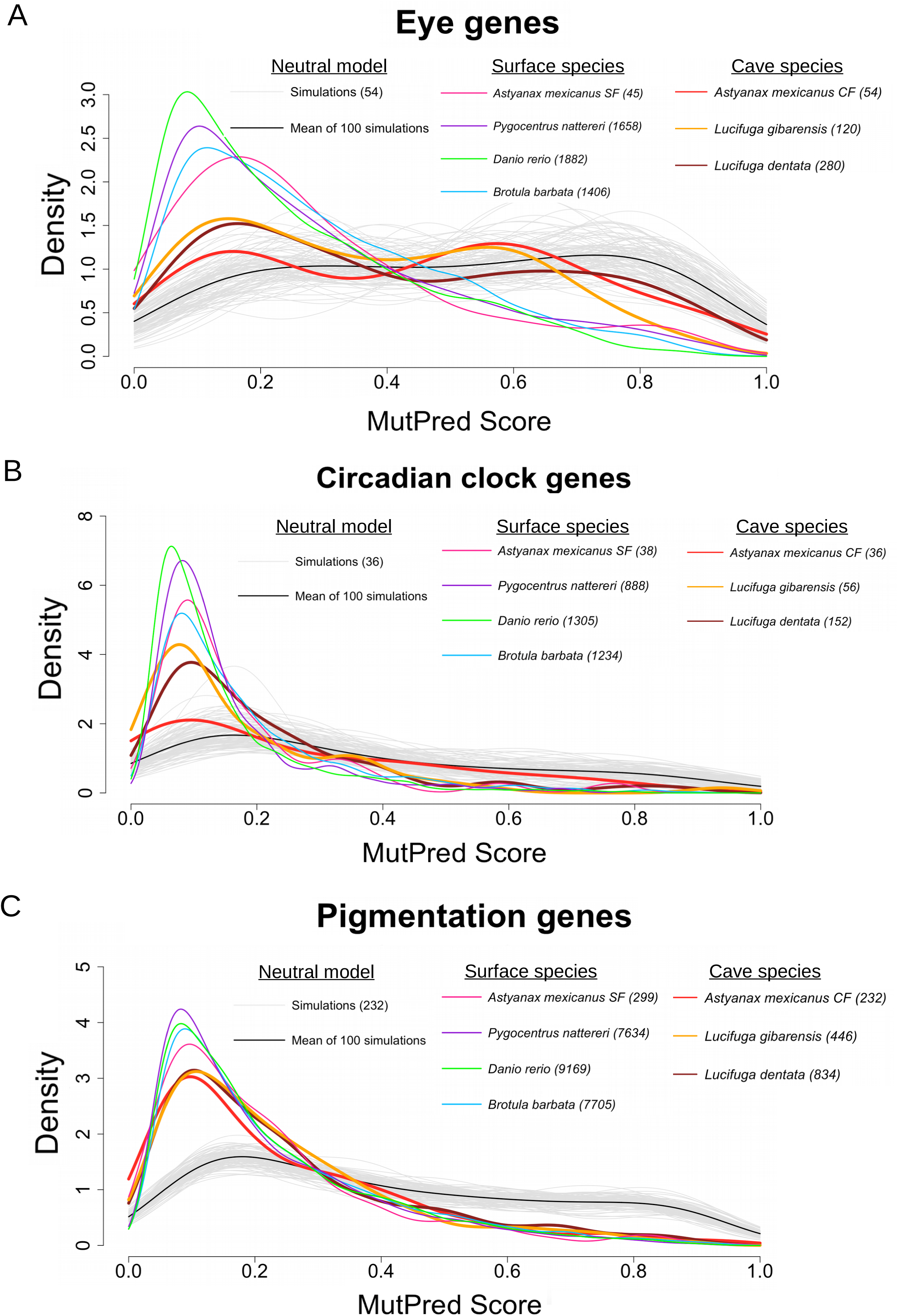
Distributions of MutPred2 scores in several fish lineages and in simulations of substitutions without selection. The number of substitutions in each lineage is given between parenthesis. One hundred simulations were performed with each gene set. In each simulation 54 non-synonymous mutations were generated in eye genes, 36 in circadian clock genes and 232 in pigmentation genes, those numbers corresponding to the numbers of non-synonymous mutations found in *Astyanax mexicanus* cavefish.

<u>With circadian genes</u>, no cavefish ECDF fitted with the expected distribution under neutral evolution (fig. 7B). However, the ECDF of *A. mexicanus* cavefish was different from those of surface fishes and the best fit was obtained with an admixture of 59% of the substitutions from the distribution under selection (**supplementary fig. S20, Supplementary Material** online). For *L. dentata* and *L. gibarensis*, the best fit involved respectively the admixture of 69% and 93% of the substitutions from the distribution under selection (fig. 7B, **supplementary fig. S20, Supplementary Material** online). In accordance with the number of pseudogenes found in *S. rhinocerous* for this set of genes, its ECDF was the closest to the neutral distribution among the three *Sinocyclocheilus* species, with a best fit found with an admixture of 39% of substitutions from the distribution under selection (**supplementary fig. S21** and **fig. S22, Supplementary Material** online).

<u>With pigmentation genes</u>, no cavefish ECDF fitted with the expected distribution under neutral evolution (fig. 7C). All cavefish distributions were very similar to the surface fish distributions, in accordance with the hypothesis that very few genes belonging to this category can be lost, even after cave colonization and/or genome duplication (**see also supplementary fig. S17, fig. S20, fig. S21 and fig. S22, Supplementary Material** online).

In summary, the different approaches consistently suggested that different levels of relaxed selection on the set of eye genes are correlated with the levels of eye degeneration in cavefishes, whereas most circadian clock and pigmentation genes are under strong purifying in these species.

### Dating relaxation of purifying selection on eye genes in *L. dentata*

In order to conciliate results suggesting that most eye genes are dispensable and the finding that selection is not totally relaxed in the *L. dentata* lineage, we postulated two successive periods of evolution, one under selection followed by another under relaxed selection. Three independent approaches were used to estimate when selection was relaxed in this lineage. First, we used the number of eye pseudogenes. With a simple analytical model assuming a LoF mutation rate equal to 0.072 x 10^-8^, the highest probability of finding 19 pseudogenes among 76 neutral genes was obtained for relaxed selection starting 367,779 generations ago (probability > 5% in a range between 273,990 and 480,980 generations) (fig. 8, red curve). Assuming that only 50 eye genes were free to accumulate LoF mutations, this time was pushed back to 611,132 [445,950 – 813,580] generations (fig. 8, pale red curve). Then, simulations were performed in order to take into account variations in the gene length and the number of introns per gene (**supplementary Data_Supp1, Supplementary Material** online), codon usage, transition/transversion ratio (r = 4.57) and effective population size (N_e_) in a range between 100 and 1,000. These simulations and the analytical model gave very similar estimations, the effects of N_e_ and per gene LoF mutation rate variation being marginal (fig. 8, black, green and blue curves, only simulations assuming 76 neutral genes are shown).

**Fig. 8.**
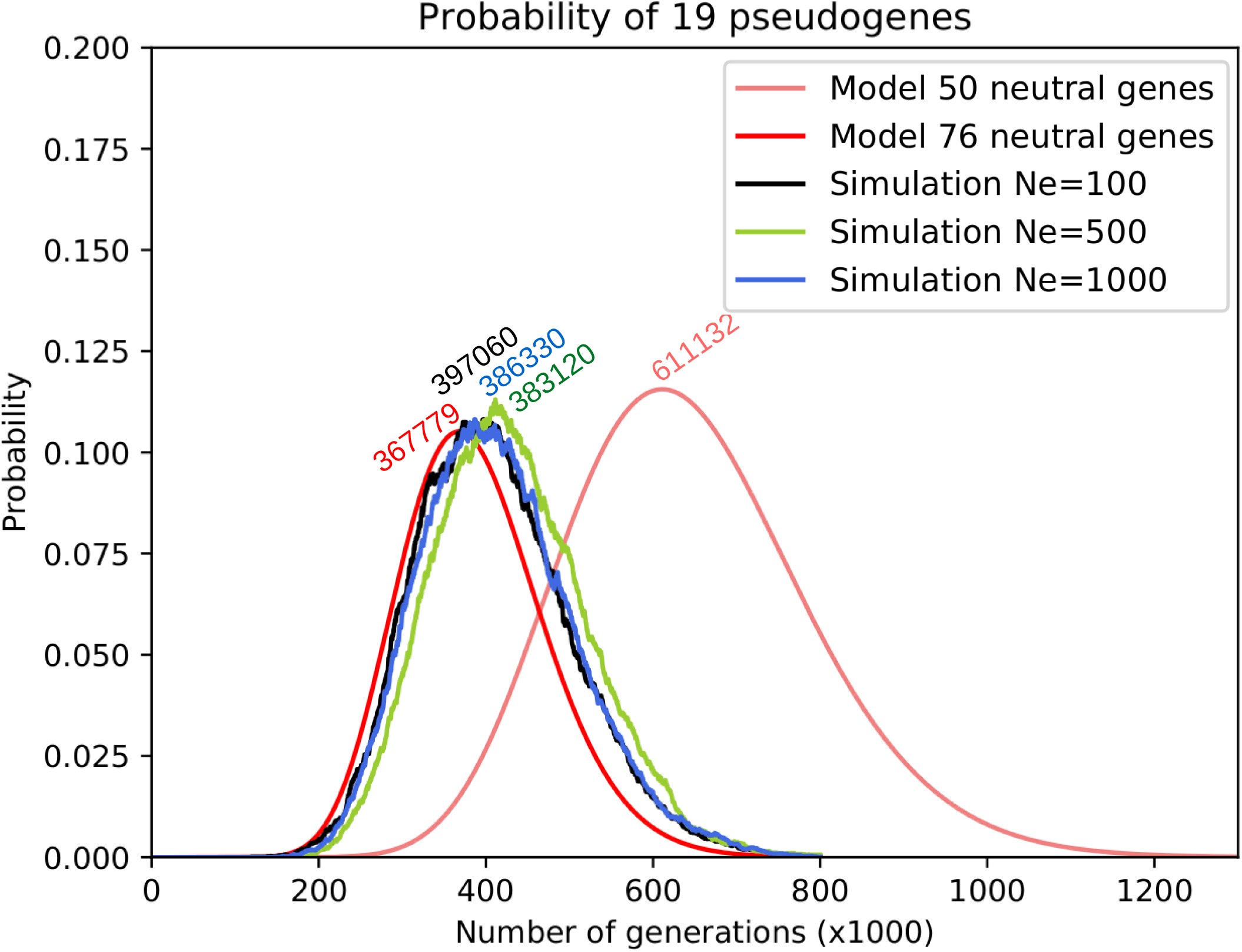
Probability of finding 19 eye pseudogenes in *L. dentata* according to the time of neutral evolution. Red and pink lines: based on an analytical model assuming 76 and 50 neutral genes respectively; other lines: estimations based on 10,000 simulations, assuming 76 neutral genes and taking into account the length and number of introns in each eye genes and considering different effective population sizes. The number of generations for which the highest probability was found is reported above each line.

Second, two dating methods were used (Li, et al. 1981; Meredith, et al. 2009), both based on the hypothesis of a shift of ω from a low value to 1 after purifying selection was relaxed in a lineage. We assumed a divergence time of 80 My between *Lucifuga* and *Brotula* (http://www.timetree.org/). Eye genes of *Lucifuga* species and B*rotula barbata* were individually aligned and alignments concatenated. With one method (Li, et al. 1981), the divergence time between *Lucifuga dentata* and *Lucifuga gibarensis* was estimated equal to 4,110,441 years ago and the time since non-functionalization of eyes genes in *L. dentata* equal to 1,486,042 years. With the other method (Meredith, et al. 2009), ω was estimated equal to 0.271 in the lineage leading to *L. gibarensis* and 0.502 in the lineage leading to *L. dentata*. Assigning these ratios respectively to functional branches and a mixed branch, the time since non-functionalization was estimated to 1,302,485 years.

Third, assuming that in the lineage leading to *L. dentata,* there is an admixture of 66% of the mutations that accumulated under relaxed selection and 34% under selection (**supplementary fig. S20, Supplementary Material** online), and that ω = 0.27 under selection (that is ω estimated in *L. gibarensis*, see **supplementary Data_Supp3, Supplementary Material** online) and ω = 1 under relaxed selection, and the divergence between *L. dentata* and *L. gibarensis* occurred 4,110,441 years ago (estimated above), using the approach described in Materials and Methods, we obtained a congruent estimation of the age of selection relaxation, that is 1,413,991 years ago (table 1, **supplementary Data_Supp3, Supplementary Material** online). Thus, the various methods to date relaxation of purifying selection in *L. dentata* lineage converge to approximately 1.5 My or 400.000 generations, estimations that would be consistent if we assume a generation time of about 4 years in *Lucifuga* cavefishes.

**Table 1.** Estimations of the time without selection on eye genes in *Lucifuga dentata*.

### Distribution of LoF mutations in eye genes of cavefishes *vs* fossorial mammals

An extensive study of the regression of visual protein networks in three fossorial mammals, the Cape golden mole *Chrysochloris asiatica*, the naked mole-rats *Heterocephalus glaber* and the star-nosed mole *Condylura cristata*, has been published (Emerling and Springer 2014). From this publication, we retrieved the number of pseudogenes, their names, and the number of LoF mutations per pseudogene in the three species (fig. 9A). In the Cape golden mole, 18 pseudogenes were found among 63 eye genes, while only 11 pseudogenes were found in the naked-mole rat and 7 in the star-nosed mole. The distributions of LoF mutations per pseudogene in these mammals and those of two blind cavefishes (*L. dentata* and *S. anshuensis*) were compared (fig. 9B). Many independent LoF mutations were found in the same eye genes in fossorial mammals and in cavefishes (fig. 9A). For each species, the distribution of the number of LoF mutations per pseudogene, either taking into account only shared pseudogenes between mammals and fishes or all the pseudogenes, were similar (fig. 9B, main graph and inset respectively). However, the distribution was sharply contrasted between mammals and fishes. In fossorial mammals, most pseudogenes carried many LoF mutations, up to 28 mutations in two pseudogenes of the golden mole and 54 mutations in a single pseudogene of the star-nosed mole (fig. 9, **supplementary Data_Supp1, Supplementary Material** online). On the contrary, in fishes, very few LoF mutations were found in each pseudogene (fig. 9, **fig. S7**, **supplementary Data_Supp1, Supplementary Material** online), the maximum being 5 LoF mutations in one pseudogene of *S. anshuiensis* and 3 LoF mutations in a pseudogene of *L. dentata*. This comparison strongly supports the hypothesis that fossorial mammals have lived in the absence of light for a much longer time than cavefishes, but a smaller subset of genes has been under relaxed selection in mammals.

**Fig. 9.**
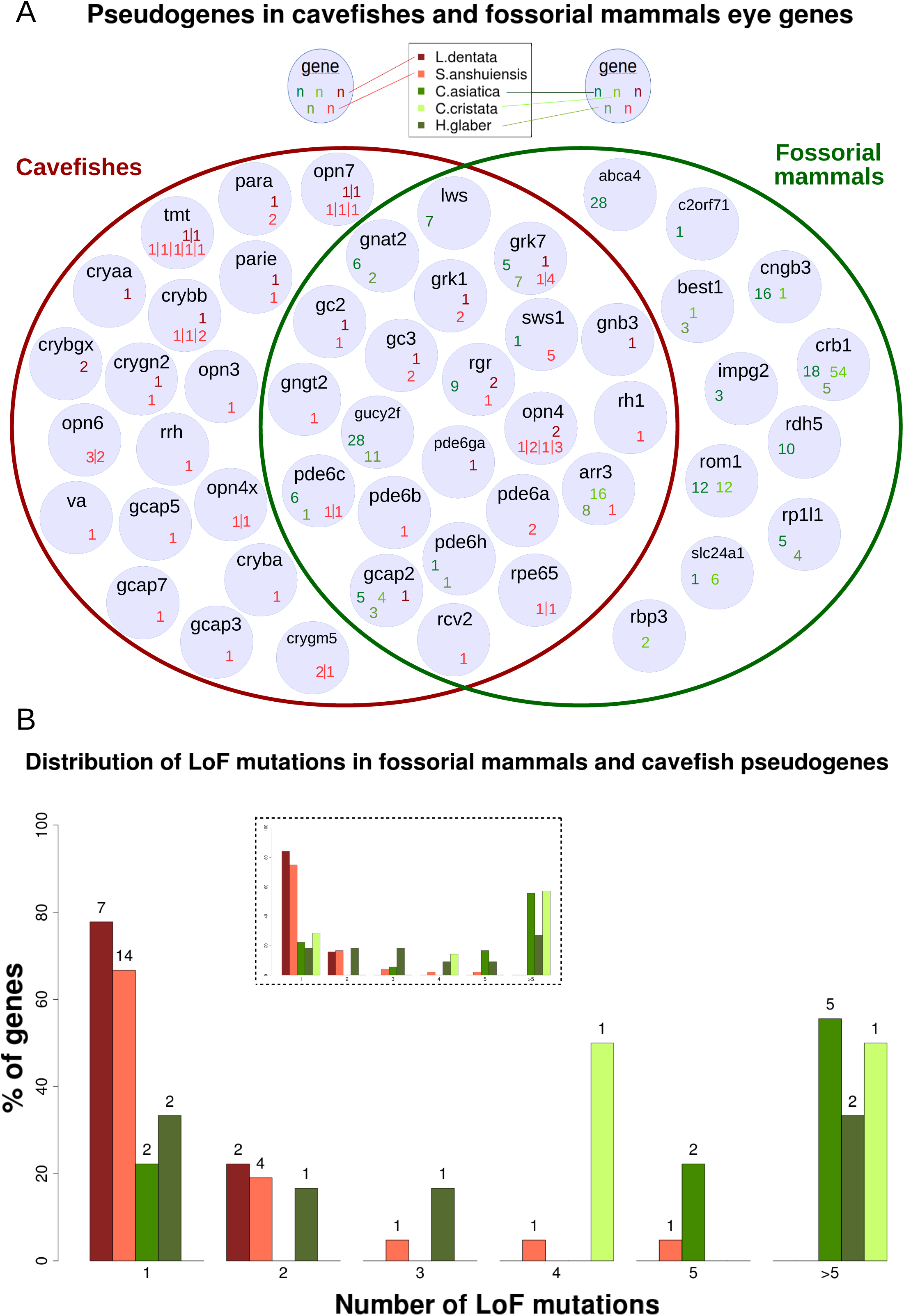
Comparison of eye gene decay in cavefishes *vs* fossorial mammals. (A) Venn diagram showing the genes carrying LoF mutations in both groups. For each gene, the number of LoF mutations found in each species is indicated. (B) Distribution of the number of LoF mutations per pseudogene. The distribution was computed with only the pseudogenes found in both groups or with all pseudogenes (inset). Genes present as one copy in fossorial mammals are often duplicated in *L. dentata* and quadruplicated in *S. anshuiensis,* after one and two WGD respectively. Other gene duplications also sporadically increased the number of paralogs in these fishes. The number of LoF mutations found in these paralogs are separated by vertical lines.

## Discussion

When selection for maintaining a protein in a functional state is relaxed, theory predicts that LoF mutations in its coding and regulatory sequences can reach fixation by random genetic drift (Lynch and Conery 2000; Lahti, et al. 2009). In an isolated population, among a set of dispensable genes, the longer the time of neutral evolution, the higher the expected number of pseudogenes. Eventually, all the genes under relaxed selection will be pseudogenized. At the level of a single gene, the longer the time of neutral evolution, the higher the expected number of LoF mutations. Thus, after a very long period of time of neutral evolution, all the neutrally-evolving genes must carry many LoF mutations. The pace of this gene decay depends essentially on the pace of appearance of LoF mutations (Li and Nei 1977). In the present study, we focussed on a subset of LoF mutations that can be readily detected in genomes, that is mutations generating internal STOP codons, eliminating START or STOP codons, disrupting intron splice sites, and small insertions/deletions (indels) causing translation frameshifts. Although this approach inevitably leads to an underestimation of the number of non-functional genes, it allows comparative studies and molecular dating of selection relaxation in different species. Below, we discuss the patterns of pseudogenization in different sets of genes involved in vision, circadian clock and pigmentation during evolution in the dark of several cavefishes. We show how pseudogenization of eye genes in *Lucifuga dentata* shed new light on gene loss in relation to eye regression in cavefishes. On this basis, we refine previous analyses of other cavefish genomes. At a broader phylogenetic scale, we discuss the contrasted dynamic of pseudogenization in cavefishes and fossorial mammals.

### Putative impact of some LoF mutations

<u>Eye genes</u>: in *L. dentata*, a frameshift was found in the alpha-crystallin, *cryaa*, whose downregulation in *A. mexicanus* cavefish plays a key role in triggering lens apoptosis (Ma, et al. 2014; Hinaux, et al. 2015). Another crystallin, *crybb1*, is pseudogenized in *L. dentata*. Mutations in this gene cause lens opacity in humans (Mackay, et al. 2002). We also found LoF mutations in two opsin receptor kinases, *grk7a* and *grk1b*. Mutations in these proteins can lead to overactive opsin and photoreceptor degeneration (Feng, et al. 2017). These two genes and *grk7b* have similar functions and are all expressed in cones. As these three kinases may have additive effect (Osawa and Weiss 2012), we can hypothesize that absence or malfunction of one of them could be compensated by the others. Such compensation could explain why we found that both *grk7b* ohnologs carry LoF mutations in *S. grahami,* despite this fish has large eyes showing no evidence of degeneration. Another interesting gene is *gnb3b* which is pseudogenized in both *L. dentata* and *L. gibarensis* and which is linked to night-blindness in humans (Vincent, et al. 2016), yet *gnb3^-/-^* mice seem to have functional photoreceptors. Finally, we found LoF mutations in *gcap2*, a guanylate cyclase activator, in both *Lucifuga* species. This gene is associated to *retinitis pigmentosa* in humans (Sato, et al. 2005) but it could be compensated by overexpression of *gcap1* in rods (Makino, et al. 2012). In *Astyanax mexicanus*, a deletion of 11 bp in the phosphodiesterase *pde6b,* a rod-expressed gene, leads to several STOP codons in the catalytic domain (Lagman, et al. 2016). Mutations in this gene were associated with night-blindness and *retinitis pigmentosa* in humans (McLaughlin, et al. 1993; Gal, et al. 1994). Moreover, in mice affected by mutations in the ortholog of *pde6b*, rod photoreceptors degenerate during development resulting in a total absence of photoreceptors in the adult (Farber and Lolley 1974; Chang, et al. 2002). Most LoF mutations were found in the subset of non-visual opsins, which makes their functional impact difficult to evaluate as the functions of these genes are still poorly understood. Two notable exceptions are *opn4m2* and *tmt3a,* pseudogenized in *S. anshuiensis* and *L. gibarensis* respectively, and known to be non-functional and as such involved in the deregulation of the circadian clock in *P. andruzzii*.

<u>Circadian clock genes</u>: in *S. rhinocerous,* both ohnologs of four circadian clock genes, *cry1b*, *cry2a*, p*er2* and *cry-dash,* carried LoF mutations. In *S. anshuiensis*, both ohnologs of *cry-dash* carried also LoF mutations which are independent from those found in *S. rhinocerous*. The gene *cry-dash,* involved in photoreactivation DNA repair, is also pseudogenized in *Phreatichthys andruzzii* (Zhao, et al. 2018) as well as *per2* that could be involved in the disruption of the circadian rhythm in this species (Ceinos, et al. 2018).

<u>Pigmentation genes</u>: both *L. dentata* (depigmented skin) and *L. gibarensis* (pigmented skin) carried independent LoF mutations in *myo7ab*. While no *myo7ab*-/- mutant has been analyzed, the paralog *myo7aa*-/- mutant in zebrafish showed an elevated photoreceptor death but pigmentation was not affected (Wasfy, et al. 2014). Both *Lucifuga* species had independently fixed LoF mutations in *smtla* which is known to increase the number of leucophores at the expense of a reduced number of xanthophores in medaka (Fukamachi, et al. 2009). In *L. dentata, slc2a11b* is pseudogenized and this gene codes for a protein that promotes yellow pigmentation (Kimura, et al. 2014; Parichy and Spiewak 2015). Two other genes, *trpm1a* and *trpm1b* are also pseudogenized in *L. dentata*. During zebrafish development, *trpm1a* is expressed in the retina and melanophores whereas *trpm1b* expression is restricted to the retina (Kastenhuber, et al. 2013). In human, mutations in their ortholog TRPM1 lead to complete congenital stationary night blindness (Audo, et al. 2009). In *L. gibarensis*, *pax7* which promotes xanthophore differentiation (Nord, et al. 2016) carried a LoF mutation as well as *edn3b* that is known to lead to a reduction in iridophore numbers when mutated in zebrafish (Krauss, Frohnhöfer, et al. 2014). In *Astyanax mexicanus*, two pigmentation genes were found with LoF mutations: *mc1r* which carried a 2 bp deletion that could be involved in pigmentation reduction in two cave populations belonging to this species (Gross, et al. 2009) and *tyrp1a* which carried a 1 bp deletion. In zebrafish, morpholino-induced knock-down of *tyrp1a* had no phenotypic effect (Krauss, Geiger-Rudolph, et al. 2014). In *S. rhinocerous* (pigmented skin) and *S. anshuiensis* (depigmented skin) both ohnologs of *gch2* and *pmelb* carried independent LoF mutations. It has been shown that *gch2* mutant lacked proper xanthophore pigmentation at larval stages in zebrafish but no effect were reported in the adult (Parichy, et al. 2000; Pelletier, et al. 2001; Lister 2019). In the same way, injection of *pmelb* morpholinos in the zebrafish had no significant effect on the number of melanosome but led to a significant loss of their cylindrical shape (Burgoyne, et al. 2015). Many pigmentation pseudogenes seem to be compensated by their teleost-specific duplicates when lost in zebrafish, such as *tyrp1a* (Krauss, Geiger-Rudolph, et al. 2014), *pmelb* (Burgoyne, et al. 2015) and *pax7b* (Nord, et al. 2016).

### Contrasted decay of eye genes *vs* circadian clock and pigmentation genes

In order to study pseudogenization in relation to the regression of three traits in cavefishes, we defined three categories, that are eye, circadian clock and pigmentation genes. For most genes, assigning a gene to a category was straightforward, yet for some genes it was more ambiguous. Most eye genes corresponded to a set of genes expressed only in eyes, however fishes also express several non-visual opsins genes that we assigned to this category on the basis of their homology to visual opsins. Genes known for being involved in the circadian clock were assigned to a second set of genes. Noteworthy, some non-visual opsins are involved in this process. Pigmentation genes comprised a large set of genes involved in several processes from pigment cell differentiation to pigment synthesis. Our *a priori* hypothesis was that eye genes should be more prone to degenerate in blind fishes as there are only expressed in eyes or involved in light sensing in other tissues, whereas many circadian clock and pigmentation genes may be maintained as their expression is not restricted to regressed structures and have pleiotropic roles. Indeed, while many pseudogenes were identified among eye genes of some cavefishes, a much smaller proportion of pseudogenes were found among circadian clock and pigmentation genes. In addition, several cases of parallel fixation of LoF mutations in different species among a small subset of genes suggested that only few genes involved in the circadian clock and pigmentation can be lost in cavefishes.

### Molecular evidence of circadian clock disruption in several cavefishes

No LoF mutations were found in the set of circadian clock genes of both *Lucifuga* species. However, *tmt3a*, a non-visual opsin is pseudogenized in *L. gibarensis* and the loss of this gene is involved in the disruption of the circadian rhythm in the cavefish *Phreatichthys andruzzii.* Whereas the survey of LoF mutations did not allow to find evidence of circadian clock loss in *L. dentata*, it is probably the case in *L. gibarensis*. Selection on circadian clock genes is also supported by the analysis of non-synonymous mutations which suggested no higher deleterious mutation accumulation in these species when compared with surface fishes. As expected, no LoF mutations in both ohnologs of circadian clock genes and non-visual ospin genes was found in *S. grahami* which is a surface fish. Unexpectedly, the small-eyed *S. rhinocerous* has accumulated more circadian clock pseudogenes (*per2*, *cry-dash*, *cry1b*, *cry2a*) than the blind *S. anshuiensis* (*cry-dash*), suggesting that the level of eye regression could be loosely correlated with the level of circadian clock disruption. Moreover, as several independent LoF mutations were found in a small number of circadian clock genes, some of them already known to be involved in the circadian clock disruption in other species, it suggests that pseudogenization of a small subset of genes can be involved in this process, in particular those belonging to cryptochromes and period families which are light-inducible genes.

### A small subset of pigmentation pseudogenes

A similar trend was observed among pigmentation genes: independent LoF mutations were found in *myo7ab* and *smtla* of *L. dentata* and *L. gibarensis* and both ohnologs of *gch2* and *pmelb* carried independent LoF mutations in *S. anshuiensis* and *S. rhinocerous*. Recurrent pseudogenization of the same genes suggests that a very small subset of pigmentation genes can be lost, and that these genes might be those which have no or few pleiotropic effects. Indeed, many pigmentation genes code for transcription factors or signaling molecules involved in neural crest-derived, pigment cell differentiation, that are repeatedly used at different times and places during development (Betancur, et al. 2010).

### Many eye pseudogenes in the ancient diploid cavefish *Lucifuga dentata*

Before the present study, there was no evidence on the possibility of pseudogenization of many eye genes in blind cavefishes. In *L. dentata*, we found up to 25% of eye genes carrying LoF mutations. Moreover, the distribution of LoF among genes is consistent with neutral evolution of a large proportion of, if not all, eye genes in this species. On the other hand, in *L. gibarensis* which has small but functional eyes, most eye genes seem under selection but the partial degeneration of the visual system is correlated with the loss of several genes well conserved in eyed fishes. These data allowed us to propose a two-step scenario for the release of selection pressure on eye genes in this genus. The common ancestor of *L. dentata* and *L. gibarensis* was an eyed cavefish that had accumulated a small number of pseudogenes in relation to life in darkness, but none among eye specific genes. In *L. gibarensis*, most eye specific genes have been under purifying selection whereas it has been relaxed in *L. dentata*. Interestingly, the population of *A. mexicanus* from the Pachón cave which is very recent but in which cavefish have highly degenerated eyes, only one eye gene carried a LoF mutation. The lack of correlation between the degree of eye regression and the number of eye pseudogenes underscores the fact that the extent of visual regression should not be taken as a proxy of the evolutionary age of cavefish populations or species.

### Dating blindness in *L. dentata*

Dating changes in selective constraints on traits and genes after cave settlement is a difficult task. Several closely-related methods have been proposed to estimate when a change of selective regime occurred on one gene in one lineage, that is when ω shifted from a value lower than one (a signature of purifying selection) to one (a signature of neutral evolution) (Li, et al. 1981; Miyata and Yasunaga 1981; Meredith, et al. 2009; Zhao, et al. 2010; Wertheim, et al. 2015). With two different methods (Li, et al. 1981; Meredith, et al. 2009), we estimated that the time since selection was released on the eye genes of *Lucifuga dentata* is between 1.3 Mya and 1.5 Mya. Taking into account that 19 pseudogenes were found among 76 eye genes that may be dispensable for a blind fish, and assuming a LoF mutation rate equal to 0.072 x 10^-8^ per site per generation, we estimated the time since *L. dentata* settled in caves about 380,000 generations ago. The generation time is unknown for this fish, and translating the number of generations into years is difficult. However, assuming that the generation time is about four years, which is realistic if we consider that they could reproduce during about ten years, the above independent estimations of relaxed selection would be coherent.

Moreover, using the distribution of the MutPred2 scores, we obtained another and very close estimation (1.4 Ma). Our results suggest that *L. dentata* and *L. gibarensis* could have diverged more than 4 million years ago. The common ancestor of these species could have had well developed eyes that slightly regressed in one lineage (*L. gibarensis*) but much more in the other (*L. dentata*) after a long period without degeneration; or else, the ancestor could have had small eyes like *L. gibarensis* which after a long stasis completely degenerated in the lineage leading to *L. dentata* but remained almost unchanged in the lineage leading to *L. gibarensis*.

Thus, the magnitude of eye degeneration that is often used as a proxy of the age of cave species because it is assumed that eyes degenerate gradually and continually in such environment is likely often misleading. A refined analysis of fish ecology is necessary to better understand the pace and the level of eye degeneration. Indeed, caves are often described as repetitions of the same environment, that is highly isolated and totally dark. However, some cavefishes such as *Lucifuga gibarensis* and closely-related small eyed species can be found in caves that are partially lighted, or sink holes in the sea. Such a complex environment could be the reason for the maintenance of small yet functional eyes in these species, like in fossorial mammals.

### Pattern of LoF mutations in recent tetraploids with different level of troglomorphy: the case of *Sinocyclocheilus*

The genus *Sinocyclocheilus*, which is endemic to southwestern karst areas in China, is the largest cavefish genus known to date (Xiao, et al. 2005). LoF mutations were found in several genes of three species, one species (*S. anshuinensis*) being blind and depigmented, another species (*S. rhinocerous*) having small eyes and being pigmented, and the last one (*S. grahami*) showing no such troglomorphic traits (Yang, et al. 2016). These species share a WGD with other cyprinids such as the common carp *Cyprinus carpio* (David, et al. 2003; Yuan, et al. 2010) which could explain why even the surface fish carry many LoF mutations in eye, circadian clock and pigmentation genes (Yang, et al. 2016). However, no thorough comparisons were performed. Our results are consistent with a rapid radiation within this genus (Xiao, et al. 2005) as only few LoF mutations were found in internal branches of their phylogenetic tree. The divergence between *Cyprinus carpio* and the *Sinocyclocheilus* species may have occurred soon after the WGD as only two shared LoF mutations were found. The number of eye pseudogenes in the blind *S. anshuiensis* is much higher than in the small-eyed *S. rhinocerous* and the eyed *S. grahami*, a result supporting the cumulative effect of tetraploidy and cave settlement on the rate of accumulation of LoF mutations. As most genes are present twice, a gene function is lost if, and only if, at least one LoF mutation is present in each ohnolog. With this criterion, seven genes were lost in *S. anshuiensis*, but only one gene in *S. rhinocerous* and *S. grahami*. Selective pressure was relaxed on one copy of these genes after the WGD, but a complete relaxation occurred only after cave settlement in *S. anshuiensis*. Among the genes for which both ohnologs are mutated in *S. anshuiensis*, the mutations in *pde6c* could have a role in photoreceptors degeneration, as suggested by a study of zebrafish mutants (Stearns, et al. 2007). *Sinocyclocheilus rhinocerous* lost the two functional copies of *gcap1* and it has been shown that two missense mutations in this gene lead to significant disruptions in photoreceptors and retinal pigment epithelium, together with atrophies of retinal vessels and choriocapillaris in zebrafish (Chen, et al. 2017). However, knockout of *gcap1* in mice showed that its absence does not change expression level of other phototransduction proteins thanks to a compensation by *gcap2*. Nevertheless, the knock-down leads to a delayed recovery after light exposure (Makino, et al. 2012).

Analyses with RELAX and the estimation of the admixture of MutPred2 score distributions that best fit with the observed score distribution also suggest that purifying selection on eye genes is much higher in *S. grahami* than in *S. anshuiensis* and *S. rhinocerous*, much lower on circadian genes of *S. rhinocerous* but high on pigmentation genes in these three species.

These results are congruent with the level of pseudogenization observed for the three gene sets in the three species.

### Very few pseudogenes in the recent settler *Astyanax mexicanus*

In the reference genome of *Astyanax mexicanus* cavefish, a LoF has been found in the eye gene *pde6b*. This mutation went unnoticed in previous studies but may well contribute to retinal degeneration. No LoF were found in clock genes. Among pigmentation genes, a 1 bp deletion was found in *tyrp1a* and a 2 bp deletion in *mc1r*. The latter mutation has been associated with the brown phenotype of some populations (Gross, et al. 2009) but the finding of a close and functional tandem duplicate suggest that it actually may not be the cause of this phenotype (Gross, et al. 2017). Overall, these results are in accordance with a very recent settlement of *Astyanax* cavefish (Fumey, et al. 2018) that did not allow the fixation of many eye pseudogenes despite the lack of purifying selection on most, if not all, eye genes. The extreme eye degeneration with only one LoF in *Astyanax* cavefish eye genes further questions the nature of the developmental mechanisms involved in eye loss in this species, the pace of eye degeneration and the correlation of eye degeneration with gene decay.

### Contrasted dynamics of pseudogenization in fossorial mammals and cavefishes

The genomes of three independently-evolved fossorial mammals have previously allowed an extensive study of LoF mutations in genes coding for proteins involved in retinal networks (Emerling and Springer 2014). These animals have functional eyes, but star-nosed moles often leave their burrows and have the greatest exposure to light whereas naked mole-rats and Cape golden moles are entirely subterranean. In addition, the eyes of Cape golden moles are subcutaneous. More pseudogenes were found in the Cape golden mole than in the naked-rat genome and the lowest number of pseudogenes was found in the star-nosed mole genome, suggesting that the decrease in retinal exposure to light allowed the decay of more eye genes. The most striking difference between cavefishes and fossorial mammals is that pseudogenes of cavefishes accumulated only one or a couple of LoF mutations per pseudogene whereas some pseudogenes of fossorial mammals carried a large number of LoF mutations. This difference in molecular decay strongly suggests that the fossorial mammals adapted to the subterranean environment a long time ago whereas colonisation of the dark environment is much more recent in the case of the cavefishes.

## Conclusion

Our analyses suggest that blind cavefishes examined so far are not very ancient. They all lost their eyes during the Pleistocene, the oldest during early Pleistocene and the most recent during the late Pleistocene or even later in the Holocene. The sequencing of a large number of blind cavefish genomes will be necessary to identify the whole set of eye genes that are dispensable in the dark, when eyes are highly degenerated. Moreover, finding a blind cavefish genome in which most eye genes are pseudogenized and carry many LoF mutations would refute our current working hypothesis that blind cavefishes cannot thrive for a very long time in cave ecosystems.

## Materials and Methods

### Assembly of L. dentata and L. gibarensis draft genomes

The sequenced *L. dentata* specimen was a female, blind and depigmented. All the fish belonging to this species are blind whereas their pigmentation is highly variable (Garcia-Machado, unpublished data). The sequenced *L. gibarensis* specimen was a male, had small eyes and was pigmented. All the fish belonging to this species have small eyes whereas their pigmentation is also highly variable (García-Machado, et al. 2011). DNA was extracted using a protocol already described elsewhere (García-Machado, et al. 2011). For *L. dentata*, paired-end libraries were prepared with different insert sizes: 200 bp, 400 bp and 750 bp. A mate-pair library was also prepared with insert size in the range 3-5 kb. For *L. gibarensis*, only one mate-pair library was prepared, which had inserts size between 3 kb and 10 kb. *Lucifuga dentata* libraries were sequenced on an Illumina HiSeq 2000 sequencer whereas *L. gibarensis* library was sequenced on an Illumina NextSeq sequencer. After cleaning steps (adaptors trimming and quality trimming), *L. dentata* assembly of a draft genome was performed using Minia (Chikhi and Rizk 2013) on all data, resulting in 662,154 contigs. After assembling, and as Minia doesn’t use the paired-end information, scaffolding steps were performed using SSPACE (Boetzer, et al. 2011) on one library at a time in ascending order of insert size. The number of scaffolds decreased from 662,154 to 161,599 with the first library (insert size of 200 bp), to finish with 48,241 scaffolds with the mate pair library. This result was corrected by REAPR (Hunt, et al. 2013) to obtain 52,944 scaffolds. The remaining gaps were filled by GapCloser (Luo, et al. 2012).

The quality and completeness of the draft genome of *L. dentata* were assessed by remapping paired-end reads to the assembly using BWA v0.7.11 (Li and Durbin 2009) and BUSCO (Kriventseva, et al. 2015) with the Actinopterygii dataset comprising a total of 4,584 conserved genes. The latter analysis was performed also on published draft genomes of three other Ophidiiformes (*Brotula barbata*, *Carapus acus* and *Lamprogrammus exutus*). Sequences from *L. gibarensis* were mapped on the genome of *L. dentata* using BWA v0.7.11.

### Assembly of *Lucifuga dentata* transcriptome

Gonads, gills, heart and brain were dissected and stored in RNA-Later (Ambion). Total RNA isolation (using Trizol) lead to yields of 870 ng/μl in gonads, 750 ng/μl in gills, 240 ng/μl in heart, 390 ng/μl in brain. ARN from gonads, gills and heart were mixed in equal proportions to construct the first library. ARN from the brain was used to construct the second library. For library preparation, polyA + RNA were extracted, fragmented, and directional libraries were prepared using the Small RNA Sample Prep Kit (Illumina). Both libraries were sequenced on an Illumina NextSeq500, on a Paired-end 2×150 bp run, using the High Output After cleaning steps (adaptors trimming and quality trimming), a *de novo* transcriptome assembly was obtained using Trinity and a quality assessment was realized following Trinity recommendations (https://github.com/trinityrnaseq/trinityrnaseq/wiki/Transcriptome-Assembly-Quality-Assessment).

### Annotation of *Lucifuga dentata* draft genome

First, repetitive elements were identified using RepeatMasker v4.0.7 (Smit, et al. 2013), Dust (Morgulis, et al. 2006) and TRF v4.09 (Benson 1999). A species specific *de novo* repeat library was built with RepeatModeler v1.0.11 (Smit and Hubley 2008) and repeated regions were located using RepeatMasker with the *de novo* and *Danio rerio* libraries. Bedtools v2.26.0 (Quinlan and Hall 2010) were used to merge repeated regions identified with the three tools and to soft masked the genome. Then, MAKER3 genome annotation pipeline v3.01.02-beta (Holt and Yandell 2011) combined annotations and evidence from three approaches: similarity with fish proteins, assembled transcripts and *de novo* gene predictions. Protein sequences from 11 other fish species (*Astyanax mexicanus*, *Danio rerio*, *Gadus morhua*, *Gasterosteus aculeatus*, *Lepisosteus oculatus*, *Oreochromis niloticus*, *Oryzias latipes*, *Poecilia formosa*, *Takifugu rubripes*, *Dichotomyctere nigroviridis*, *Xiphophorus maculatus*) found in Ensembl were aligned to the masked genome using Exonerate v2.4 (Slater and Birney 2005). RNA-Seq reads were mapped to the genome assembly using STAR v2.5.1b (Dobin, et al. 2013) with outWigType and outWigStrand options to output signal wiggle files. Cufflinks v2.2.1 (Trapnell, et al. 2010) was used to assemble the transcripts which were used as RNA-seq evidence. A *de novo* gene model was built using Braker v2.0.4 (Hoff, et al. 2016) with wiggle files provided by STAR as hints file for GeneMark and Augustus trainings. The best supported transcript for each gene was chosen using the quality metric called Annotation Edit Distance (AED) (Eilbeck, et al. 2009). The annotation completeness of coding genes was assessed by BUSCO using the Actinopterygii gene set. Homology to uniprot database was used to infer functions of predicted genes with Blastp and an e-value cutoff of 1e^-6^. Interproscan 5.35 (Jones, et al. 2014) was used to detect proteins with known functional domains.

### Analysis of repeated elements

The *de novo* library of repeated elements was refined with the following procedure: removal of short (<80 bp) consensus repeats; reannotation of satellite sequences as well as of putative DNA or LTR transposable elements (TEs) by aligning each consensus against itself (this procedure allows to visualize internal repeats); Blastn (Altschul, et al. 1990) of the library against itself and removal of redundant TEs; Blastx (Altschul, et al. 1990) of « Unknown » repeats against the NCBI protein database and removal of multigene families erroneously identified as putative TEs; reannotation of putative SINEs according to the SINE-scan program (Mao and Wang 2017). Finally, the library was manually curated: consensus sequences were compared to an in-house library of transposable element proteins using BlastX. Matching Unknown elements were renamed according to their hits against this library. Consensus sequences showing incongruent annotations between RepeatModeler automatic classification and our manual annotation were further submitted to Censor (Kohany, et al. 2006). This TE library was used as repeat database for a RepeatMasker search in the genome (Smit, et al. 2013). Overlaps in RepeatMasker output were discarded by selecting highest scoring elements. Repeat fragments closer than 20 bp and having the same name were merged. The Landscape was reconstructed from RepeatMasker align output using the calcDivergenceFromAlign.pl and createRepeatLandscape.pl utilities of the RepeatMasker suite.

### Identification of eye, circadian and pigmentation genes

The set of eye genes included all opsins, visual opsins that are expressed in eye photoreceptor cells (cone and rods) but also non-visual opsins that are expressed in a wide variety of tissues. It also comprised eye specific crystallin genes. Crystallin genes code for several families of proteins that are implicated in the transparency of the lens and fine tuning its refraction index, but can also have other functions, not well known for many of them (Thanos, et al. 2014). Expression patterns in zebrafish reported in ZFIN database (https://zfin.org/) and in *A. mexicanus* (Hinaux, et al. 2015) were used to identify and select a subset of eye specific crystallins. Noteworthy, *crygm2* paralogs were excluded from the analysis because many copies (more than 50 copies in *A. mexicanus*) were found as in other fish genomes most likely allowing relaxed selection on some copies independently to relaxed selection due to environmental shift. The set of eye genes also included genes coding for proteins involved in the phototransduction cascade: RPE65, Arrestins, Recoverins, Transducins, PDE6, CNGA3 and CNGB3, GCAPs, zGCs, and GRKs. These genes code for a highly heterogeneous set of proteins with regard to their structure and functions (Imanishi, et al. 2002; Wada, et al. 2006; Schonthaler, et al. 2007; Matveev, et al. 2008; Nishiwaki, et al. 2008; Rätscho, et al. 2009; Renninger, et al. 2011; Fries, et al. 2013; Lagman, et al. 2015; Zang, et al. 2015; Lagman, et al. 2016). Only genes whose expression was restricted to the retina and/or the pineal complex were retained. Sets of circadian clock and pigmentation genes were defined on the basis of gene lists established in previous studies (Li, et al. 2013; Lorin, et al. 2018). The set of circadian genes was completed with *ck1da* and *ck1db* genes which are specific kinases of *cry* and *per* genes (Takahashi, et al. 2008) and *aanat1* and *aanat2* genes whose expression are regulated by the circadian clock in zebrafish (Vatine, et al. 2011). The sequences of visual and non-visual opsins of zebrafish were retrieved from (Davies, et al. 2015). Other eye genes, circadian and pigmentation genes of zebrafish were retrieved from GenBank. Series of blastn and tblastx (Altschul, et al. 1990) with zebrafish sequences were performed against *A. mexicanus* surface and Pachón cave genomes (GCF_000372685.2 and GCF_000372685.1 respectively), *S. grahami*, *S. rhinocerous*, *S. anshuiensis*, *P. nattereri*, *B. barbata*, *C. acus* and *L. exutus* genomes (GCF_001515645.1, GCF_001515625.1, GCF_001515605.1, GCF_001682695.1, GCA_900303265.1, GCA_900312935.1 and GCA_900312555.1 respectively), and *L. dentata* and *L. gibarensis* genomes (this study). Matching regions were extracted using samtools (Li 2011) and coding DNA sequences (CDS) were predicted using Exonerate with protein sequences of zebrafish (Slater and Birney 2005).

### Phylogenetic analyses

Orthologous and paralogous relationships between genes were inferred through phylogenetic analyses. First, coding sequences were aligned using MUSCLE (Edgar 2004), after having taken into account indels (*i.e.* adding N where nucleotides were missing or removing additional nucleotides). For each alignment, DNA sequences were translated into protein sequences and a maximum likelihood phylogenetic tree was inferred using IQ-TREE (Nguyen, et al. 2015) with the optimal model found by ModelFinder (Kalyaanamoorthy, et al. 2017) and the robustness of the nodes was evaluated with 1,000 ultrafast bootstraps (Hoang, et al. 2018). The trees were rooted and visualized using iTOL (Letunic and Bork 2006). Phylogenetic trees and IQ-TREE files can be found in **supplementary folder phylogenies, Supplementary Material** online.

### Identification of LoF mutations

We classified CDS in three classes: 1) complete, 2) pseudogene (characterized by the presence of at least one among the following mutations: an internal STOP codon, an indel leading to a frameshift, the loss of the initiation codon, the loss of the STOP codon, a mutation in a splice site of an intron), 3) incomplete. Incomplete genes can be artifacts of different origins such as missing data, assembly errors (Florea, et al. 2011) and gene prediction errors due to sequence divergence. Nonetheless, they can be real, resulting from large genomic deletions. In the case of the *A. mexicanus* cavefish genome, using PCR, we could check that about 85% of the incomplete genes were assembly errors (data not shown) and they were not further analyzed. Given the low quality of the *A. mexicanus* cavefish genome assembly compared to the surface one and in order to get good gene sequences, cave reads were retrieved and mapped onto the surface genome using the NCBI remapping service. This approach allowed the identification of an opsin gene repertoire (36 genes) slightly larger than the one recently published (33 genes) using only the cavefish genome (Simon, et al. 2019). Similarly, *Lucifuga gibarensis* reads were mapped on the *Lucifuga dentata* genome. Orthologous genes from a cod (*Gadus morhua*), a medaka (*Oryzias latipes*), a platyfish (*Xiphophorus maculatus*), a stickleback (*Gasterosteus aculeatus*), a pufferfish (*Dichotomyctere nigroviridis*), a tilapia (*Oreochromis niloticus*) and a spotted gar (*Lepisosteus oculatus*) were downloaded from Ensembl (Ensembl IDs can be found in **supplementary Data_SuppS2, Supplementary Material** online). For these fishes, visual opsin sequences were retrieved from an extensive study at the scale of ray-finned fishes (Lin, et al. 2017).

### Testing randomness of LoF mutation locations along the genes

In order to evaluate whether LoF mutations were randomly distributed or clustered along the genes, we used a method initially designed for estimating the randomness of intron insertions (Lynch and Kewalramani 2003). We computed the effective number of gene segments defined by: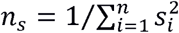, with *n* being the number of segments of genes separated by *n*-1 LoF mutations and *s_i_* being the length of the *i*th segment. As LoF mutations are found in several genes with different lengths, the position of each LoF mutation was normalized by dividing by the length of the coding sequence, the sum of *s_i_* was thus equal to 1 for each gene. The most extreme case of LoF dispersion is the one in which all segments are of the same length (1/*n*), *i.e.* the LoF mutation are regularly spaced out, yielding *n_s_* = *n*. On the other hand, if all LoF are clustered at one end of the genes, one segment approaches length 1.0, while all others approach 0.0, yielding *n_s_* = 1. In order to obtain the distribution of the values of *n_s_* under the null model of fixation of LoF at random positions, 100,000 simulations of random distribution of the observed number of LoF mutations along a gene of length 1.0 were performed.

### Estimation of the number of eye genes under relaxed selection in *Lucifuga* spp. using the distribution of LoF mutations per gene

In order to estimate the number genes under relaxed selection (*V*) in a sample of *a priori* useless eye genes (*T*) in *L. dentata* and *L gibarensis*, we compared the observed distribution of LoF mutations per eye gene with the expected distribution, taking into account that only a fraction (*V*) of these genes are under relaxed selection and can accumulate LoF mutations and that *T* - *V* genes are under selection and cannot carry LoF mutations. Assuming that a LoF mutation has a probability 1/*V* to appear in a gene among *V* genes under relaxed selection, the probability that a gene contains *X* LoF mutations can be computed as follows:

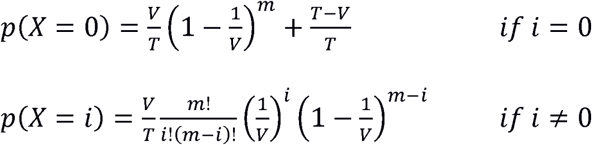

where *m* is the total number of LoF mutations.

In order to take into account that eye genes do not have the same length and the same number of introns and thus mutations do not have the same probability of occurring in each gene (they are more likely in a gene with several large exons and several introns than in a gene with only one short exon), we ran 10,000 simulations of the distribution of *m* mutations in a random sample of *V* genes taken at random among *T* eye genes, and taking into account the length and the number of introns in each gene to estimate its relative mutation rate. The distributions of the number of LoF mutations per gene in *L. dentata* and *L. gibarensis* were compared with expected distributions obtained with the two methods described above and for different values of *V*.

### Sequence divergence and evidence of relaxed selection in cavefishes

For diploid species, genes belonging to the same gene set (eye, circadian clock or pigmentation) were concatenated. In order to analyze the genes of the tetraploid *Sinocyclocheilus* species, another alignment was produced in which each ohnolog of a given gene was concatenated with one ohnolog taken at random of the other genes, leading to two sets of concatenated genes for each species. With both alignments of concatenated sequences, maximum likelihood estimates of ω were obtained using the program codeml from the PAML package (Yang 2007) with a free-ratio model allowing a different ratio for each branch (**supplementary fig. S13, Supplementary Material** online).

Another approach used for detecting relaxed selection was based on analyses with the program RELAX (Wertheim, et al. 2015), assigning surface fishes as reference and excluding the small eyed fish *Lucifuga gibarensis*, the eyeless fishes *Lucifuga dentata* and *Astyanax mexicanus* CF. Each cavefish was independently assigned as the test branch. The value of the parameter k which is <1 if selection is relaxed and >1 if selection is intensified was considered as evidence of a change in the selective regime (**supplementary fig. S14**, **fig. S15** and **fig. S16, Supplementary Material** online).

### Inferring the deleterious impact of amino acid variants with MutPred2

Maximum likelihood inference of amino acids substitutions were performed using the program aaml from the PAML package (Yang 2007). For each amino acid substitution, MutPred2 scores (Pejaver, et al. 2017) and Grantham’s distances (Grantham 1974) were computed to estimate the deleterious impact of the substitutions.

In order to compare the distribution of scores (or distances) for a set of genes and along a branch with the distribution expected under relaxed selection, simulations of random substitutions were generated in these genes, taking into account the length of the coding sequence of each gene and the transition/transversion ratio (https://github.com/MaximePolicarpo/Molecular-decay-of-light-processing-genes-in-cavefishes/blob/master/Neutral_evolution_for_mutpred.py). MutPred2 output files can be found **supplementary folder MutPred2_results, Supplementary Material** online).

### Dating relaxation of selection with the number of eye pseudogenes in *L. dentata*

In absence of selection, the probability of fixation of a LoF mutation, initially absent in a population, is:

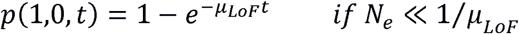

where *μ_LoF_* is the LoF mutation rate, N_e_ is the effective population size and *t* is the number of generations (Li and Nei 1977).

Thus, if *μ_LoF_* is identical for a set of genes, the probability that D among T genes have fixed a LoF after time *t* is:

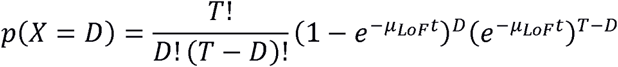

The derivative of this function with respect to *t* allows to find for which value of *t* the probability *p*(*X* = *D*) is maximal:

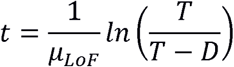

For a given set of genes, the rate of LoF mutation was computed as follows:

*i)* The genetic code implies that among 549 (61 x 9) mutations in sense codons, 23 lead to a STOP codon, that is ∼ 4% if the frequency of each codon is 1/61 and transitions are as frequent as transversions. As among those 23 mutations, 5 are transitions and 18 are transversions, the transition/transversion ratio (*r*) can be taken into account to estimate more accurately the fraction of mutation leading to a STOP codon *f* = (5r + 18)/(183r + 366). Using a R script, the estimation of *f* was further refined by taking into account codon frequencies (frequency_new_stop.py). For eye genes, taking into account estimations of r in *Lucifuga* spp. and *Sinocyclocheilus* spp. (4.57 and 1.95 respectively) and the codon frequencies of their eye gene sequences, *f* was estimated equal to 0.031 and 0.037 respectively in these groups of species. Moreover, taking into account that 13 internal STOP codons were found in *Lucifuga* spp. and 47 in *Sinocyclocheilus* spp., we estimated a weighted mean *f* = 0.036 for the whole eye gene dataset. Applying the same approach, we found *f* = 0.038 for the circadian clock genes and *f* = 0.036 for pigmentation genes (Details in **Data_Supp1, Supplementary Material** online). For the three datasets taken together, weighting by the length of the concatenated genes in each dataset, we estimated a global mean *f* = 0.036. For a set of coding sequences of length *l* (sum of the CDS lengths), the rate of mutation to a STOP codon *μ_stop_* = *fμl*, where *μ* is the nucleotide mutation rate / site.

*ii)* The rate of indels leading to frameshifts (*i.e.* indel length modulo 3 ≠ 0) relative to the rate of new STOP codons is 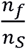 where *n_f_* and *n_S_* are the numbers of indels leading to frameshifts and new STOP codons respectively. The rate of frameshifts and new STOP codons respectively. The rate of frameshifts is 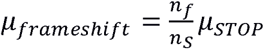.

*iii)* The rate of splice site mutations is 4*n_i_μ* (where *n_i_* is the number of introns in the set of genes).

*iv)* The rate of START codon loss is 3*n_g_μ* (where *n_g_* is the number of genes).

*v)* The rate of STOP codon loss is 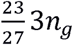 (where 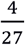 is the proportion of mutations in a STOP codon which leads to another STOP codon).

Globally, for the set of genes, the LoF mutation rate is

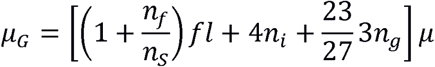

if all genes have the same CDS length and the same number of introns.

The rate of LoF mutations per gene is 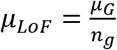

In order to assess the effect of the high variability of gene length and intron number observed in eye genes on pseudogene accumulation through time, a program was written to simulate decay of this set of genes through accumulation of STOP codons, frameshifts, splice site mutations, initiation and STOP codon losses, taking into account the length and the number of introns in each gene. At each generation and for each gene, the probability that a new LoF appears in one ancestral and functional allele at frequency *q* in a population of size *N_e_* is: 2N_e_q*μ_LOF_*. When a new LoF mutation appears its frequency is 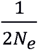 and the total frequency of LoF mutations is *p* + 1/2*N_e_*, where *p*=1 - *q*. We assumed random mating, no selection and no migration and a constant population size. Genetic drift between two generations was simulated taking into account the new allele frequencies if a mutation occurred, and 2N_e_ (the number of alleles sampled to generate the next generation).

The simulation program was written in Python (https://github.com/MaximePolicarpo/Molecular-decay-of-light-processing-genes-in-cavefishes/blob/master/SimulationScript.py).

### Other methods for dating selection relaxation on eye genes of *L. dentata*

Eye genes of the two Cuban cave brotulas (*L. dentata* and *L. gibarensis*) and an outgroup (*Brotula barbata*) were concatenated and aligned. We supposed that eye genes have been under selection along the branches of the phylogenetic tree, except in the lineage leading to *L. dentata* which is a mixed branch (with a period of time under selection followed by a period of time under relaxed selection). The time since selection was relaxed was estimated using two slightly different methods both relying on a shift of the nonsynonymous substitution rate after relaxed selection (Li, et al. 1981; Meredith, et al. 2009). The time of divergence between *Brotula barbata* and Cuban cave brotulas was set to 80 Mya (http://www.timetree.org/).

As an alternative approach, we used the distribution of MutPred2 scores in the lineage leading to *L. dentata*. First we computed the proportions of two distributions, one under selection as in the zebrafish lineage (*p_s_*) and one without selection as in simulated data (*p_n_*), that produce a mixture distribution that best fit the distribution of MutPred2 scores in the lineage leading to *L. dentata*. We assumed that *ω_s_* under selection shifted to *ω_n_* when selection is relaxed. We called *T_d_* the period of time since the separation of *L. dentata* and *L gibarensis*, *t_s_* the period of time of evolution under selection and *t_n_* the period of time under relaxed selection in the lineage leading to *L. dentata*. In this lineage, the proportion of nonsynonymous substitutions that accumulate under selection depends on *ω_s_* and *t_s_* and the proportion of nonsynonymous substitutions that accumulate under relaxed selection depends *ω_n_* and *t_n_*. Thus 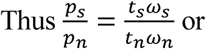 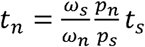

### Comparison of eye gene decay in cavefishes and fossorial mammals

In order to compare the decay of eye genes in cavefishes and fossorial mammals, the number of pseudogenes and the number of LoF mutations per pseudogene among genes coding for proteins involved in retinal networks in three fossorial mammals (Cape golden mole *Chrysochloris asiatica*, naked mole-rat *Heterocephalus glaber* and star-nosed moles *Condylura cristata*) were retrieved from a publication (Emerling and Springer 2014).

## Data Availability

*Lucifuga dentata* Whole Genome Shotgun project has been deposited at DDBJ/ENA/GenBank under the accession VXCM00000000. The version described in this paper is version VXCM01000000

*Lucifuga dentata* Transcriptome Shotgun Assembly project has been deposited at DDBJ/EMBL/GenBank under the accession GIAU00000000. The version described in this paper is the first version, GIAU01000000.

*Lucifuga gibarensis* raw sequences were submitted to the SRA Bioproject: PRJNA610231 The original GFF3 annotation file of *Lucifuga dentata* and scaffolds smaller than 200 bp are available in Supplementary files.

Python programs and R scripts used in this paper can be found in: https://github.com/MaximePolicarpo/Molecular-decay-of-light-processing-genes-in-cavefishes.

## Supplementary Material

Supplementary data are available at Molecular Biology and Evolution.

## Acknowledgments

This work was supported by a collaborative grant from Agence Nationale de la Recherche (BLINDTEST to S.R. and D.C.) and from Institut Diversité Ecologie et Evolution du Vivant (to S.R. and D.C). We thank Yan Jaszczyszyn, Jean Mainguy, Nina Paffoni and Isabelle Germon for their help in sequencing and analyzing the genomes of *L. dentata* and *L. gibarensis*. We also thank Carlsbergfondet for financial support (grant no. 2013_01_0501) for sampling *Lucifuga gibarensis*.

## Ethics approval

Animals were treated according to the French and European regulations for handling of animals in research.

## Sampling authorization

*Lucifuga dentata*: a permit [LH 112 AN (135) 2013] was provided to the Centro de Investigaciones Marinas, University of Havana by the Cuban authorities in December 2013 to study cave species diversity including nematodes, crustaceans and fishes. As the species was listed Vulnerable (VU) by the IUCN, only two adult individuals (MFP 18.000278) were sampled (12 January 2014) from one of its largest and demographically stable populations (Emilio Cave, Las Cañas, Artemisa Province, Cuba).

*Lucifuga gibarensis*: a permit [PE 2014/82] was provided to the Centro de Investigaciones Marinas, University of Havana by the Cuban authorities in November 2014 to study cave species diversity including nematodes, crustaceans and fishes. A single adult fish (MFP 18.000279) was sampled (20 November 2014) from the Macigo Cave (Aguada de Macigo del Jobal), Gibara, Holguín Province, Cuba.

---

**Fig. S1.** Photos of the specimens used for genome sequencing. (A) *Lucifuga dentate*. (B) *Lucifuga gibarensis*.

**Fig. S2.** *L. dentata* scaffold size distribution. Based on 3,537 scaffolds longer than 50kb (49,407 scaffolds < 50 kb not used).

**Fig. S3.** BUSCO analyses using the Actinopterygii gene database (v3.1.0). We assessed the completeness of three published Ophidiiformes genomes (*Brotula barbata*, *Carapus acus* and *Lamprogrammus exutus*), *Lucifuga dentata* genome, gene models resulting from the annotation pipeline and transcriptome assembly.

**Fig. S4.** Transcriptome statistics. We followed the transcriptome assembly quality assessment of Trinity (https://github.com/trinityrnaseq/trinityrnaseq/wiki/Transcriptome-Assembly-Quality-Assessment).

**Fig. S5.** Genome annotation pipeline used on the *Lucifuga dentata* draft genome.

**Fig. S6.** Interspersed repeat landscape of *Lucifuga dentata*.

**Fig. S7.** List of eye genes retrieved from cavefishes and related species. Colors represent the type of LoF mutation. When higher than one, the number of LoF mutations is also reported.

**Fig. S8.** A large deletion between GNL3L and SWS2 at the origin of LWS gene loss in Ophidiiformes. This figure was generated using SimpleSynteny (Veltri D., Malapi-Wight M. and Crouch J.A. SimpleSynteny: a web-based tool for visualization of microsynteny across multiple species. *Nucleic Acids Research* 44(W1):W41-W45, 2016, doi:10.1093/nar/gkw330).

**Fig. S9.** Distribution of the effective segment size generated by random insertion of STOP codons and frameshifts (100,000 simulations).

**Fig. S10.** Frameshift size distribution for each dataset.

**Fig. S11.** Observed and theoretical frequencies of different types of LoF mutations in three gene sets, and the frequency of different types of mutations found in eye genes of fossorial mammals (Emerling CA, Springer MS. 2014. Eyes underground: Regression of visual protein networks in subterranean mammals. Molecular Phylogenetics and Evolution 78:260-270).

**Fig. S12.** (A) Number of difference per gene between *Astyanax mexicanus* morphs and between *Lucifuga dentata* and *Lucifuga gibarensis*. (B) Estimation of ω for each eye gene. Grey lines represent values of dn or ds < 0.01, leading to non-reliable estimations of ω.

**Fig. S13.** Estimations of ω with concatenated sequences. Branch colors are scaled depending on the ω values. Trees were generated using ggtree (Yu, G., Smith, D.K., Zhu, H., Guan, Y. and Lam, T.T.-Y. (2017), ggtree: an R package for visualization and annotation of phylogenetic trees with their covariates and other associated data. Methods Ecol Evol, 8: 28-36. doi:10.1111/2041-210X.12628).

**Fig. S14.** RELAX results with species assigned as test branch for eye genes. The k parameter and p-value are displayed along with ω plots.

**Fig. S15.** RELAX results with species assigned as test branch for circadian clock genes. The k parameter and p-value are displayed along with ω plots.

**Fig. S16.** RELAX results with species assigned as test branch for pigmentation genes. The k parameter and p-value are displayed along with ω plots.

**Fig. S17.** Empirical cumulative distributions of MutPred2 scores. The number of scores is indicated between parenthesis. 100 neutral simulations were performed for each dataset with 54 random non synonymous mutations in eye genes, 36 in circadian clock genes and 232 in pigmentation genes, which are the number of non-synonymous mutations found in *Astyanax mexicanus* cavefish. The statistical significance of the difference between each pair of distributions was assessed using the Kolmogorov-Smirnov test (significant differences are shown on a red background whereas non-significant differences are shown on a green background).

**Fig. S18.** Empirical cumulative distributions of Grantham’s distances. The number of distances is indicated between parenthesis. 100 neutral simulations were performed for each dataset with 54 random non synonymous mutations in eye genes, 36 in circadian clock genes and 232 in pigmentation genes which are the number of non-synonymous mutations found in *Astyanax mexicanus* cavefish. The statistical significance of the difference between each pair of distributions was assessed using the Kolmogorov-Smirnov test (significant differences are shown on a red background whereas non-significant differences are shown on a green background).

**Fig. S19.** Effect of the Transition/Transversion ratio on the cumulative distribution of MutPred2 scores in simulated amino acid substitutions.

**Fig. S20.** Fit of mixture distributions of MutPred2 scores with the distributions found in two *Lucifuga* spp. and two *Astyanax mexicanus* morphs. The p-values of Kolomogorv-Smirnov tests between the observed distributions in each species and mixture distributions were plotted according to different proportions of mutations that reached fixation under relaxed selection.

**Fig. S21.** Distributions of MutPred2 scores in three *Sinocyclocheilus* species, *Danio rerio* and in simulations of substitutions without selection. The number of substitutions in each lineage is given between parenthesis. One hundred simulations were performed with each gene set. In each simulation 54 non-synonymous mutations were generated in eye genes, 36 in circadian clock genes and 232 in pigmentation genes, those numbers corresponding to the numbers of non-synonymous mutations found in *Astyanax mexicanus* cavefish.

**Fig. S22.** Fit of mixture distributions of MutPred2 scores with the distributions found in three *Sinocyclocheilus* species. The p-values of Kolomogorv-Smirnov tests between the observed distributions in each species and mixture distributions were plotted according to different proportions of mutations that reached fixation under relaxed selection.

**Table S1.** List of LoF mutations found in *Lucifuga dentata* and *Lucifuga gibarensis* genomes, and their coverage. LoF mutations in red were also found in the transcriptome of *L. dentata*.

**Data_Supp1.** Summary of the number of genes retrieved from each species and for each gene set, along with the number of pseudogenes and the number of LoF mutations.

**Data_Supp2.** Sequences predicted with exonerate and ID of sequences retrieved from Ensembl.

**Data_Supp3.** Results obtained with different methods for dating relaxed selection on eye genes in *Lucifuga dentata*.

Description of Supplementary files content:

Divergence_values: Pairwise nucleotidic distances between species for each gene set. Lucifuga_Supplementary_files_Genome: Original GFF3 file with functional annotations and scaffolds smaller than 200 bp not uploaded to NCBI.

MutPred2_Results: Raw output of MutPred2. Parsed results files to be used with the script provided in github (MutPred2_Script.R) are also provided.

Phylogenies: Gene phylogenies computed with iQTree and displayed with iTOL. The model used for each phylogeny can be found on the “Models” folder.

Concatenated_Alignments: Concatenated alignments for vision, circadian and pigmentation genes.

