## Supplementary figures and images for "Contrasted gene decay in subterranean vertebrates: insights from cavefishes and fossorial mammals"

### Supplementary fig. S1

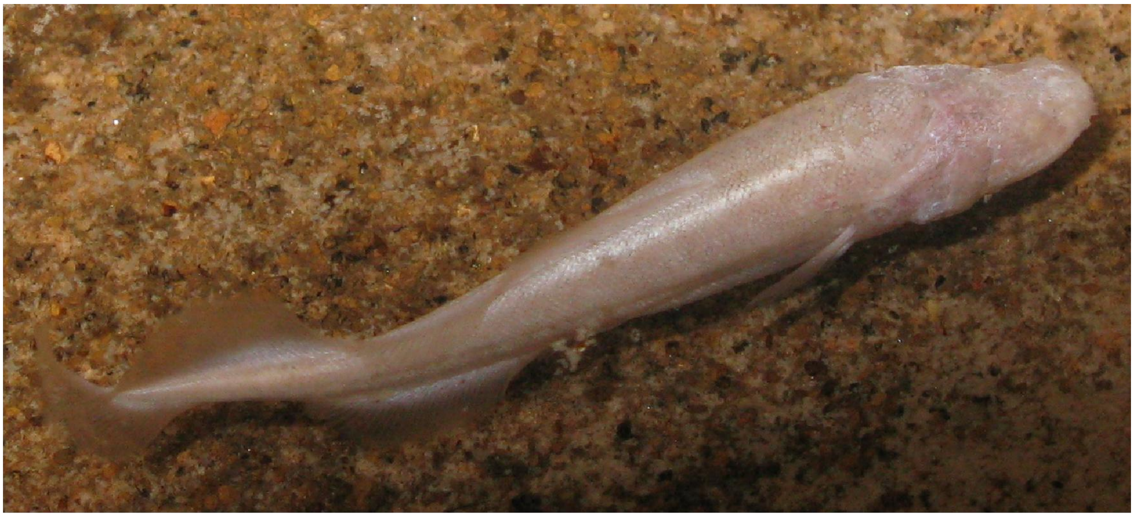

*Lucifuga dentata*

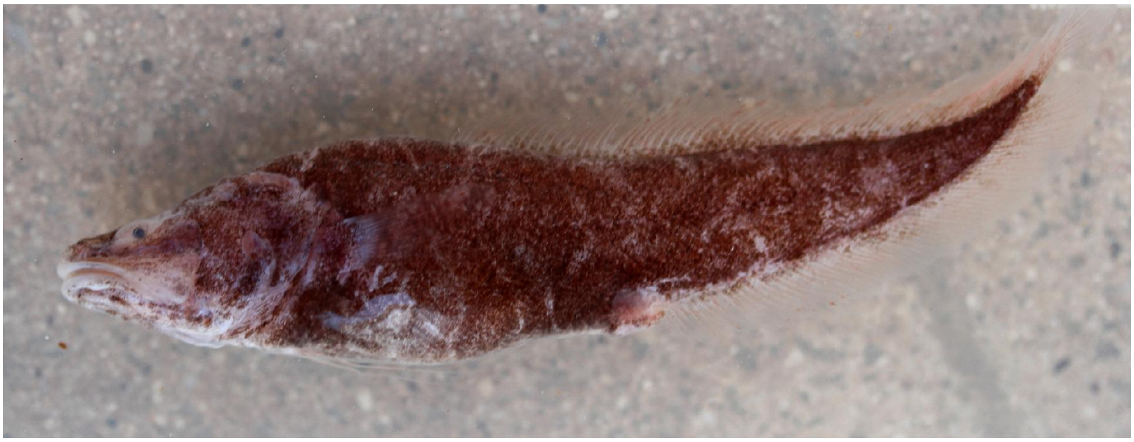

*Lucifuga gibarensis*

### Supplementary fig. S2

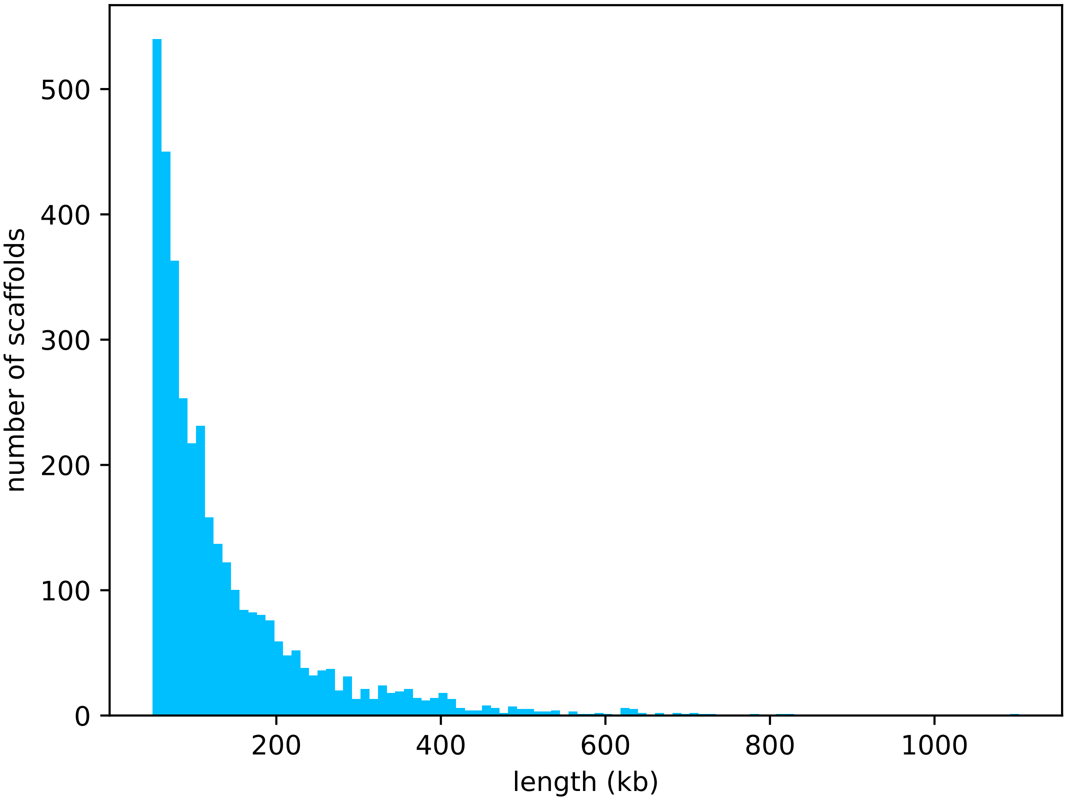

### Supplementary fig. S3

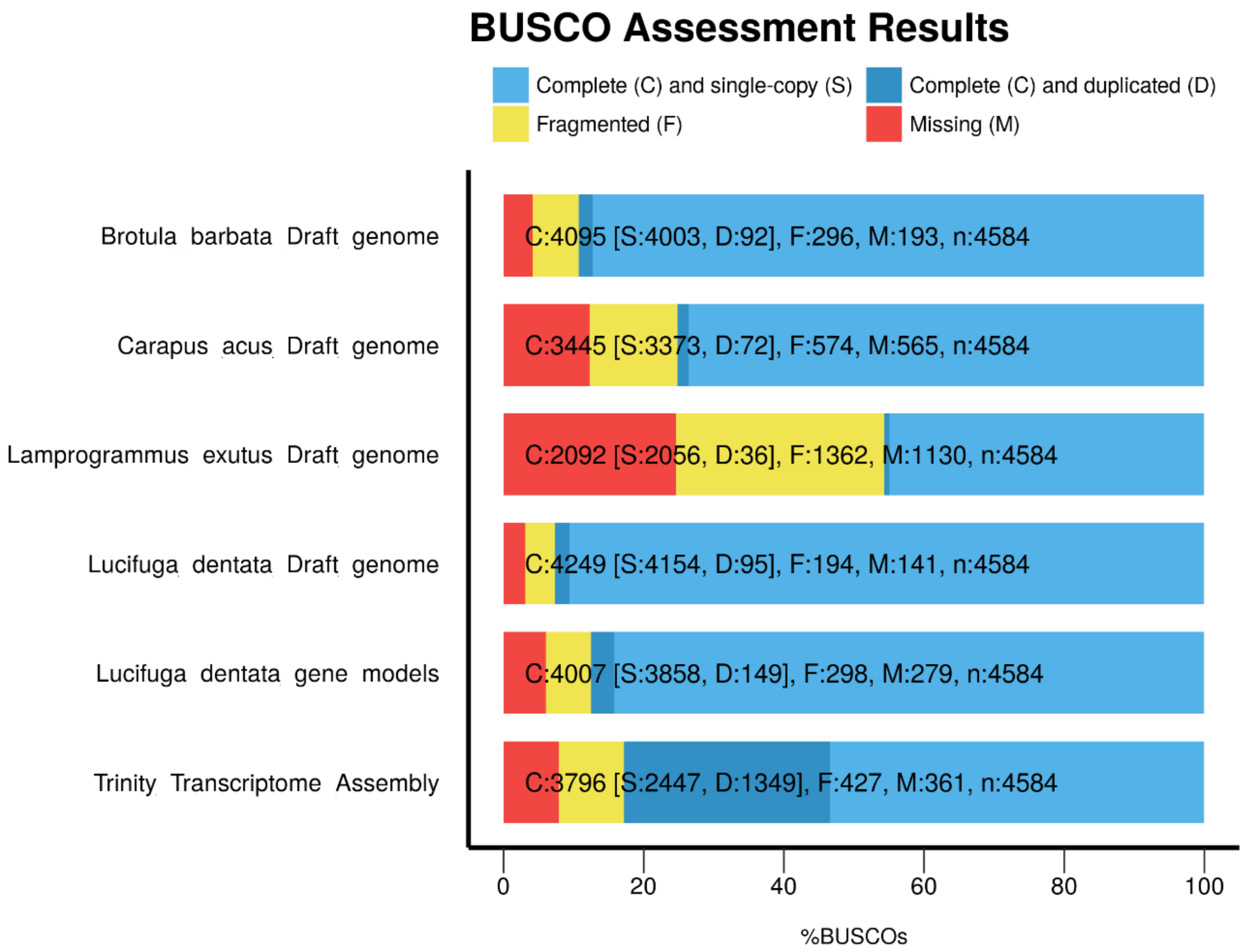

### Supplementary fig. S5

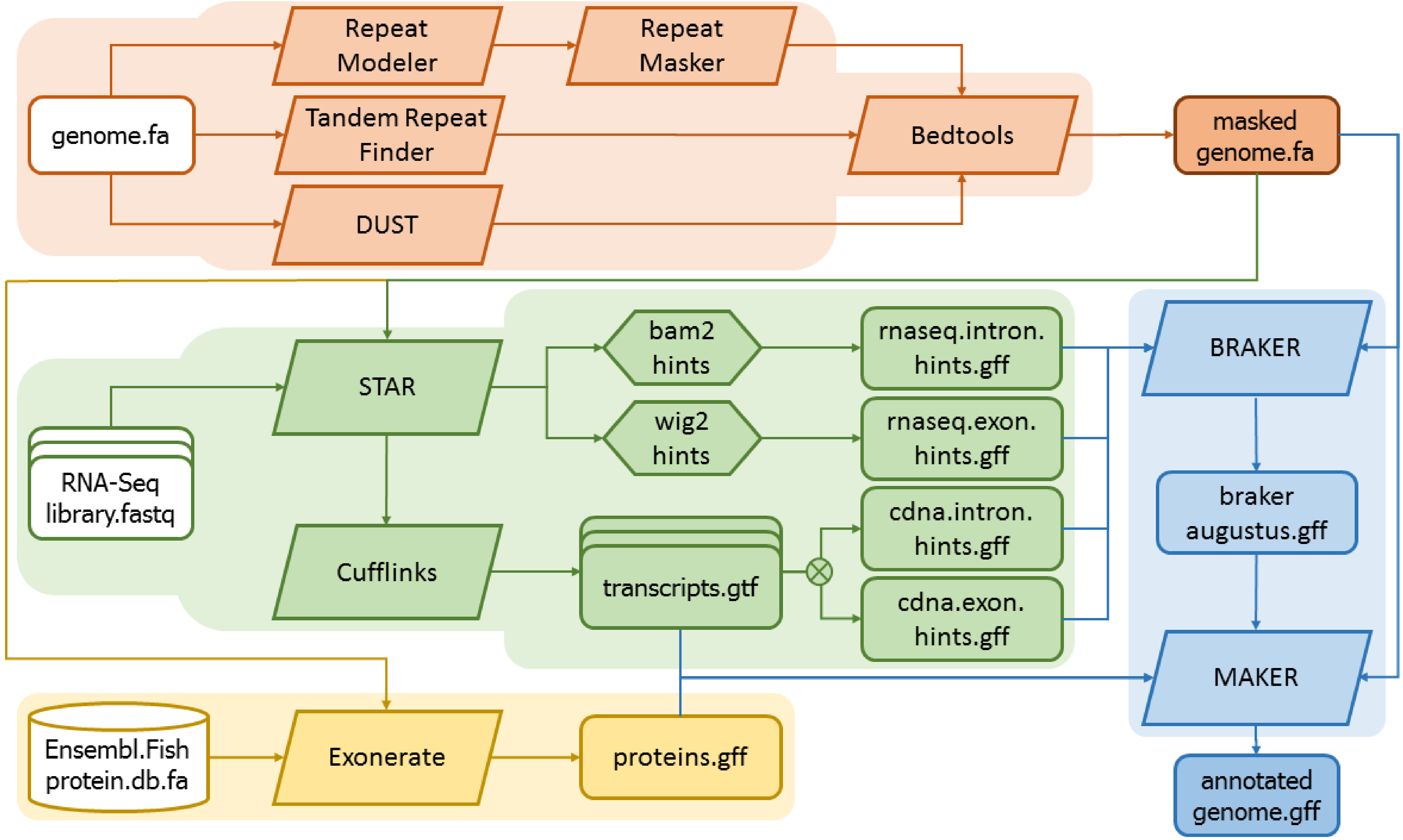

### Supplementary fig. S6

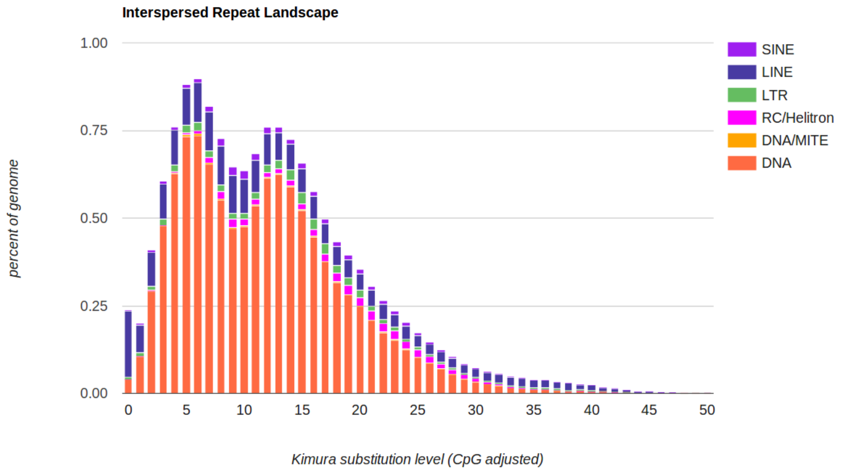

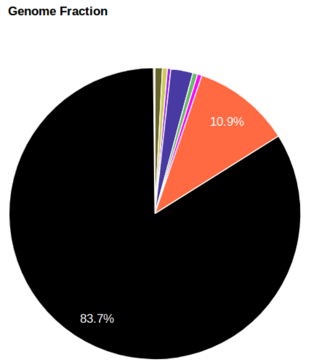

### Supplementary fig. S7

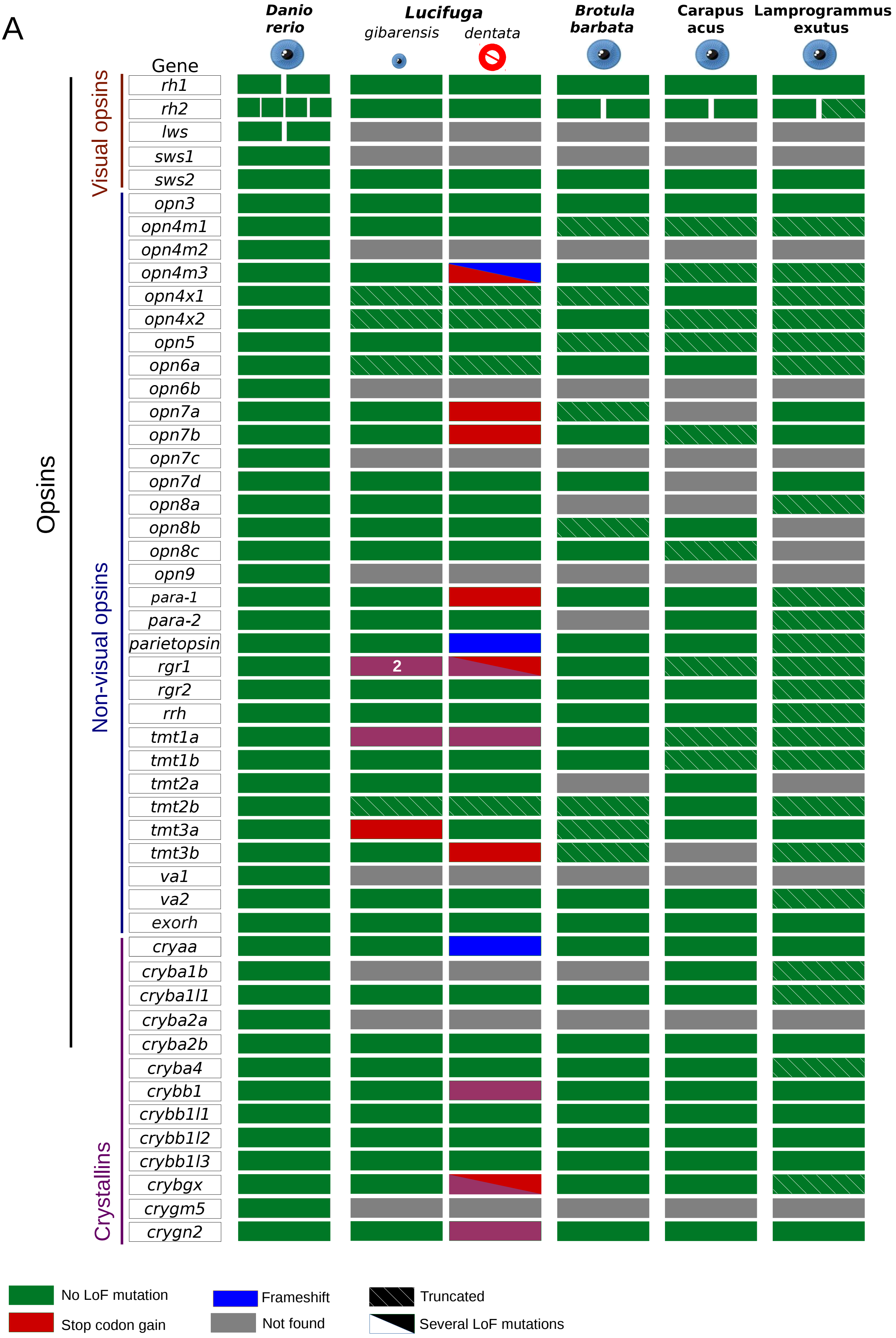

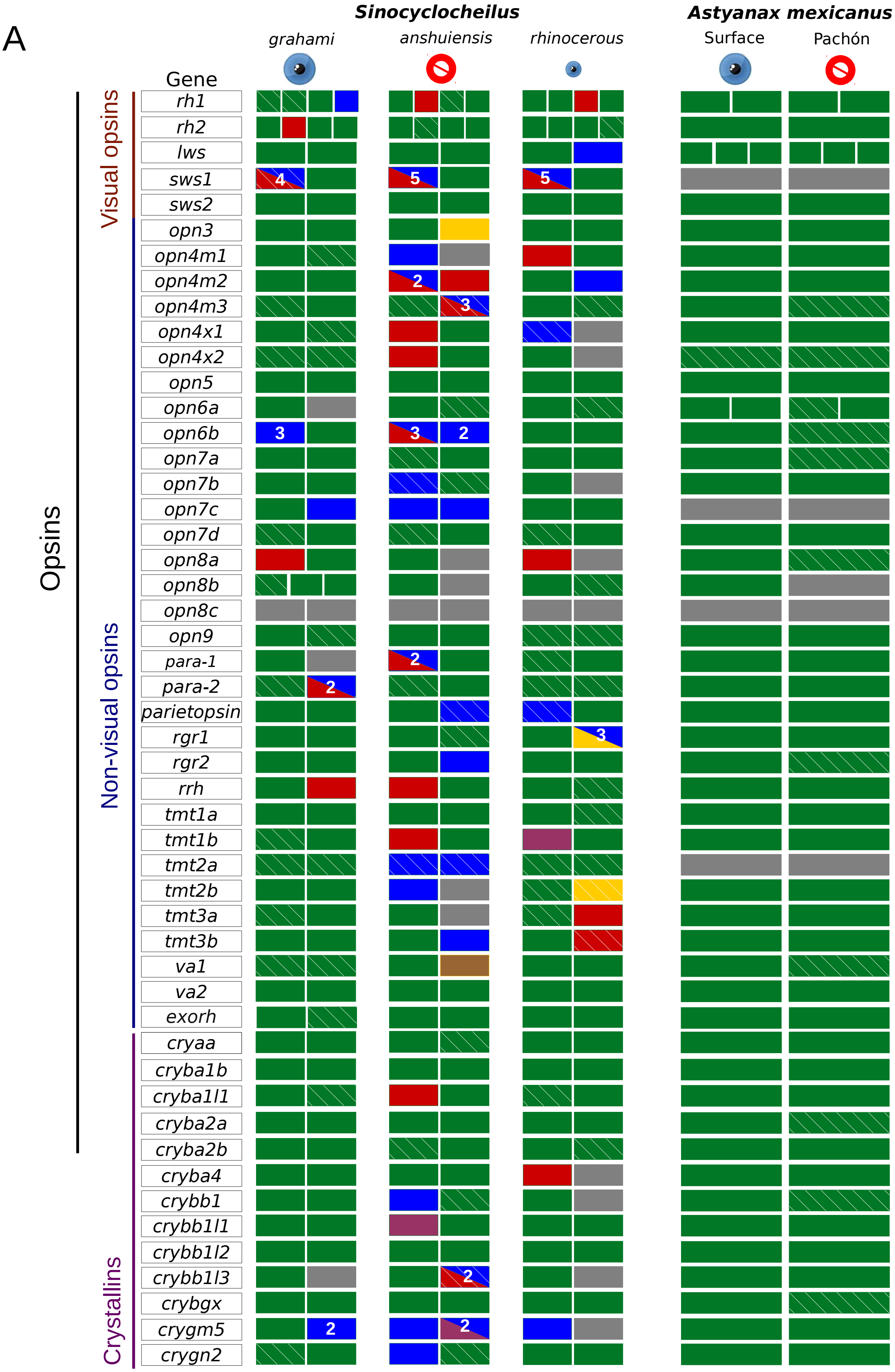

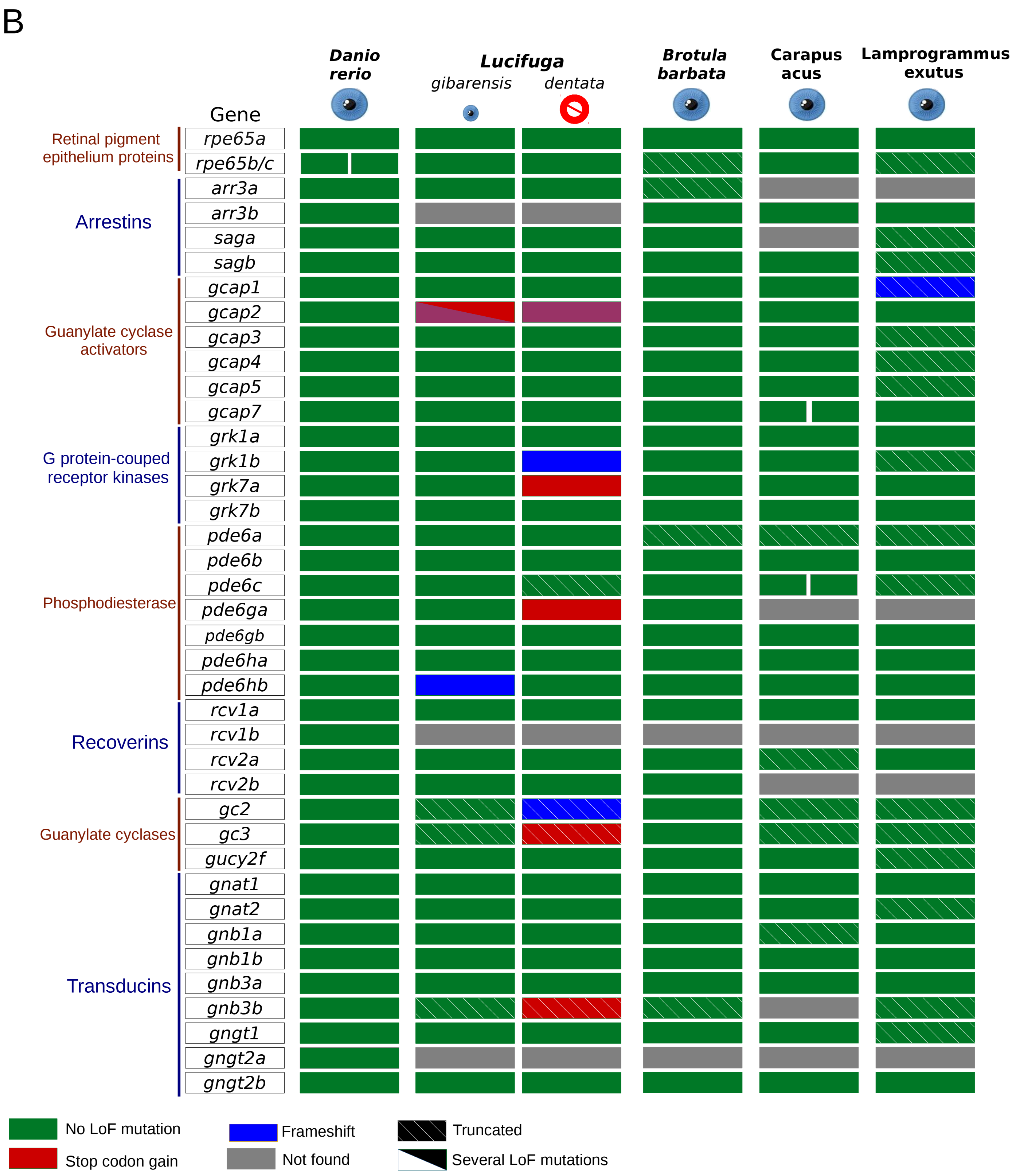

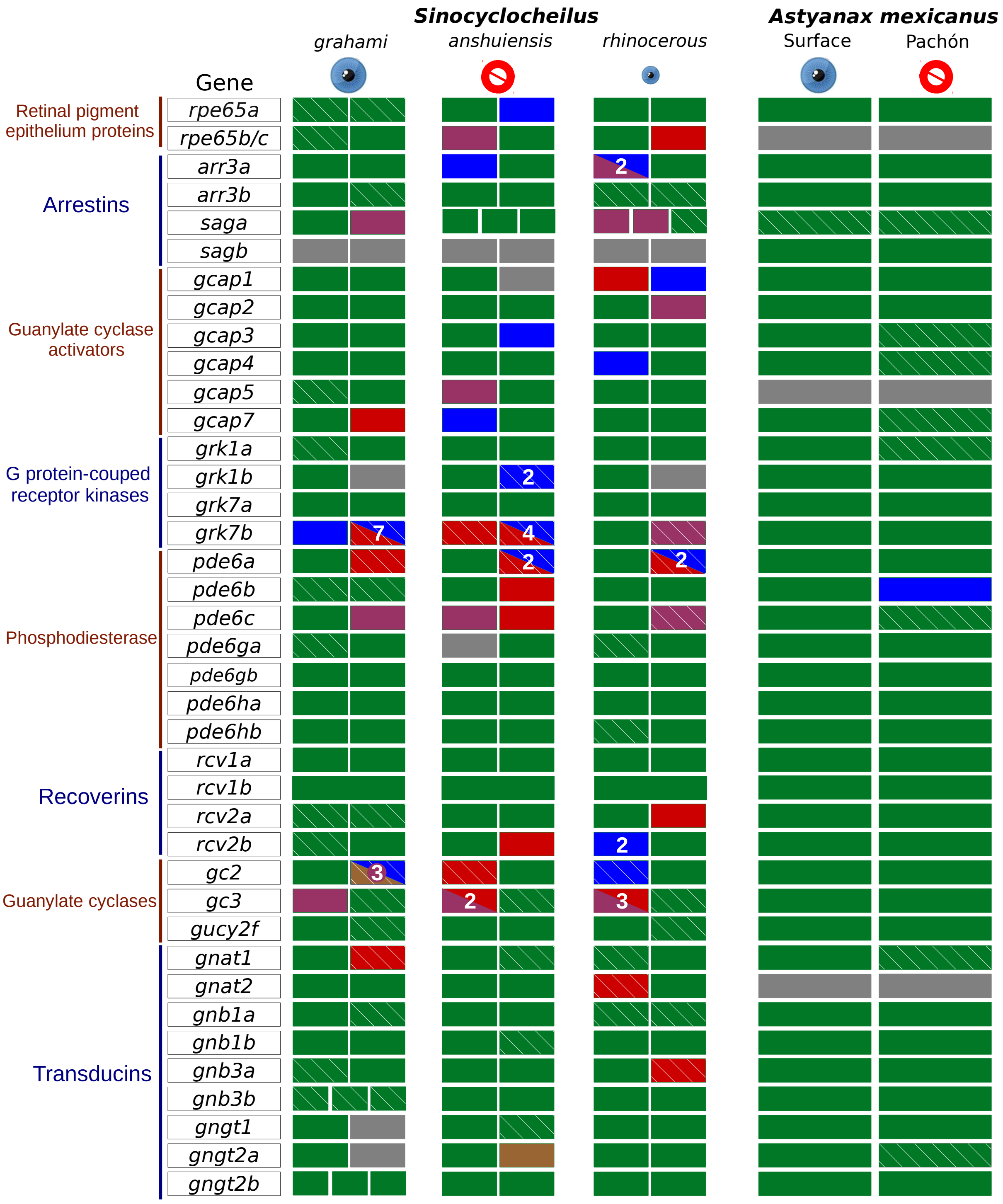

### Supplementary fig. S8

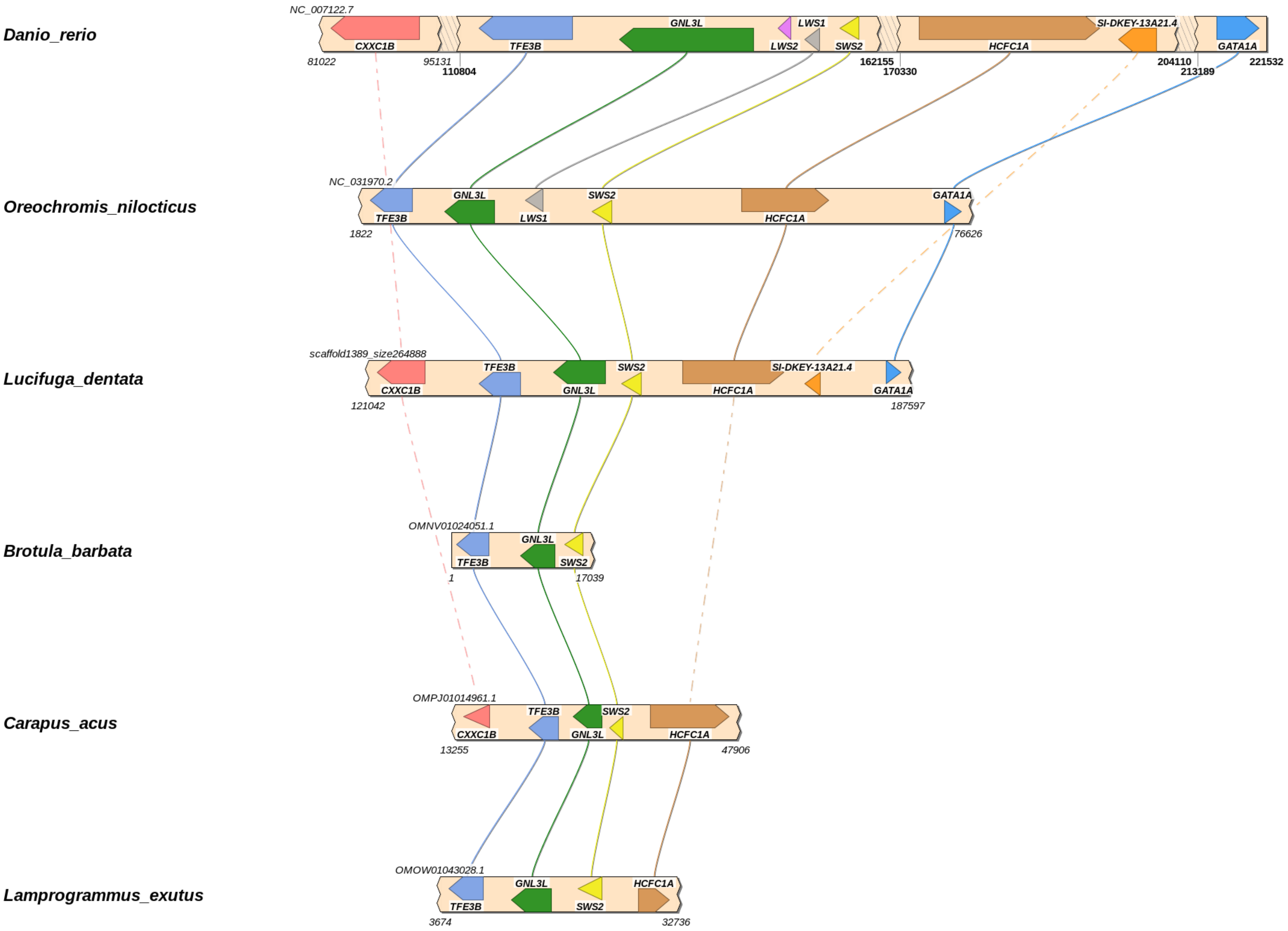

### Supplementary fig. S9

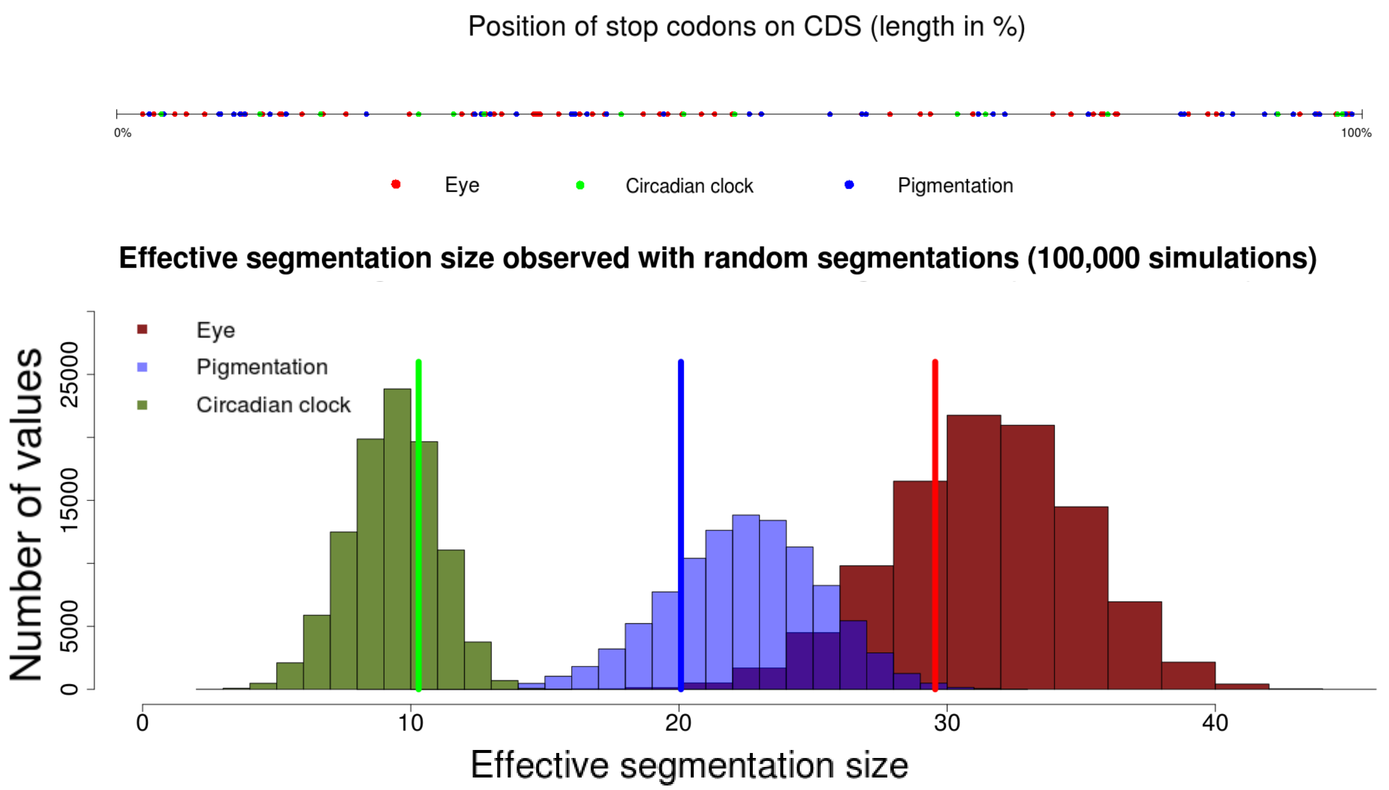

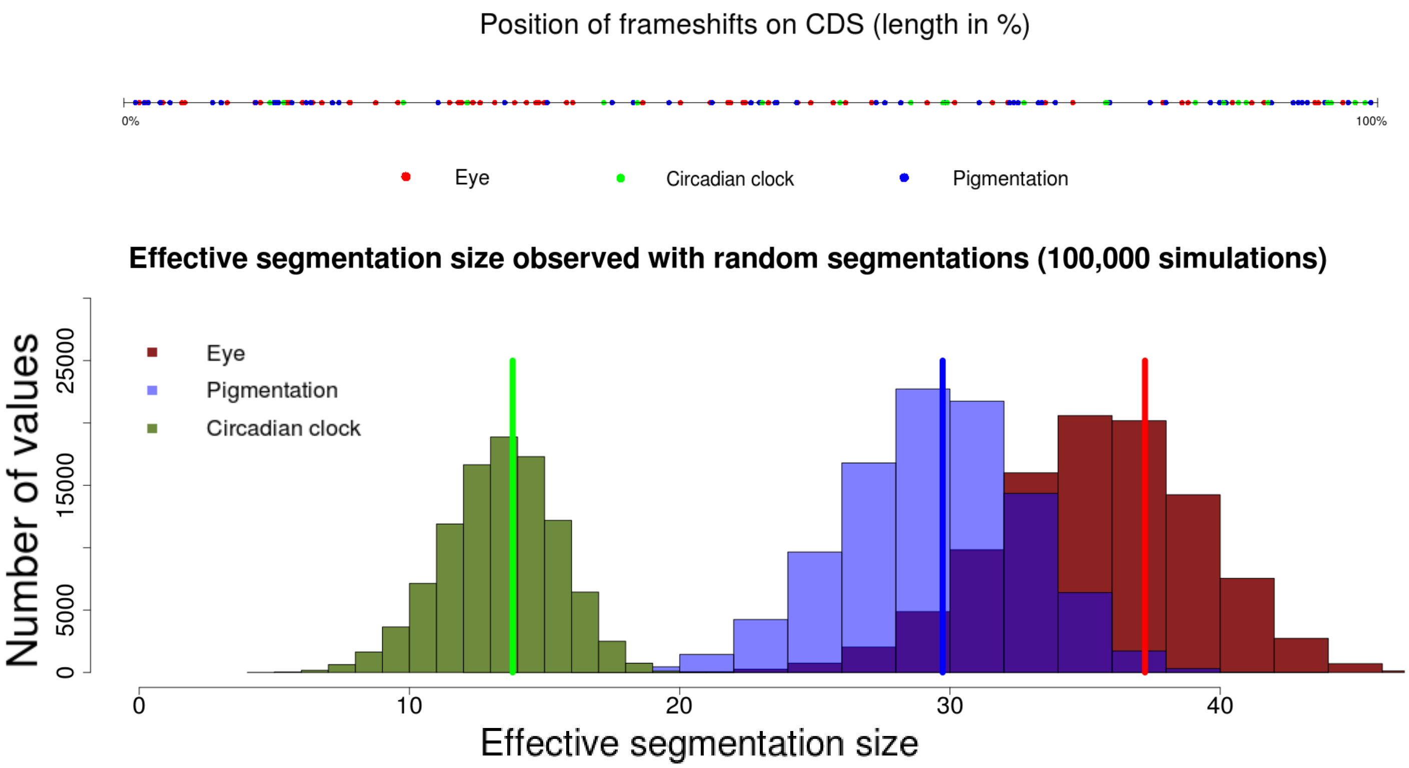

### Supplementary fig. S10

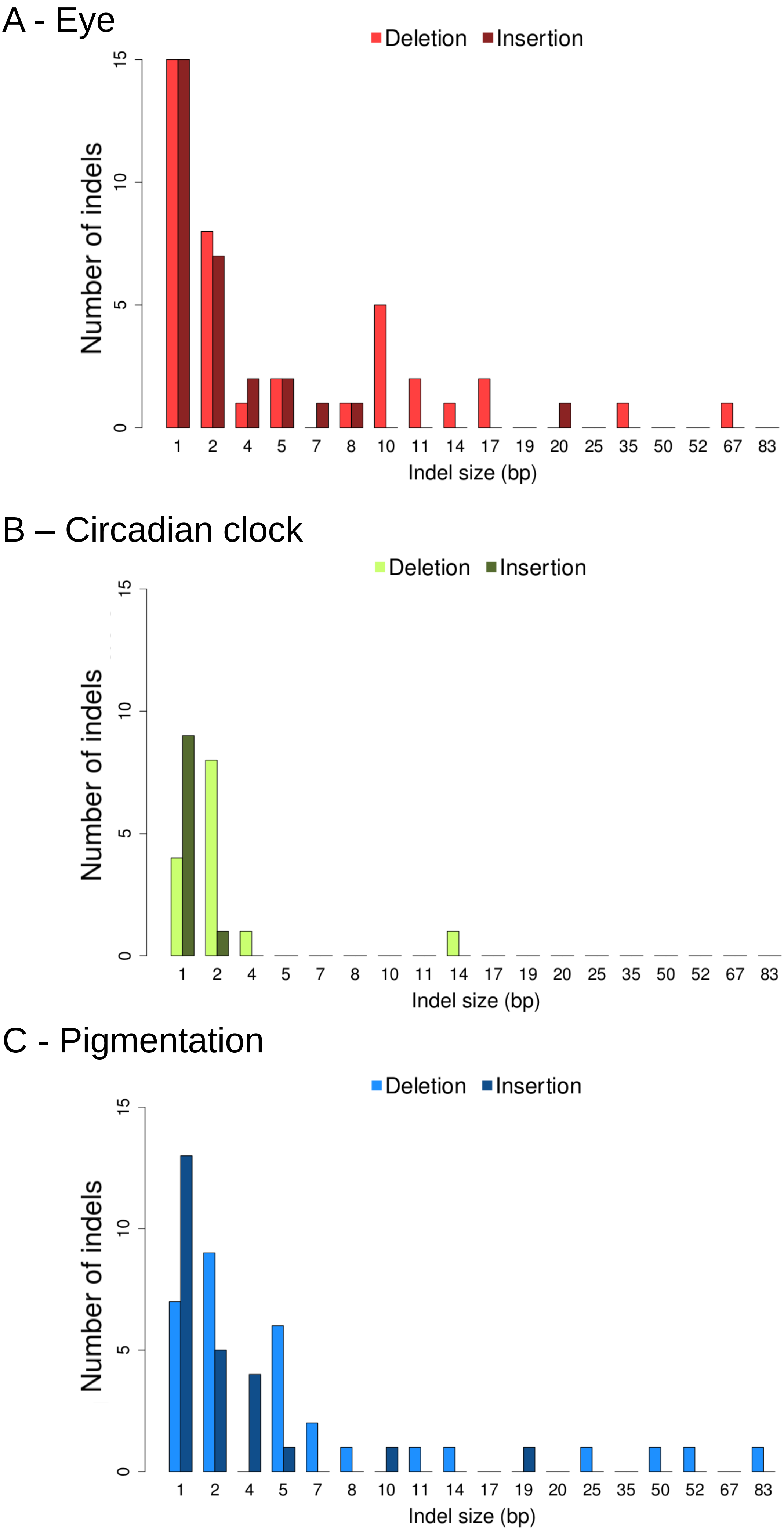

### Supplementary fig. S11

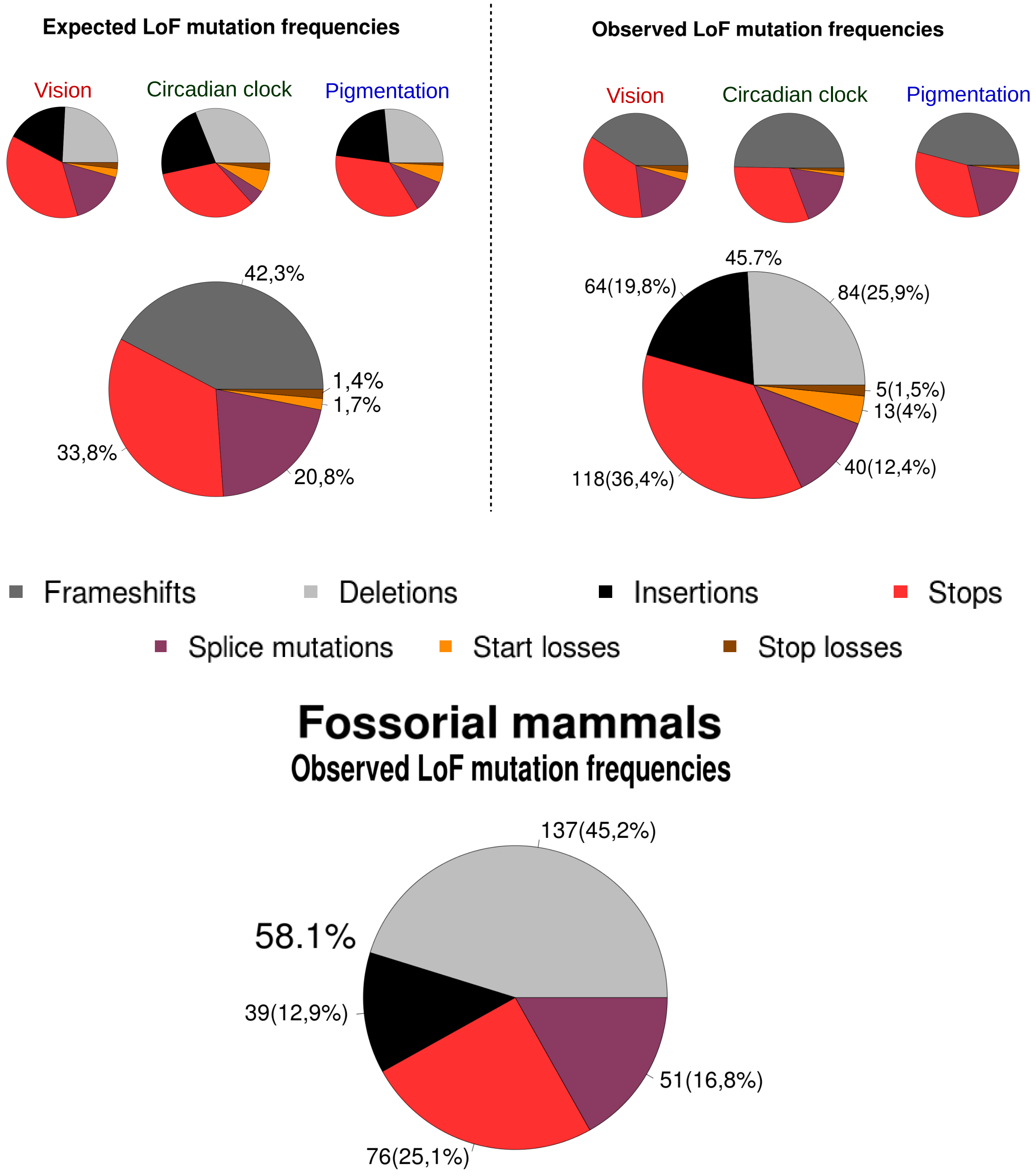

### Supplementary fig. S12

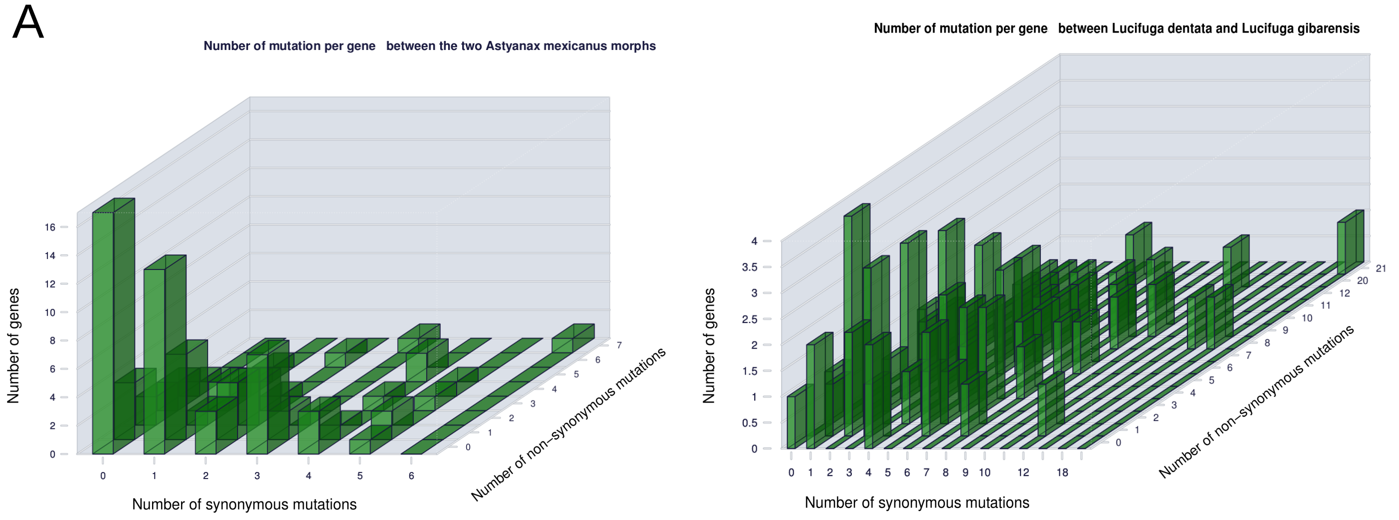

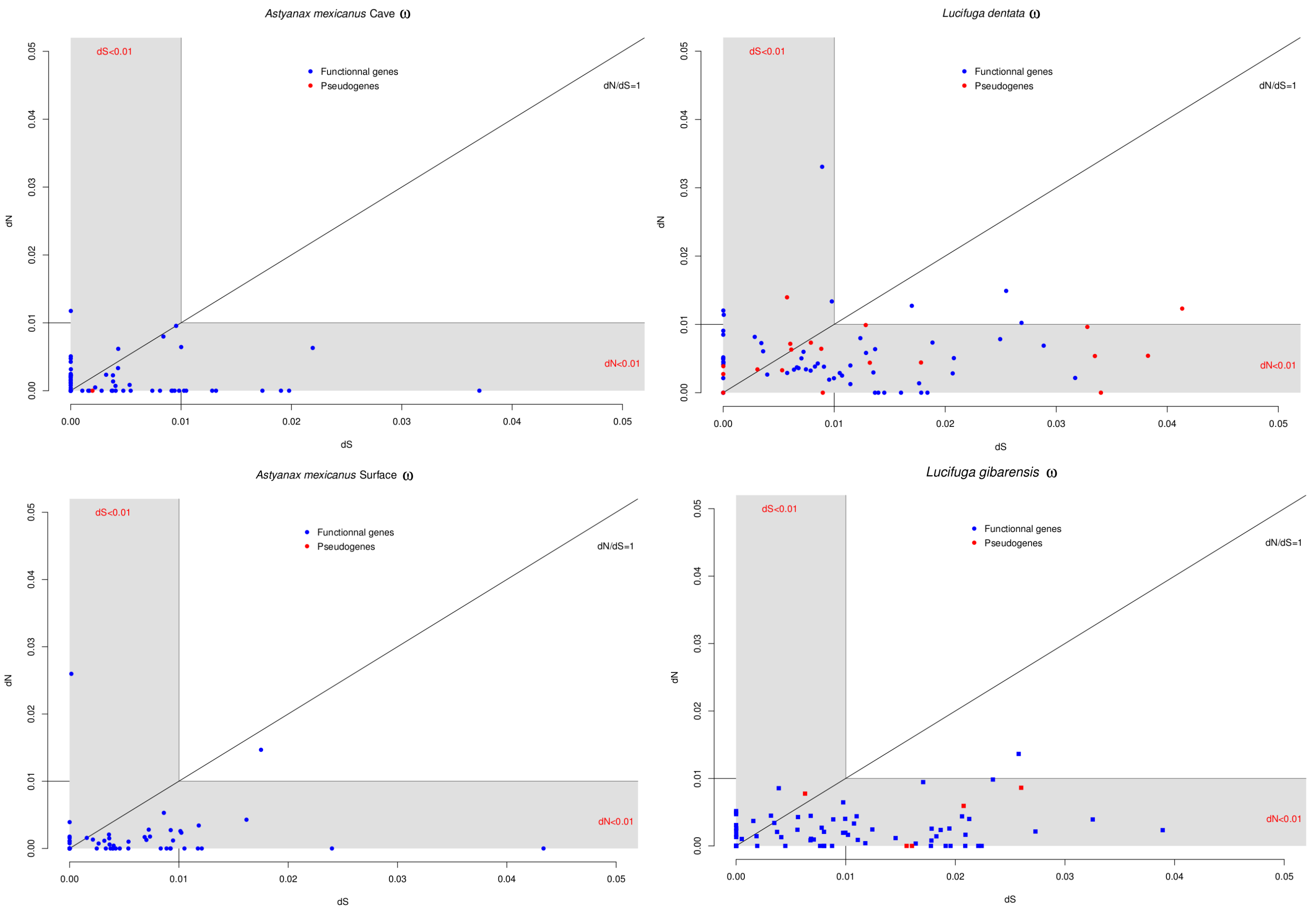

### Supplementary fig. S19

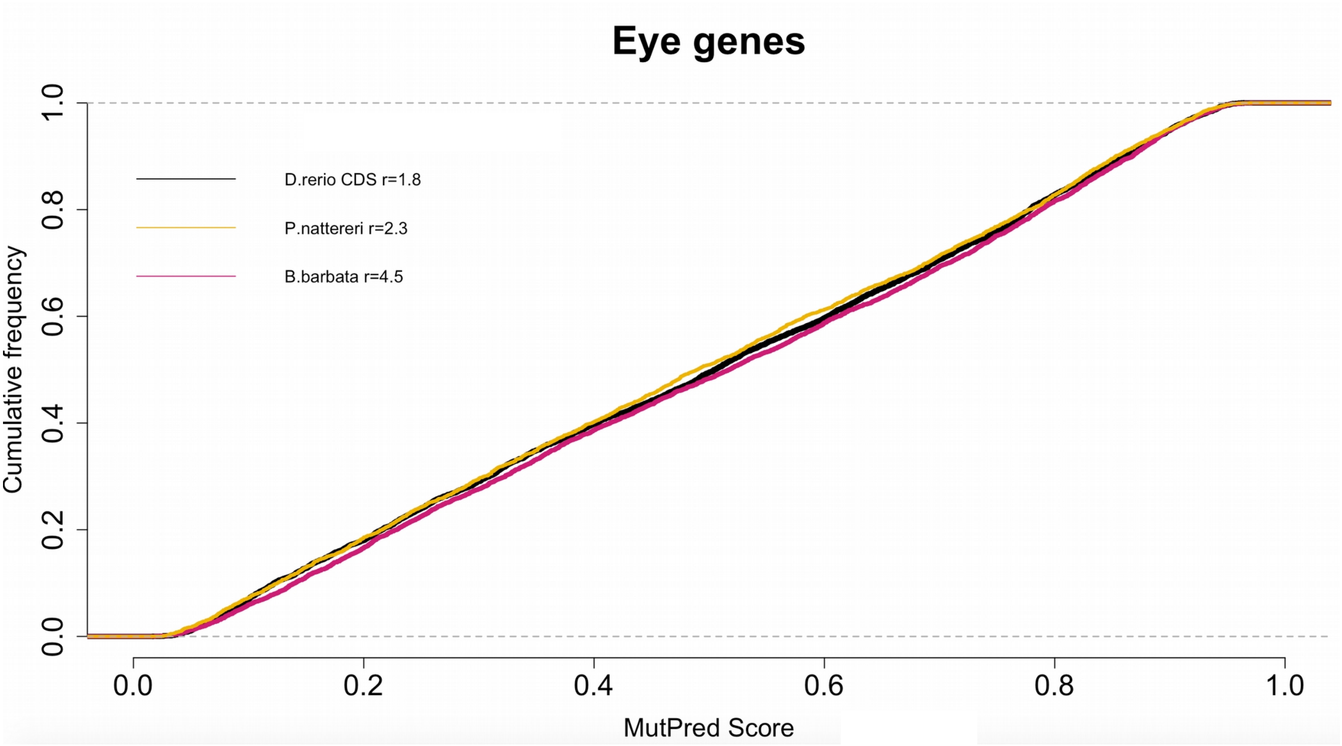

### Supplementary fig. S20

#### Eye

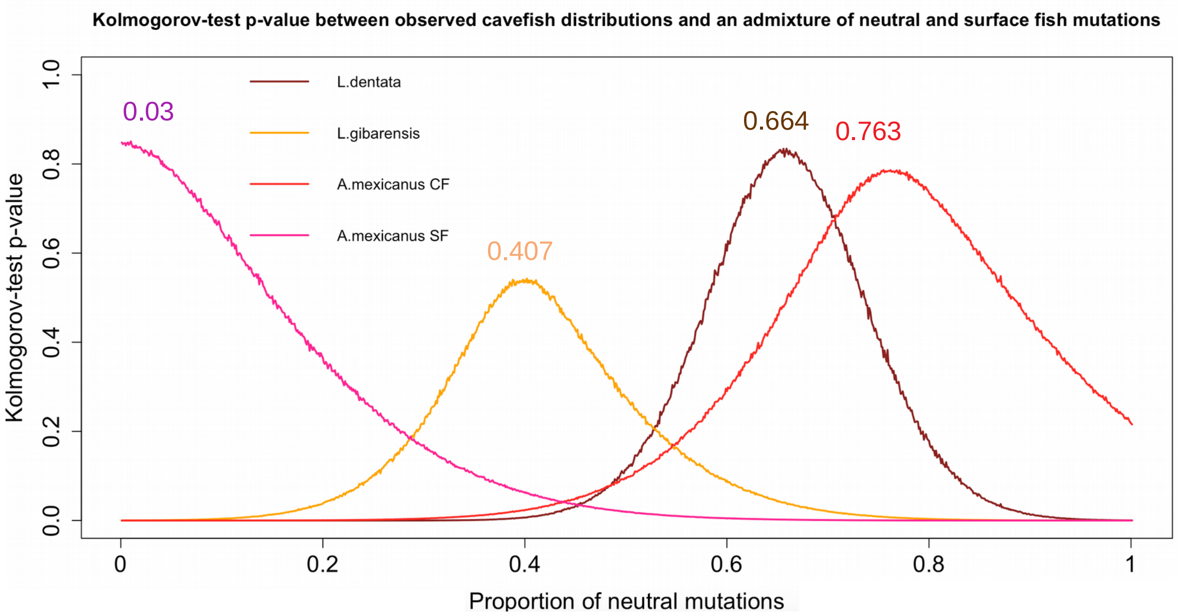


#### Circadian clock

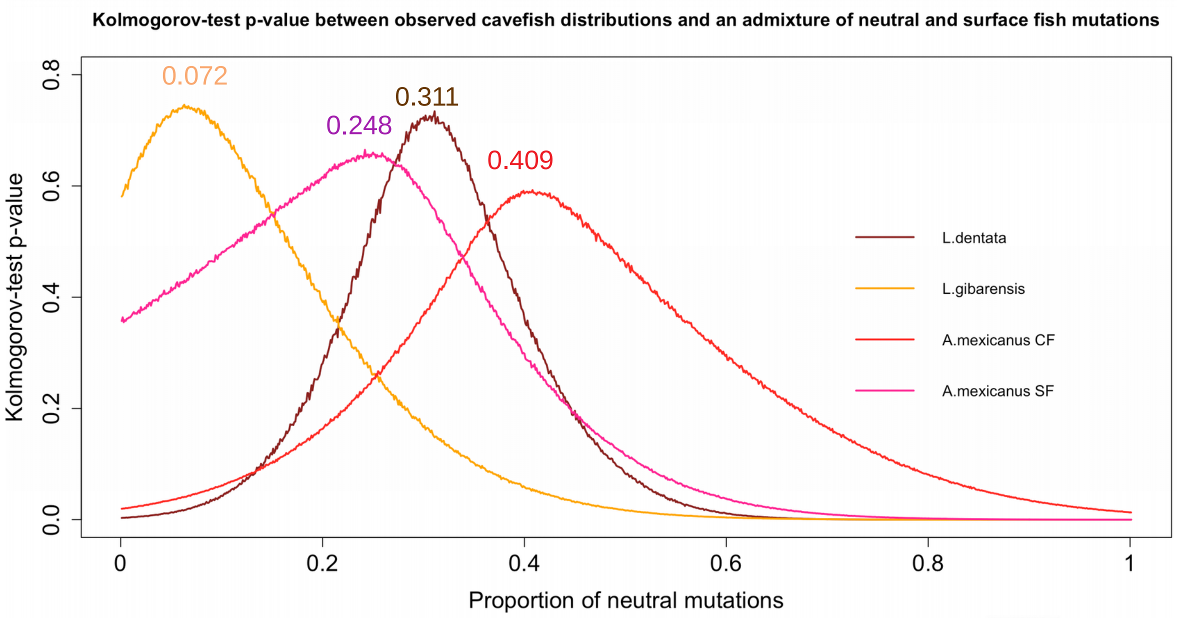


#### Pigmentation

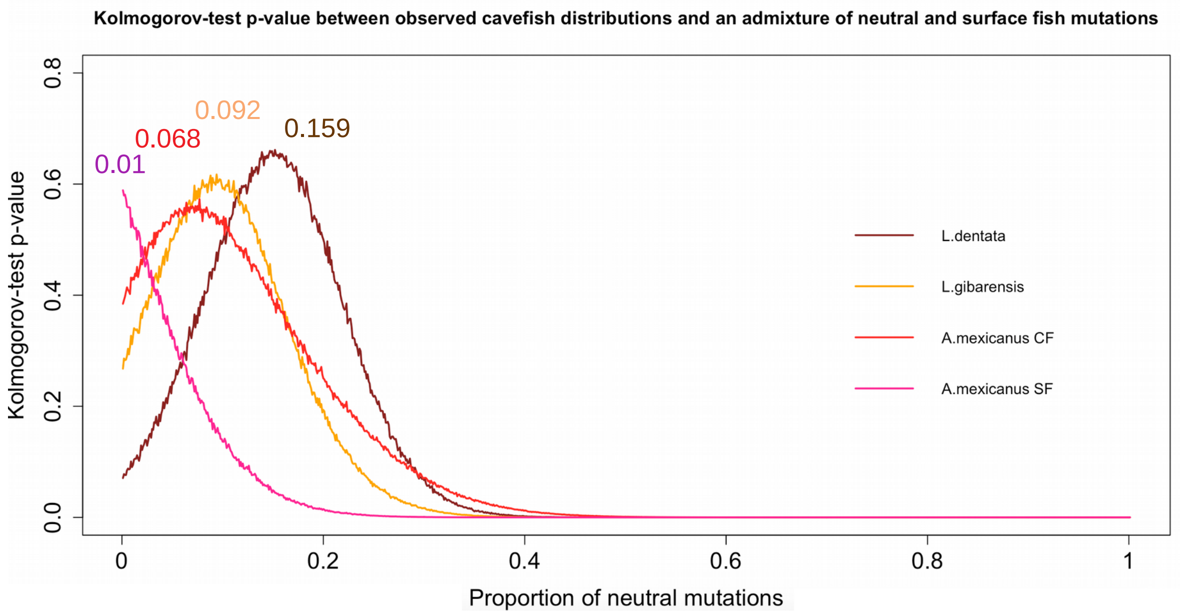

### Supplementary fig. S22

#### Eye

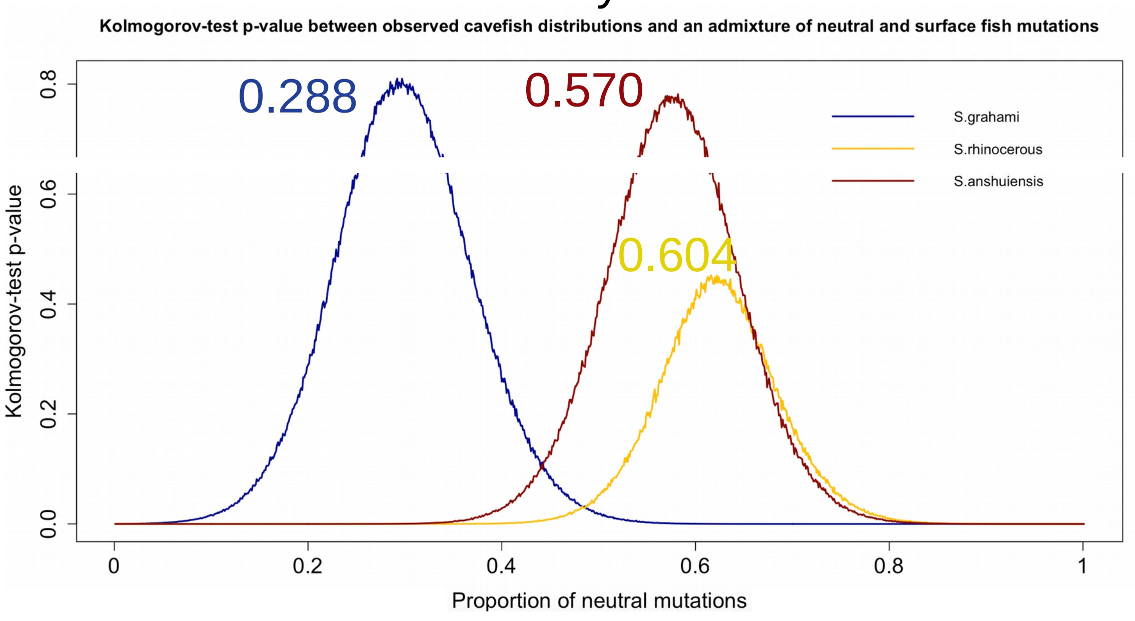


#### Circadian clock

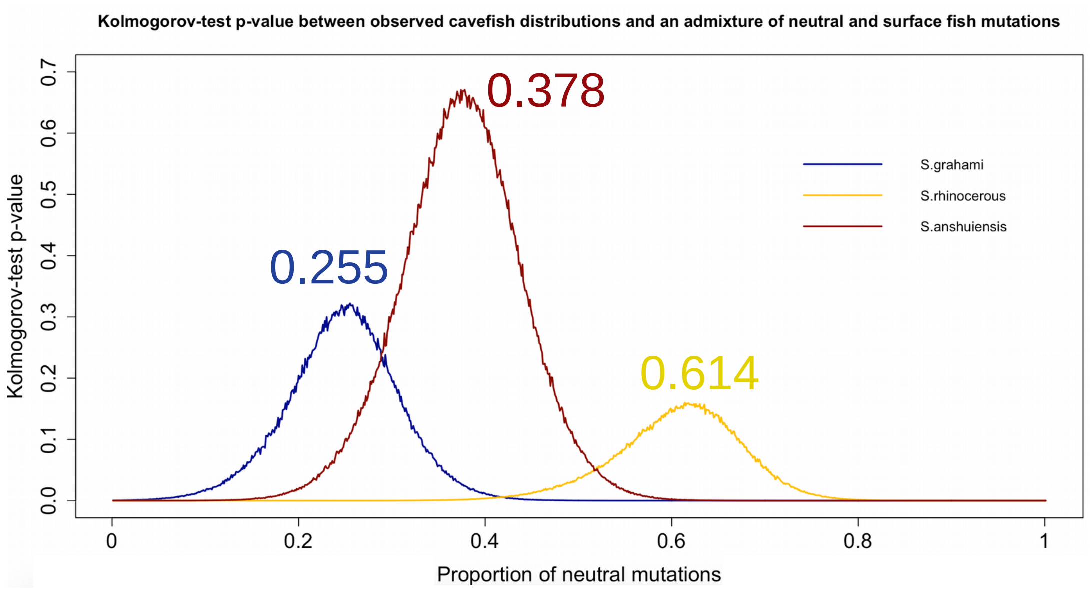


#### Pigmentation

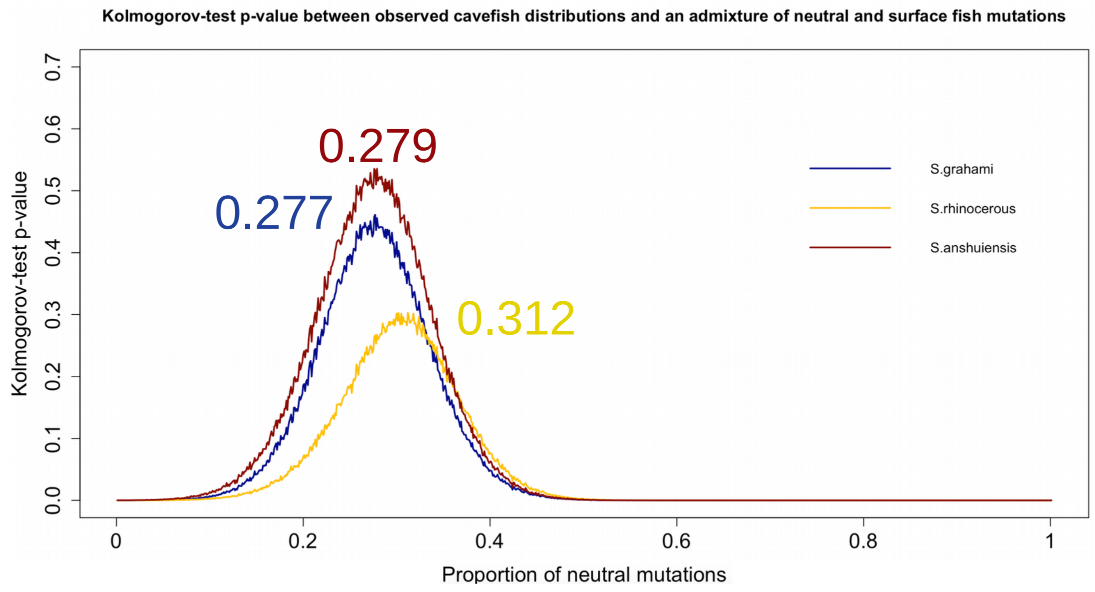
