## Supplementary fig. S4 for "Contrasted gene decay in subterranean vertebrates: insights from cavefishes and fossorial mammals"

######################## RNA Seq Read Representation by Trinity Assembly ########################

159178098 reads; of these:
  159178098 (100.00%) were paired; of these:
    25554370 (16.05%) aligned concordantly 0 times
    22283453 (14.00%) aligned concordantly exactly 1 time
    111340275 (69.95%) aligned concordantly >1 times
    ----
    25554370 pairs aligned concordantly 0 times; of these:
      1167404 (4.57%) aligned discordantly 1 time
    ----
    24386966 pairs aligned 0 times concordantly or discordantly; of these:
      48773932 mates make up the pairs; of these:
        24491684 (50.21%) aligned 0 times
        4741471 (9.72%) aligned exactly 1 time
        19540777 (40.06%) aligned >1 times
92.31% overall alignment rate

######################## Counting Full Length Trinity Transcripts ########################

| #hit_pct_cov_bin | count_in_bin | >bin_below |
|---|---|---|
| 100 | 7644 | 7644 |
| 90 | 2304 | 9948 |
| 80 | 1841 | 11789 |
| 70 | 1691 | 13480 |
| 60 | 1825 | 15305 |
| 50 | 1939 | 17244 |
| 40 | 2169 | 19413 |
| 30 | 2260 | 21673 |
| 20 | 2103 | 23776 |
| 10 | 793 | 24569 |

######################## Transcriptome Contig Nx Statistic ########################

################################
### Counts of transcripts, etc.
################################
Total trinity 'genes': 327313
Total trinity transcripts: 511116
Percent GC: 44.35

########################################
Stats based on ALL transcript contigs:
########################################

```
    Contig N10: 4707
    Contig N20: 3358
    Contig N30: 2545
    Contig N40: 1932
    Contig N50: 1408

    Median contig length: 368
    Average contig: 752.13
    Total assembled bases: 384426633


#######################################################
## Stats based on ONLY LONGEST ISOFORM per 'GENE':
#######################################################

    Contig N10: 3722
    Contig N20: 2418
    Contig N30: 1536
    Contig N40: 944
    Contig N50: 629

    Median contig length: 310
    Average contig: 525.20
    Total assembled bases: 171905036
```

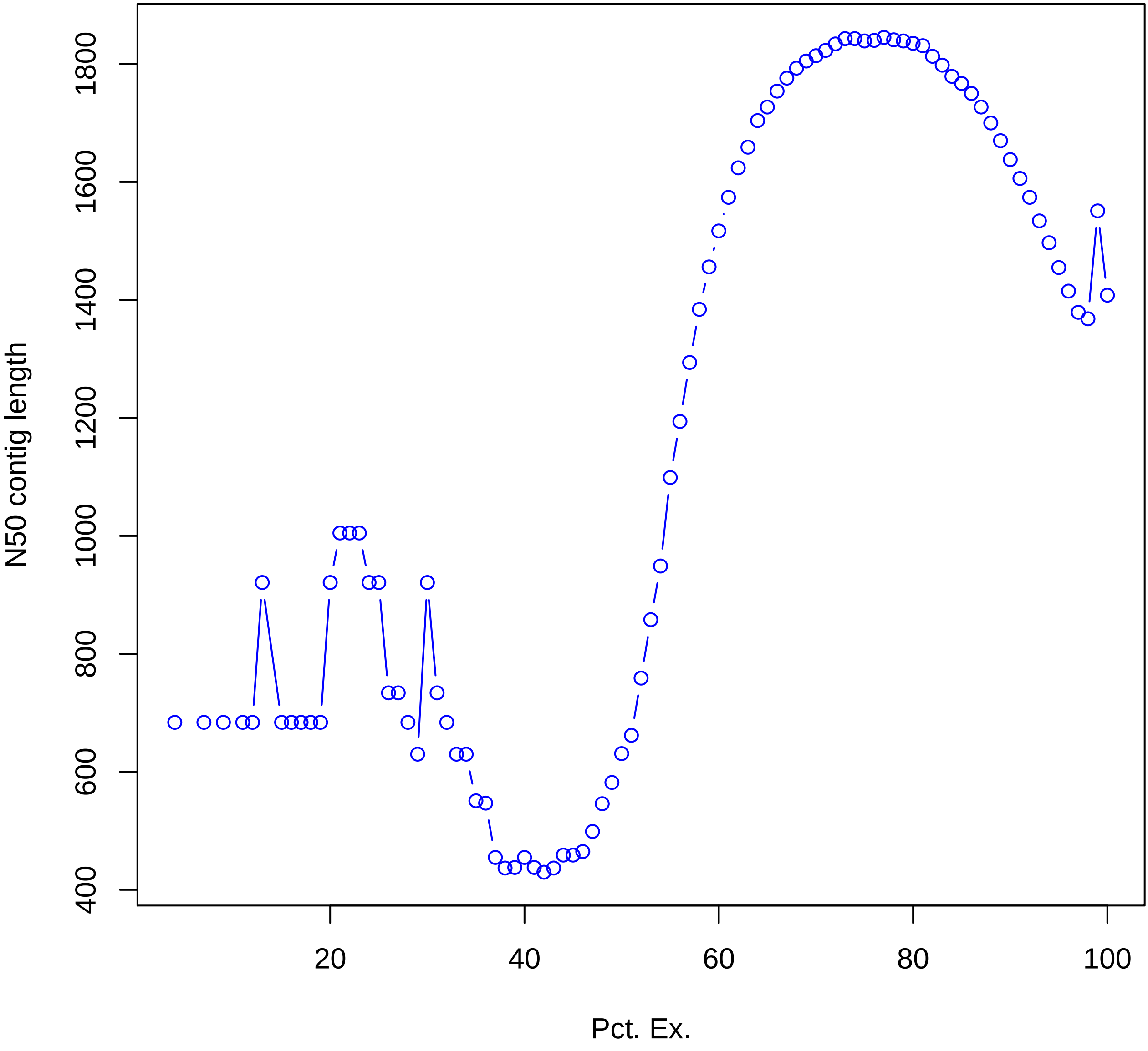
N50 contig length
400
600
800
1000
1200
1400
1600
1800
20
40
60
80
100
Pct. Ex.
