## Supplementary fig. S13 for "Contrasted gene decay in subterranean vertebrates: insights from cavefishes and fossorial mammals"

**A**

**Concatenated eye genes**

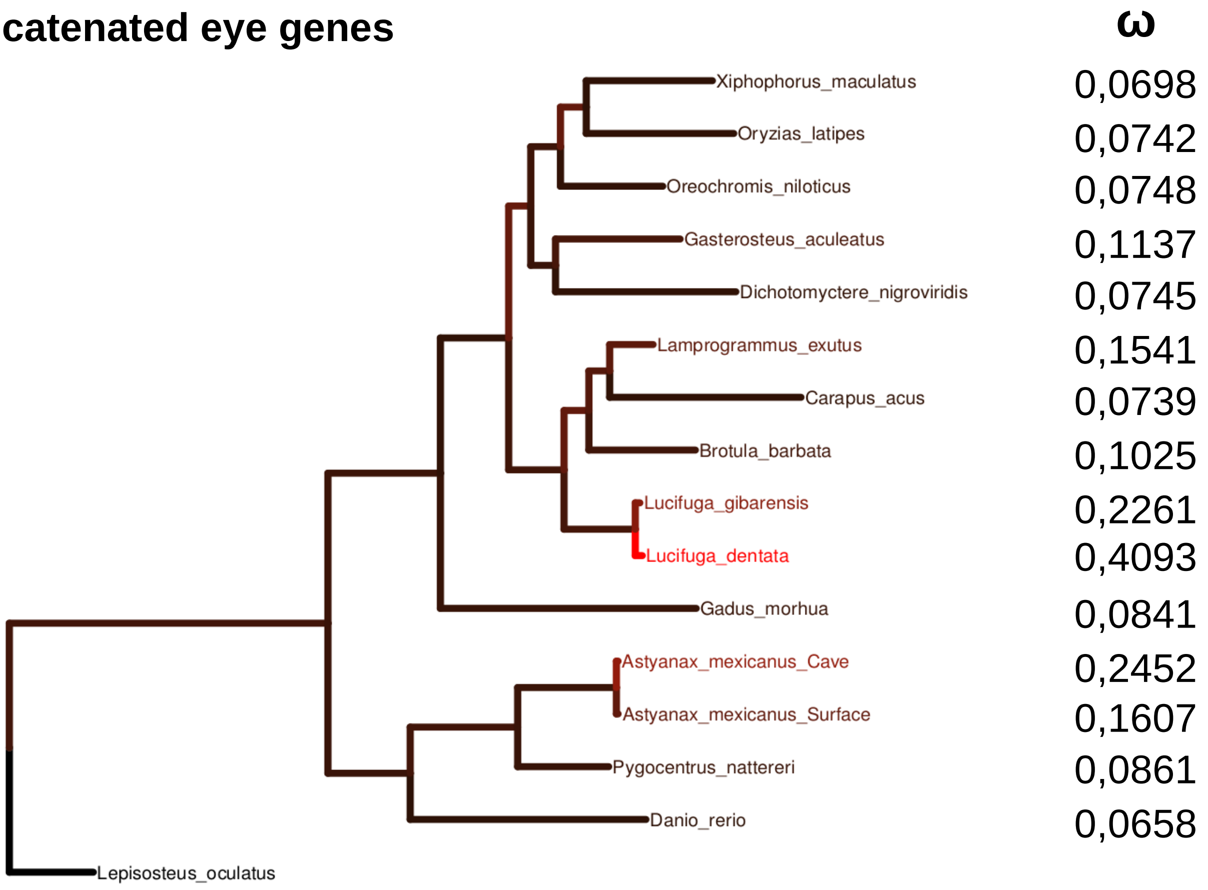


**B**

**Concatenated circadian clock genes**

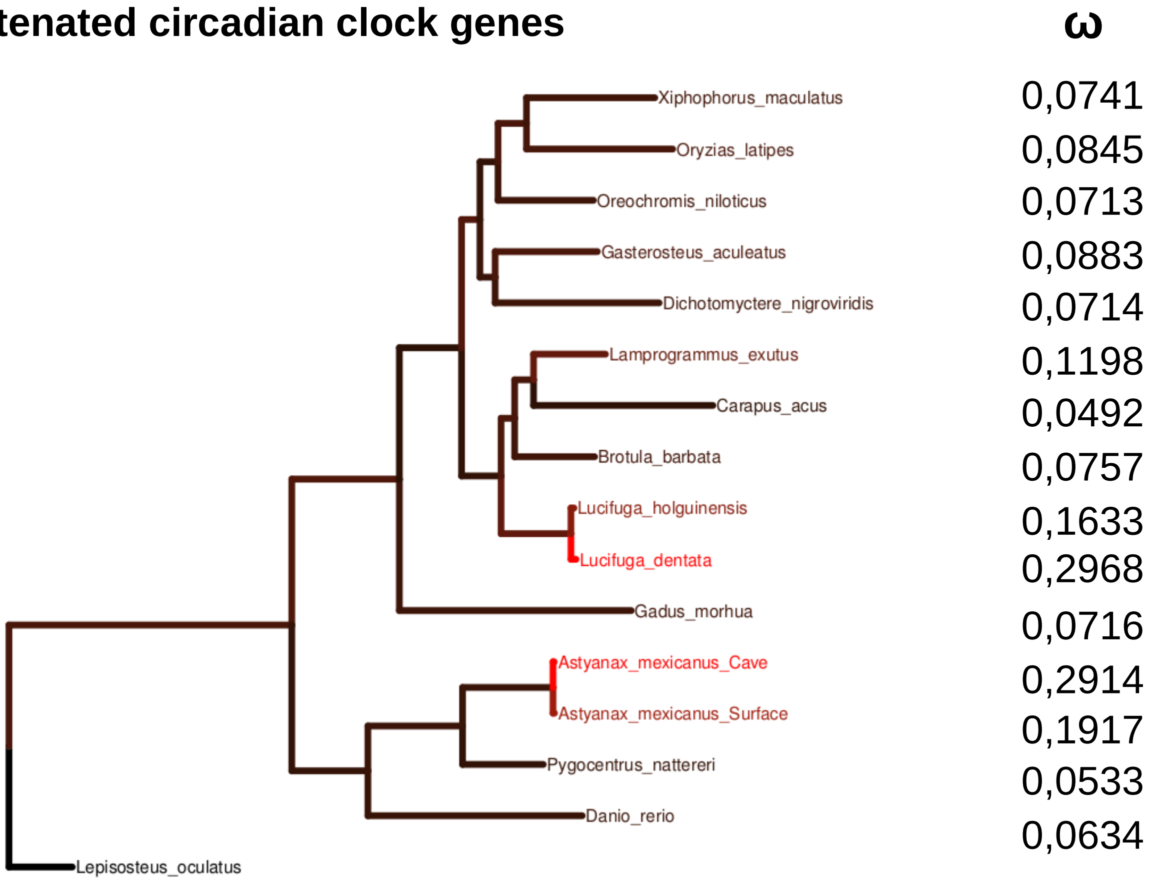

**C**

### Concatenated pigmentation genes

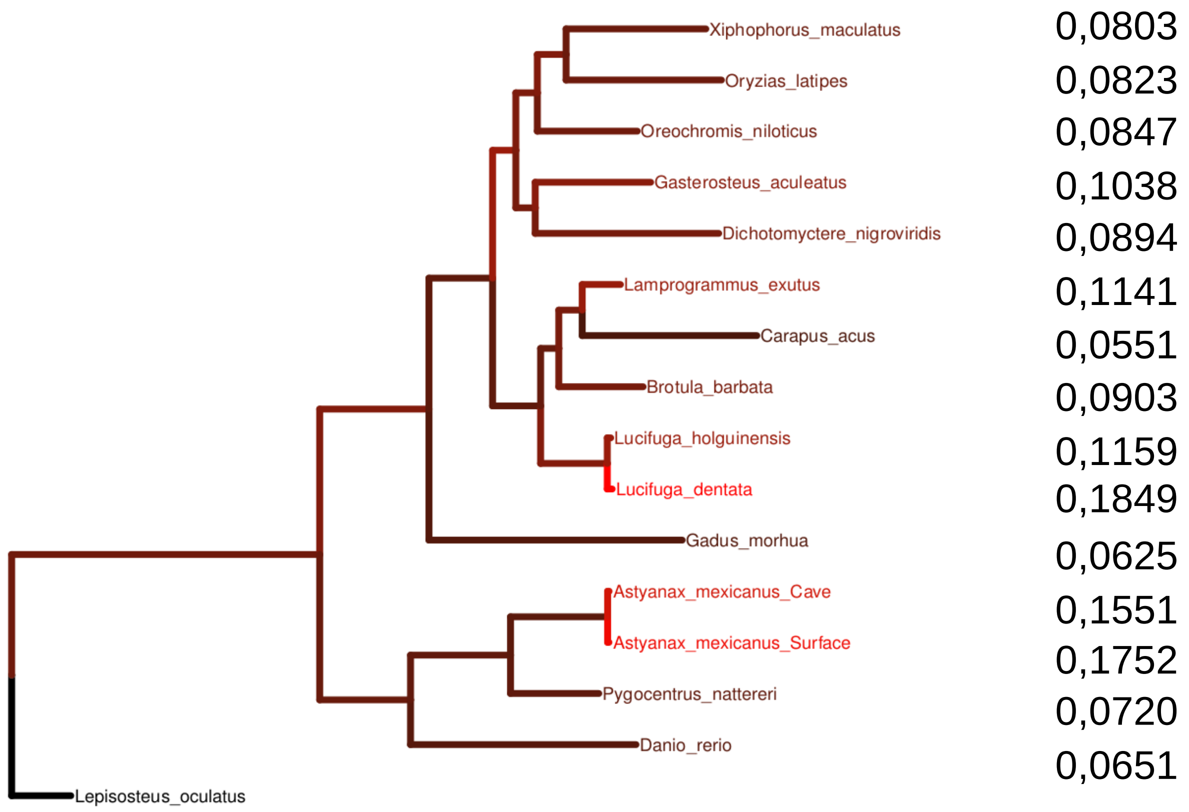


**D**

### Concatenated eye genes - Sinocyclocheilus

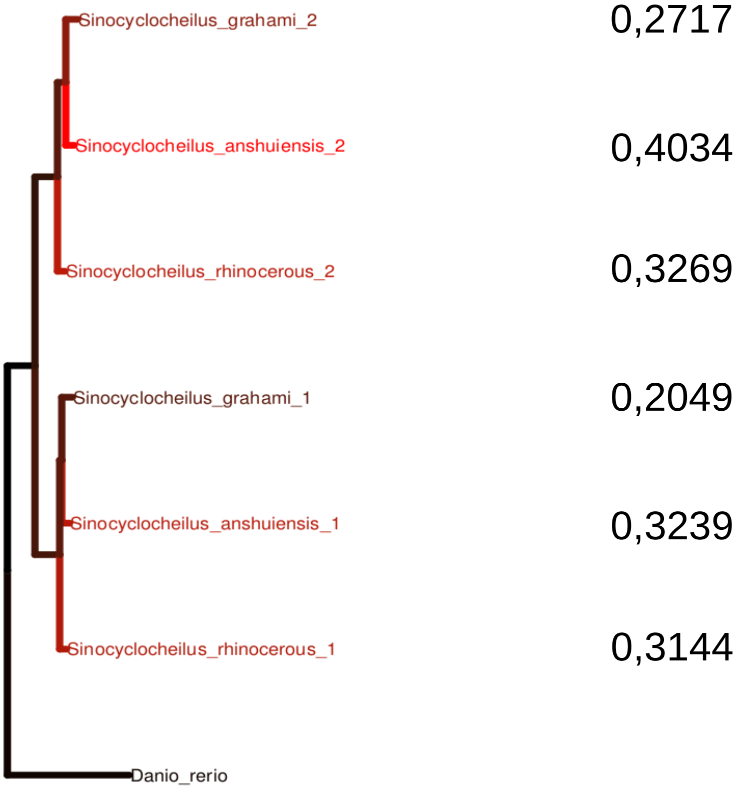

**E**

**Concatenated circadian clock genes - Sinocyclocheilus**

**ω**

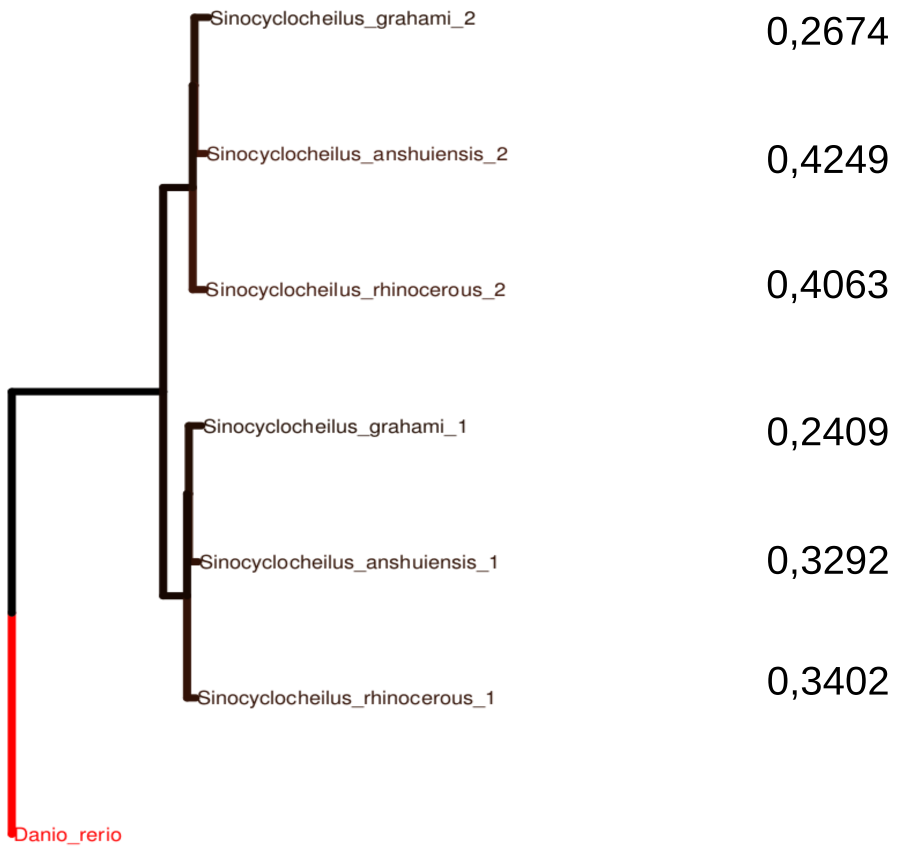


**F**

**Concatenated pigmentation genes - Sinocyclocheilus**

**ω**

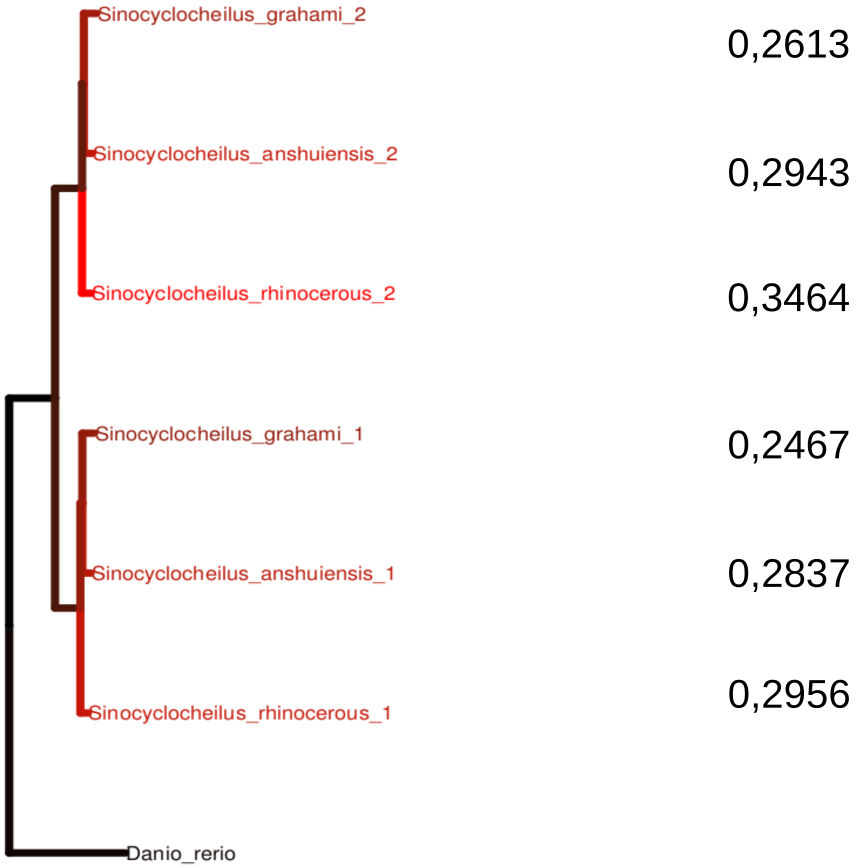
