## Supplementary fig. S14 for "Contrasted gene decay in subterranean vertebrates: insights from cavefishes and fossorial mammals"

### Astyanax mexicanus SF

Test for selection **relaxation** (K = **0.69**) was **not significant** (p = **0.053**, LR = **3.75**).

See here for more information about this method.

Please cite PMID 123456789 if you use this result in a publication, presentation, or other scientific work.

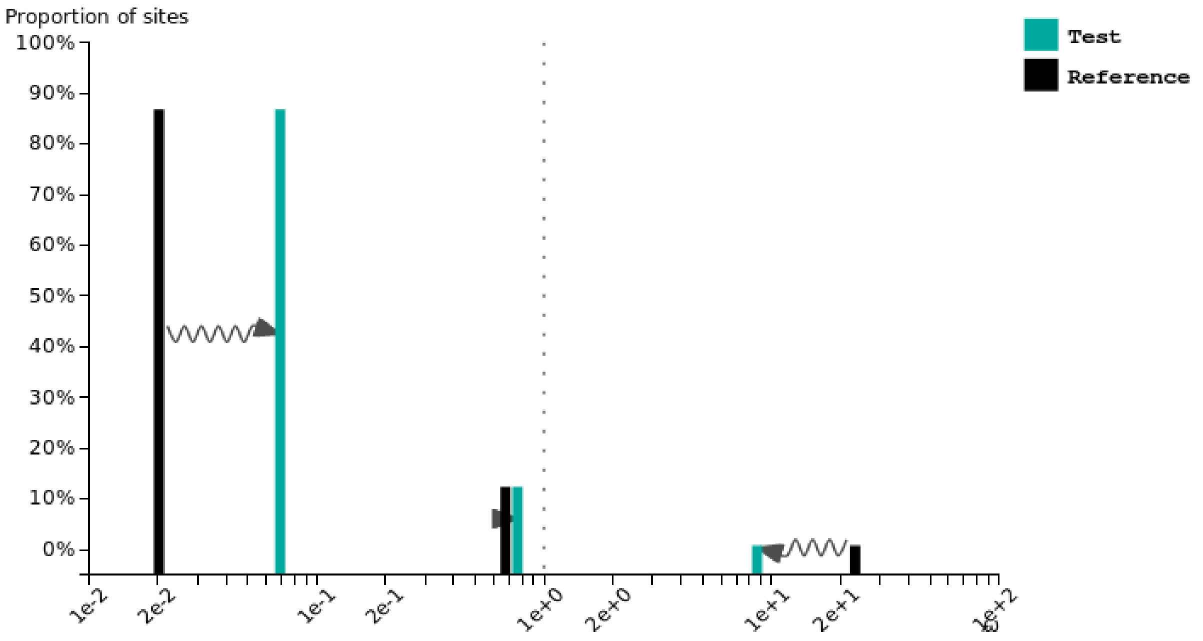


### Astyanax mexicanus CF

Test for selection **relaxation** (K = **0.50**) was **significant** (p = **0.021**, LR = **5.37**).

See here for more information about this method.

Please cite PMID 123456789 if you use this result in a publication, presentation, or other scientific work.

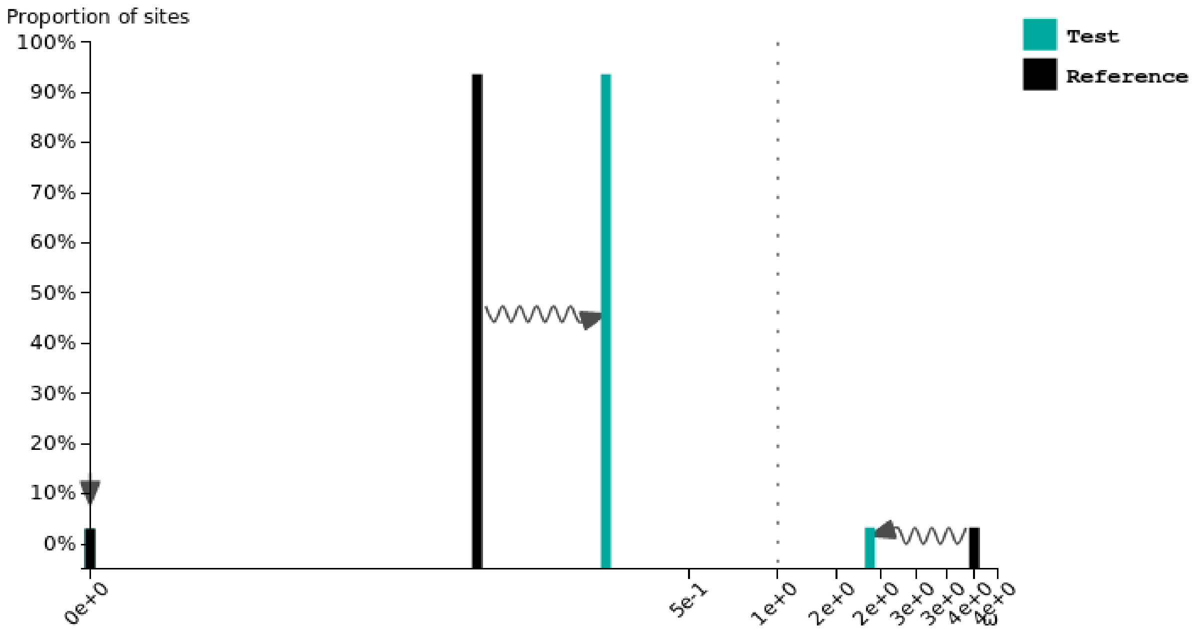

### Brotula barbata

Test for selection **relaxation** (K = **0.91**) was **significant** (p = **0.003**, LR = **8.82**).

See here for more information about this method.

Please cite PMID 123456789 if you use this result in a publication, presentation, or other scientific work.

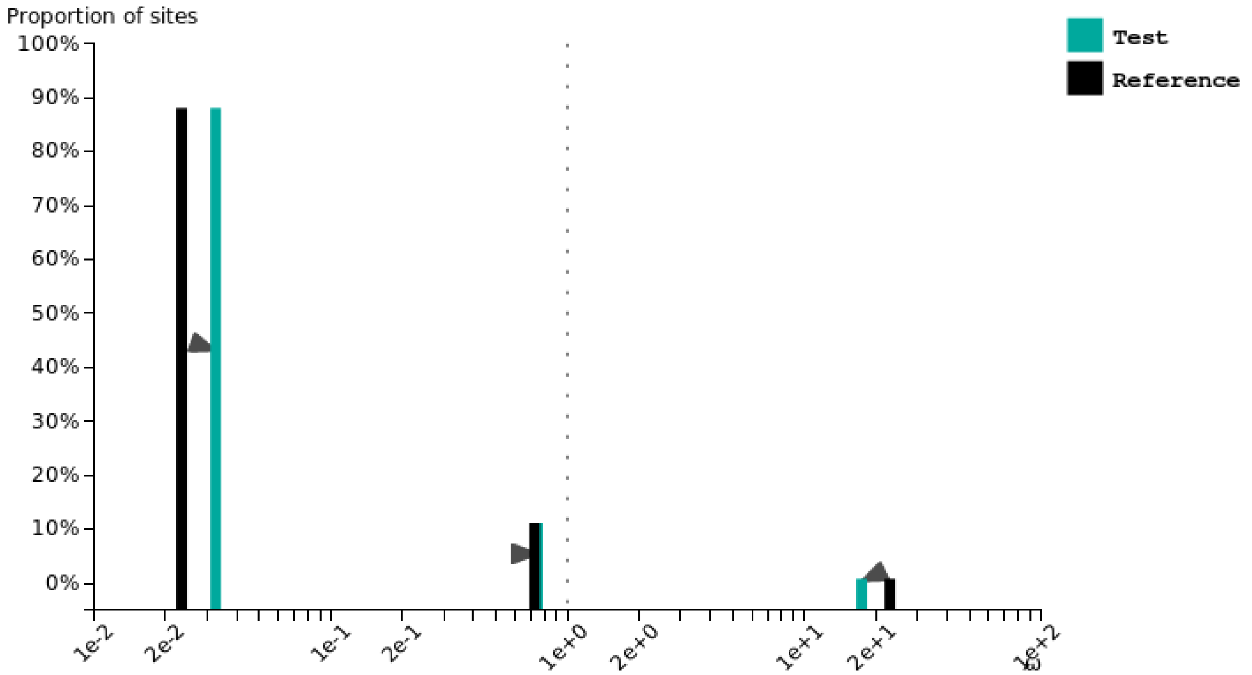


### Carapus acus

Test for selection **intensification** (K = **1.06**) was **significant** (p = **0.030**, LR = **4.73**).

See here for more information about this method.

Please cite PMID 123456789 if you use this result in a publication, presentation, or other scientific work.

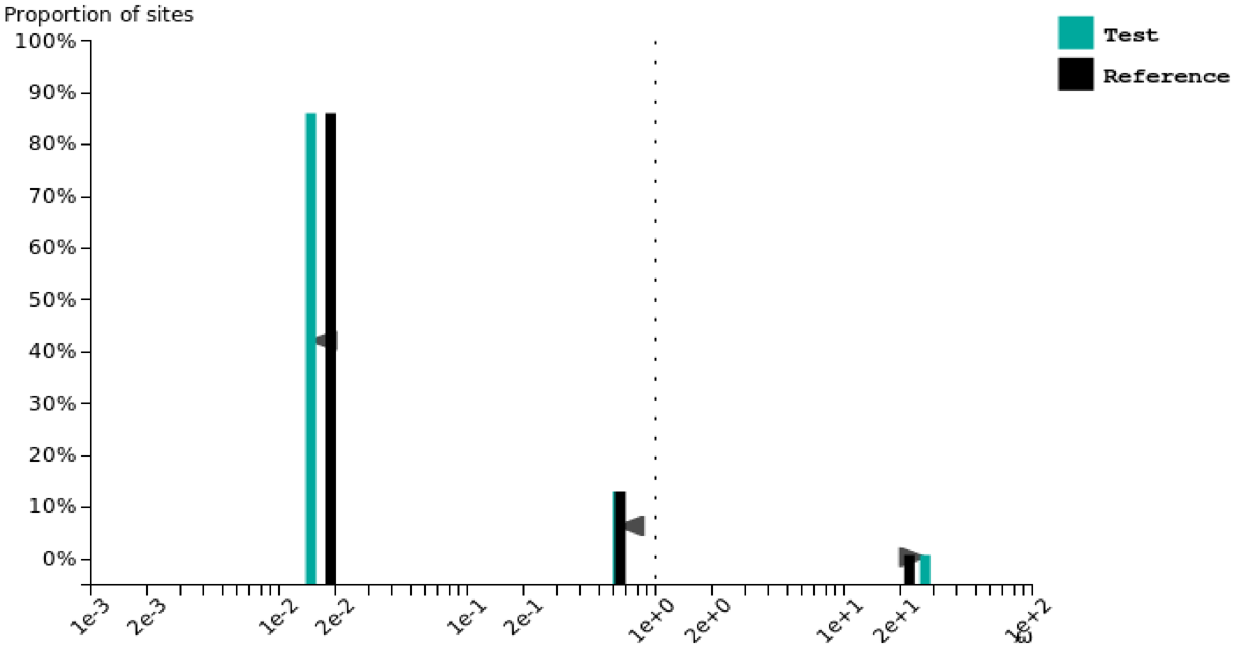

### Lamprogrammus exutus

Test for selection **relaxation** (K = **0.88**) was **significant** (p = **0.004**, LR = **8.49**).

See here for more information about this method.

Please cite PMID 123456789 if you use this result in a publication, presentation, or other scientific work.

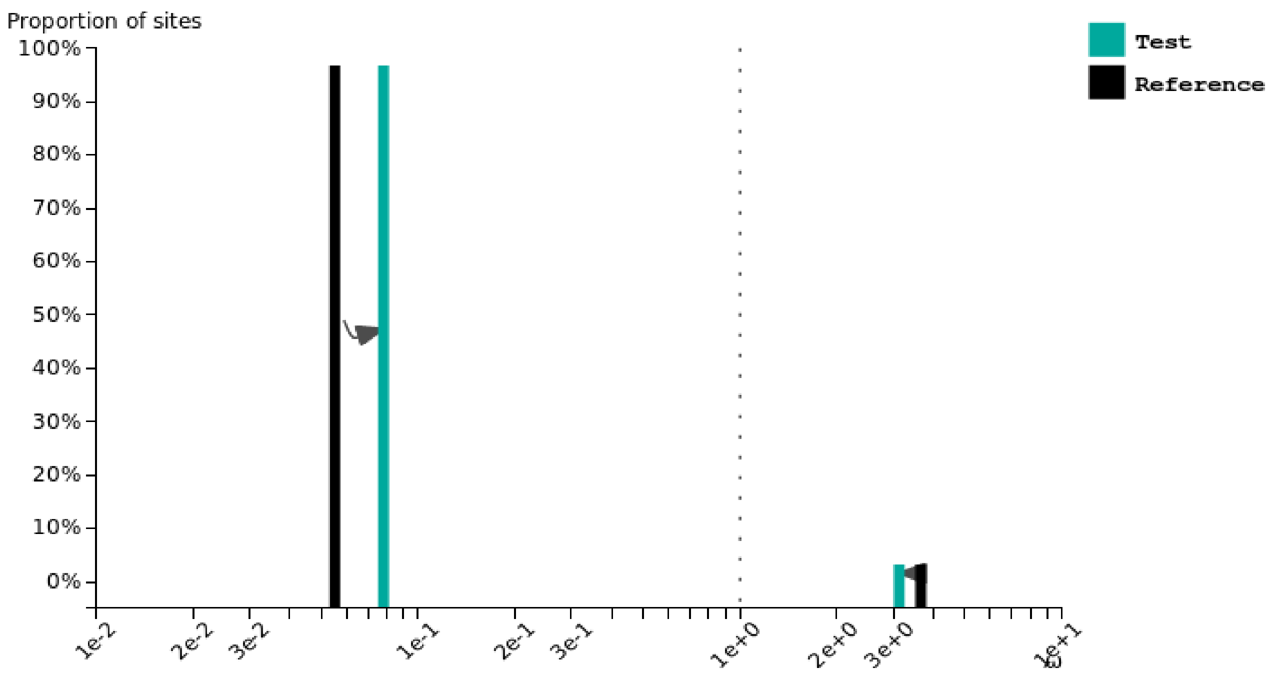


### Lucifuga gibarensis

Test for selection **intensification** (K = **1.63**) was **significant** (p = **0.004**, LR = **8.34**).

See here for more information about this method.

Please cite PMID 123456789 if you use this result in a publication, presentation, or other scientific work.

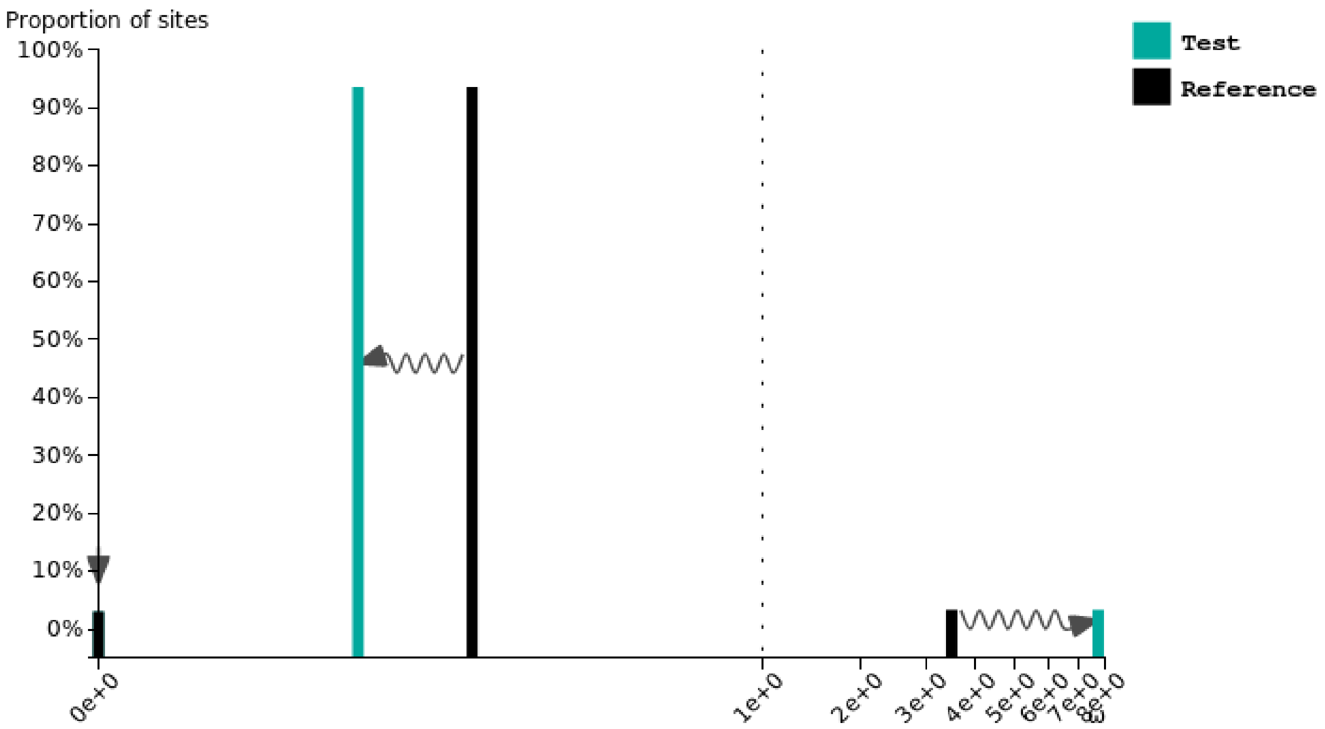

### Lucifuga dentata

Test for selection **relaxation** (K = **0.20**) was **significant** (p = **0.000**, LR = **82.56**).

See here for more information about this method.

Please cite PMID 123456789 if you use this result in a publication, presentation, or other scientific work.

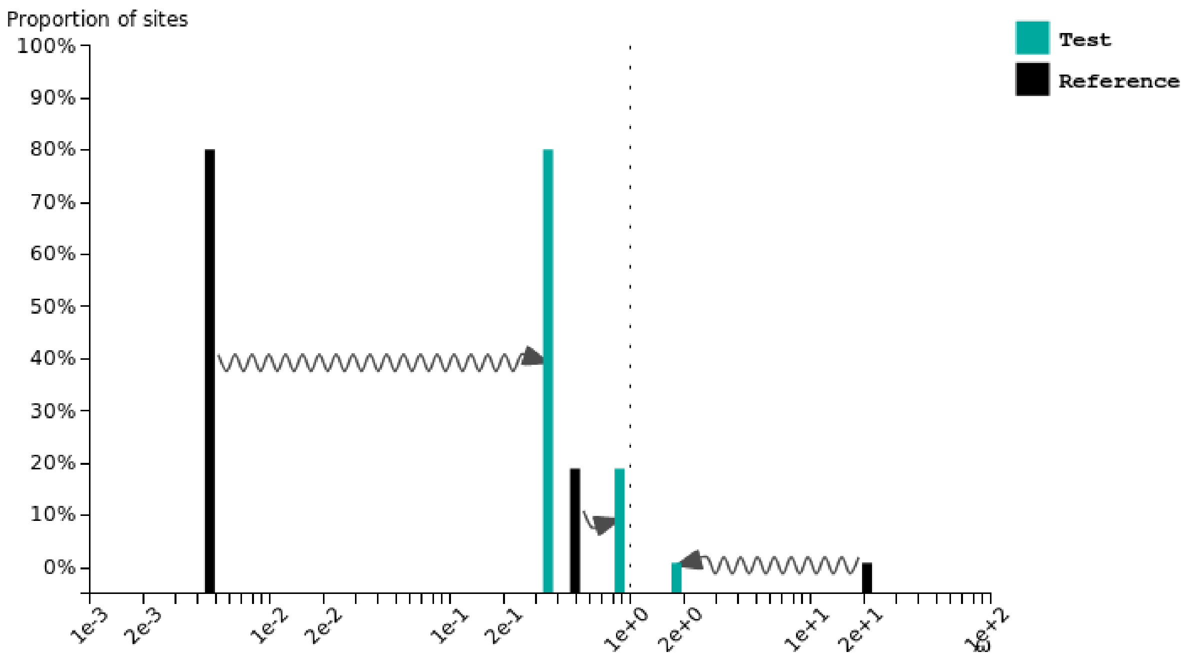


### Danio rerio

Test for selection **intensification** (K = **1.02**) was **not significant** (p = **0.173**, LR = **1.86**).

See here for more information about this method.

Please cite PMID 123456789 if you use this result in a publication, presentation, or other scientific work.

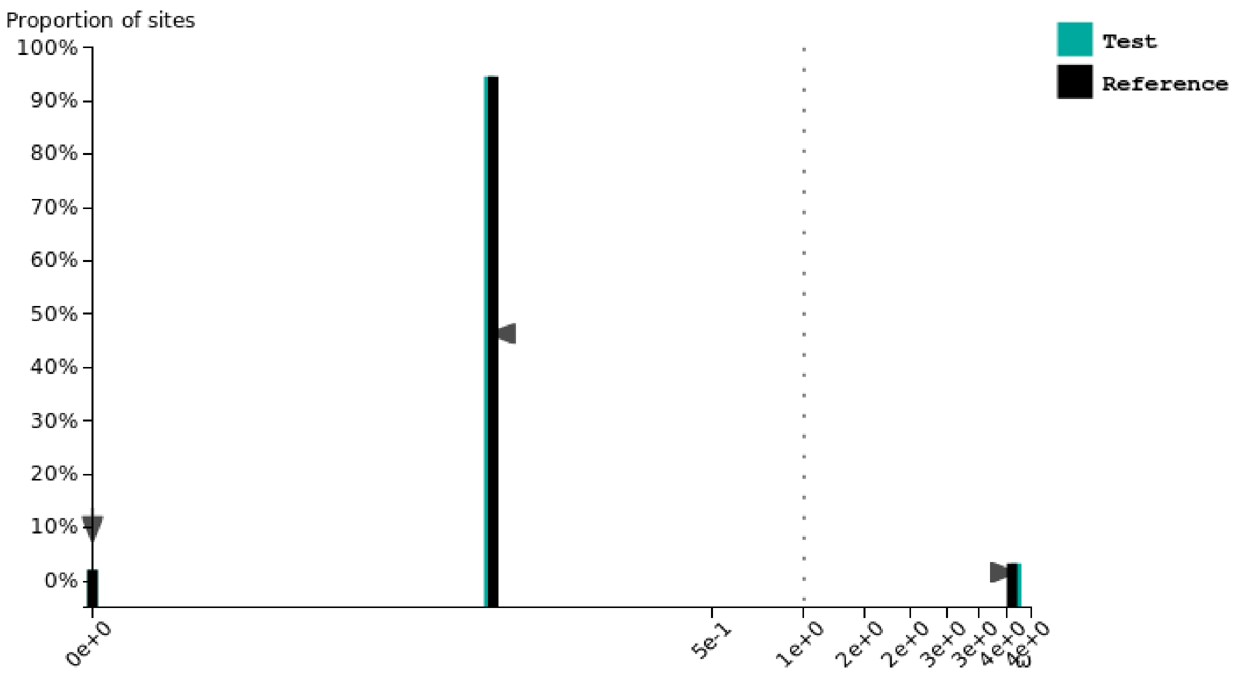

### Dichotomyctere nigroviridis

Test for selection **intensification** (K = **1.03**) was **not significant** (p = **0.429**, LR = **0.63**).

See here for more information about this method.

Please cite PMID 123456789 if you use this result in a publication, presentation, or other scientific work.

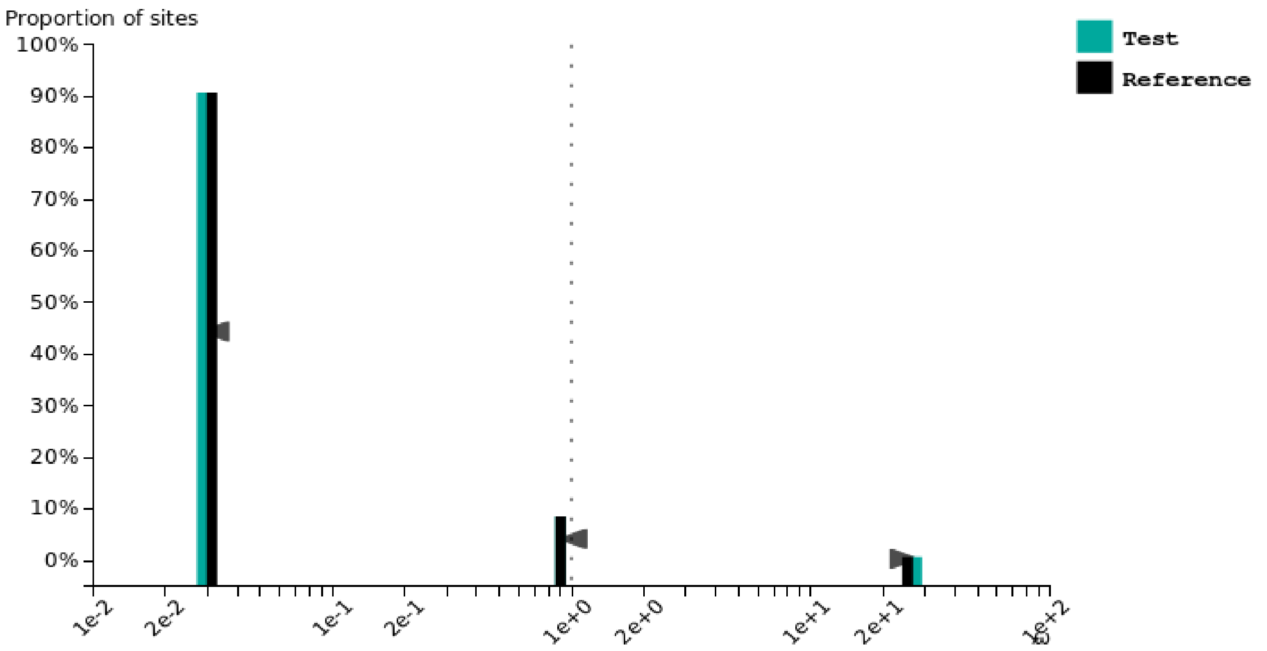


### Gadus morhua

Test for selection **relaxation** (K = **0.94**) was **significant** (p = **0.002**, LR = **9.54**).

See here for more information about this method.

Please cite PMID 123456789 if you use this result in a publication, presentation, or other scientific work.

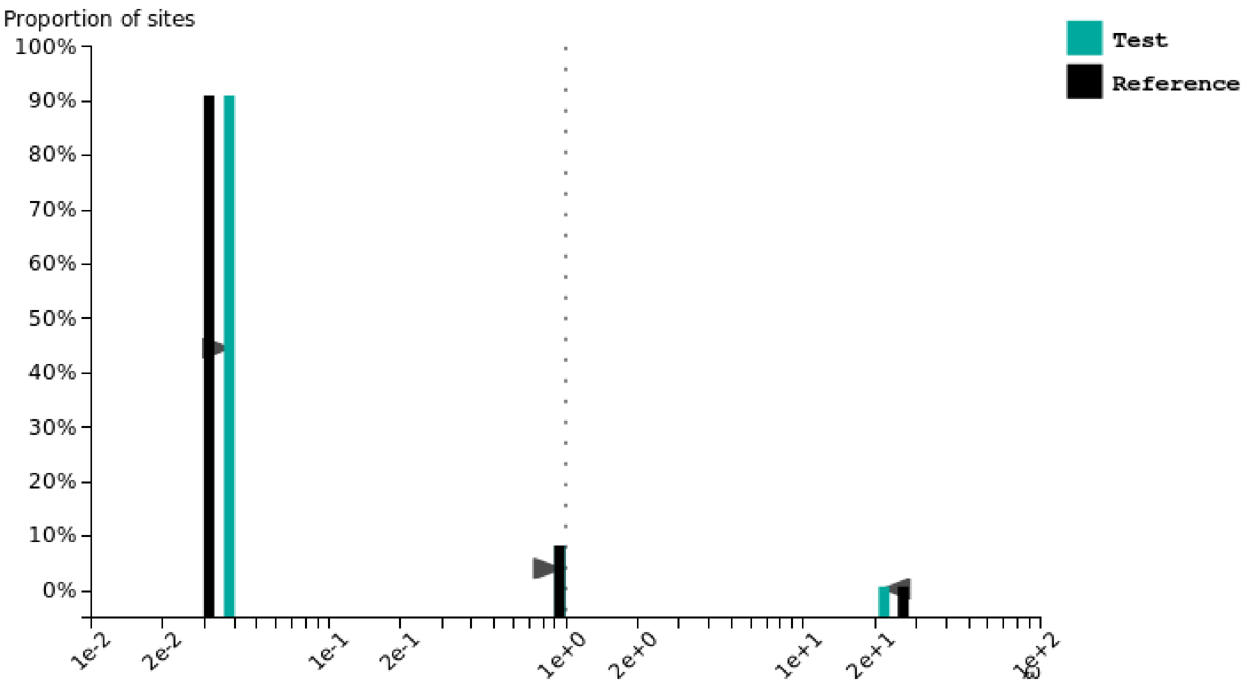

### Gasterosteus aculeatus

Test for selection **relaxation** (K = **0.80**) was **significant** (p = **0.000**, LR = **25.30**).

See here for more information about this method.

Please cite PMID 123456789 if you use this result in a publication, presentation, or other scientific work.

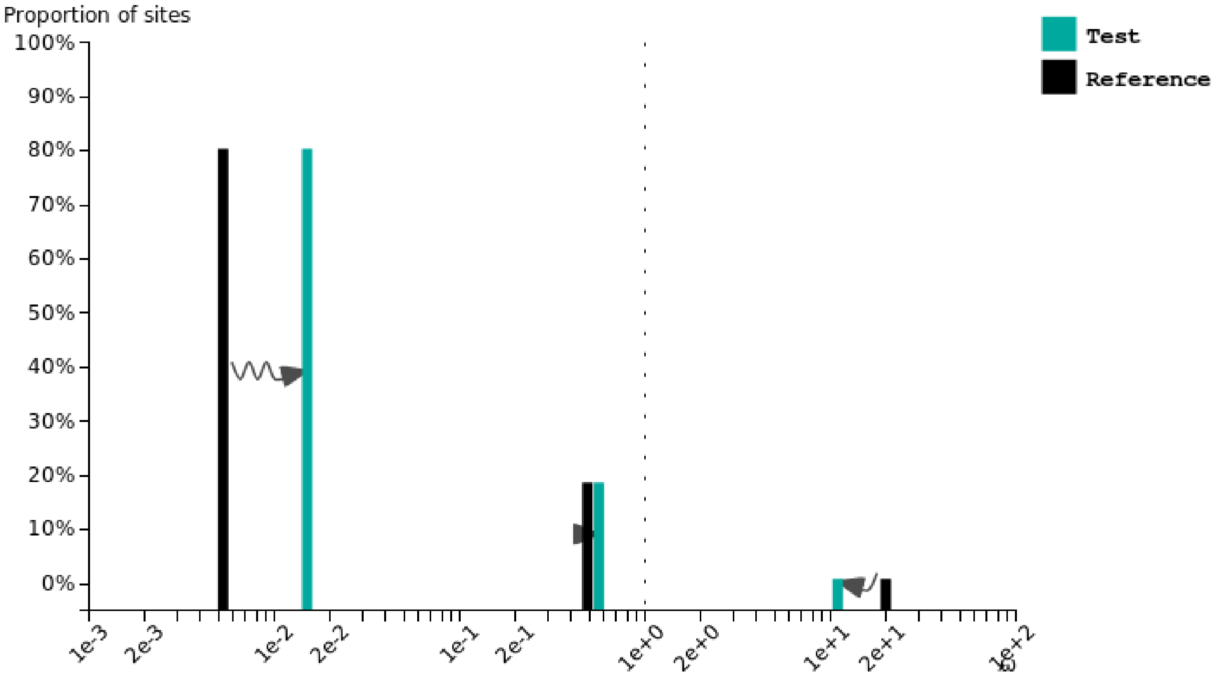


### Oreochromis niloticus

Test for selection **intensification** (K = **1.04**) was **not significant** (p = **0.093**, LR = **2.82**).

See here for more information about this method.

Please cite PMID 123456789 if you use this result in a publication, presentation, or other scientific work.

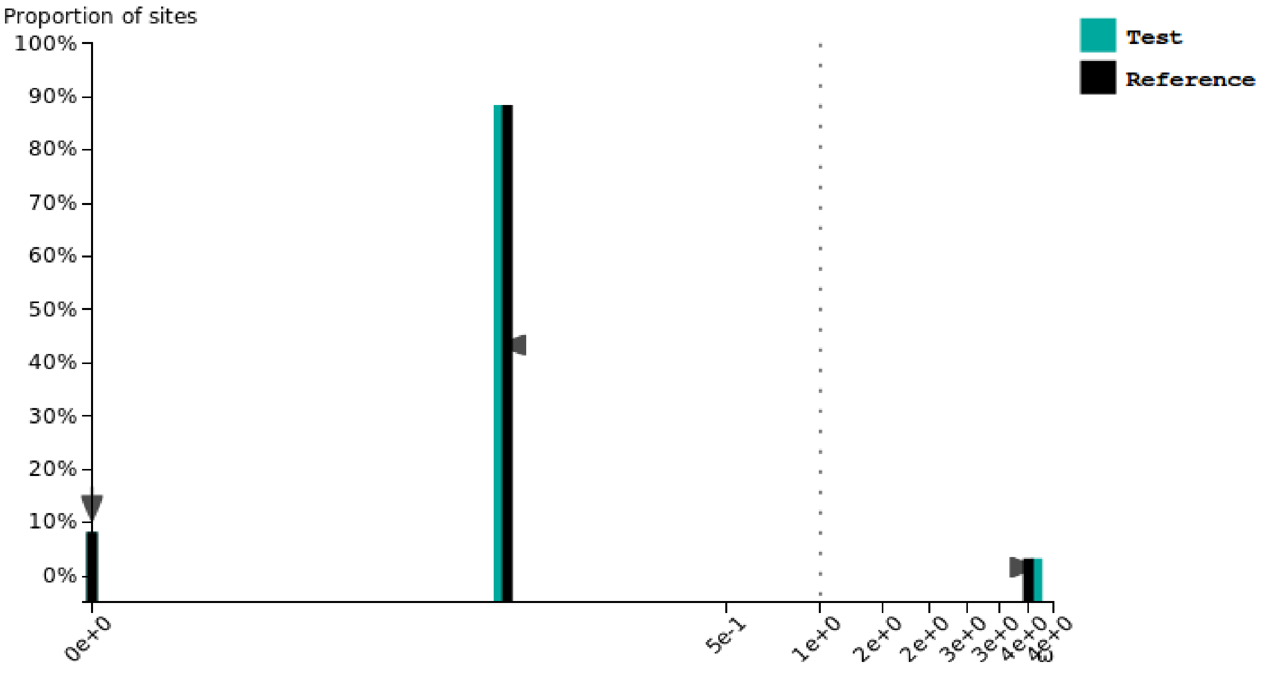

### Oryzias latipes

Test for selection **intensification** (K = **1.27**) was **significant** (p = **0.000**, LR = **35.13**).

See here for more information about this method.
Please cite PMID 123456789 if you use this result in a publication, presentation, or other scientific work.

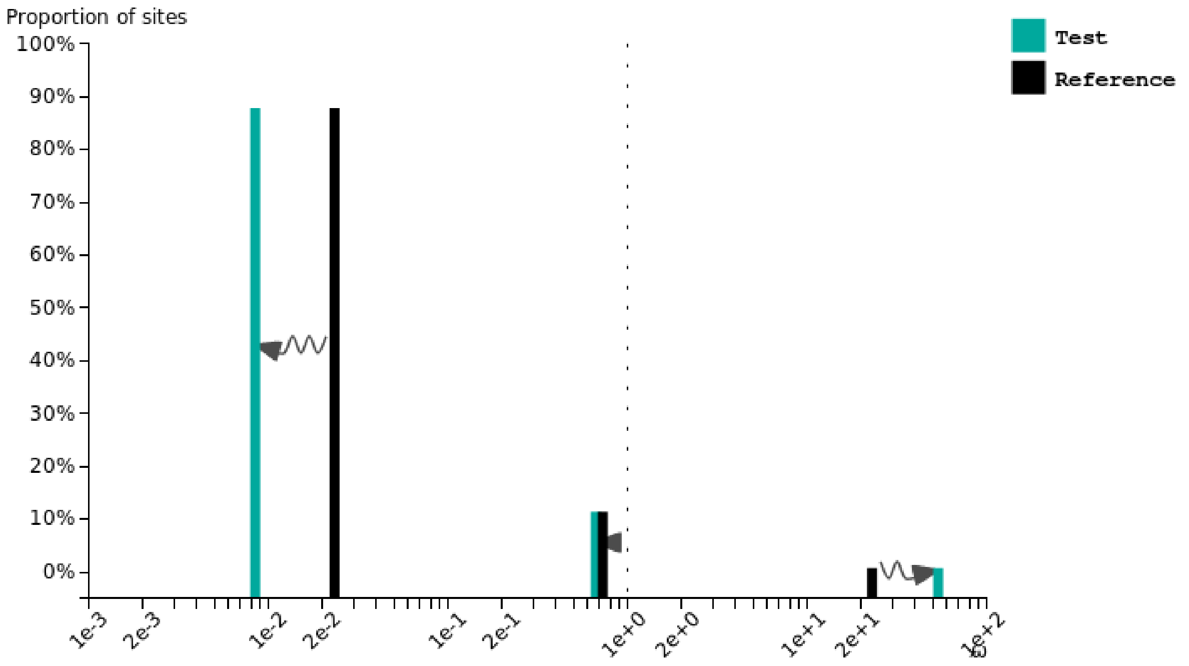


### Xiphophorus maculatus

Test for selection **intensification** (K = **1.06**) was **significant** (p = **0.014**, LR = **6.08**).

See here for more information about this method.
Please cite PMID 123456789 if you use this result in a publication, presentation, or other scientific work.

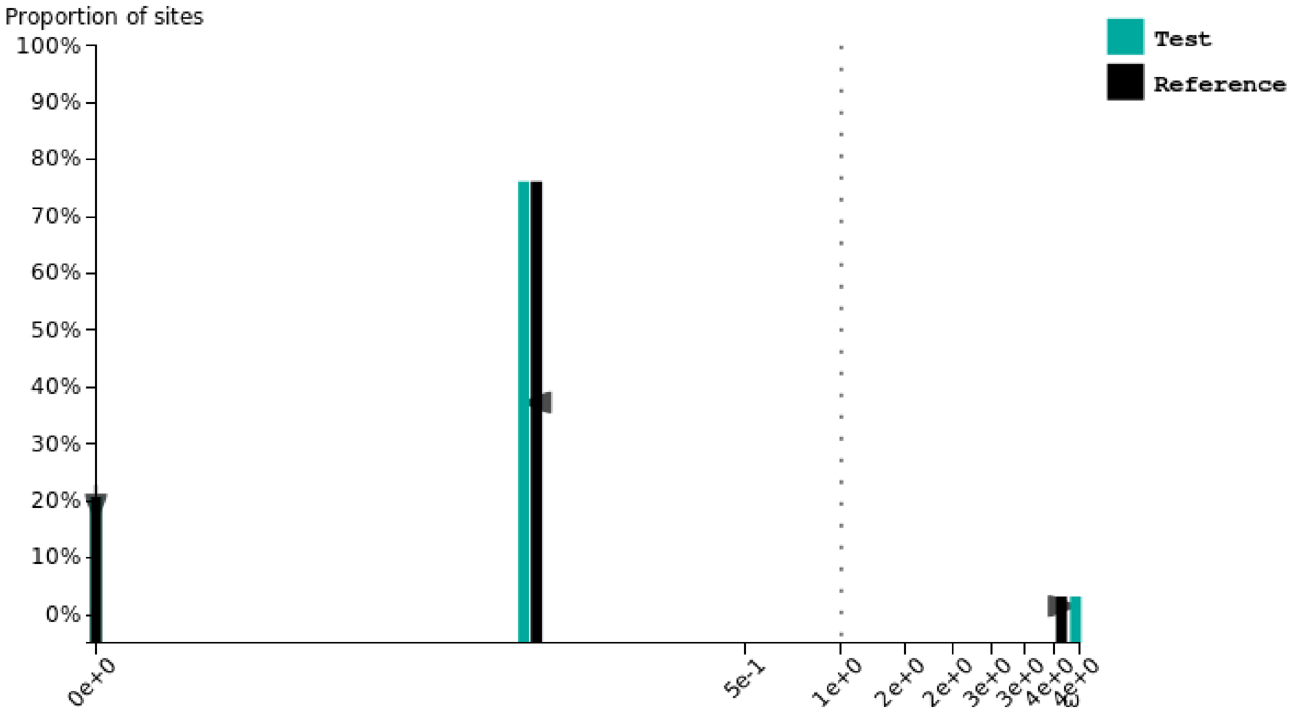

### Pygocentrus nattereri

Test for selection **intensification** (K = **1.04**) was **not significant** (p = **0.661**, LR = **0.19**).

See here for more information about this method.

Please cite PMID 123456789 if you use this result in a publication, presentation, or other scientific work.

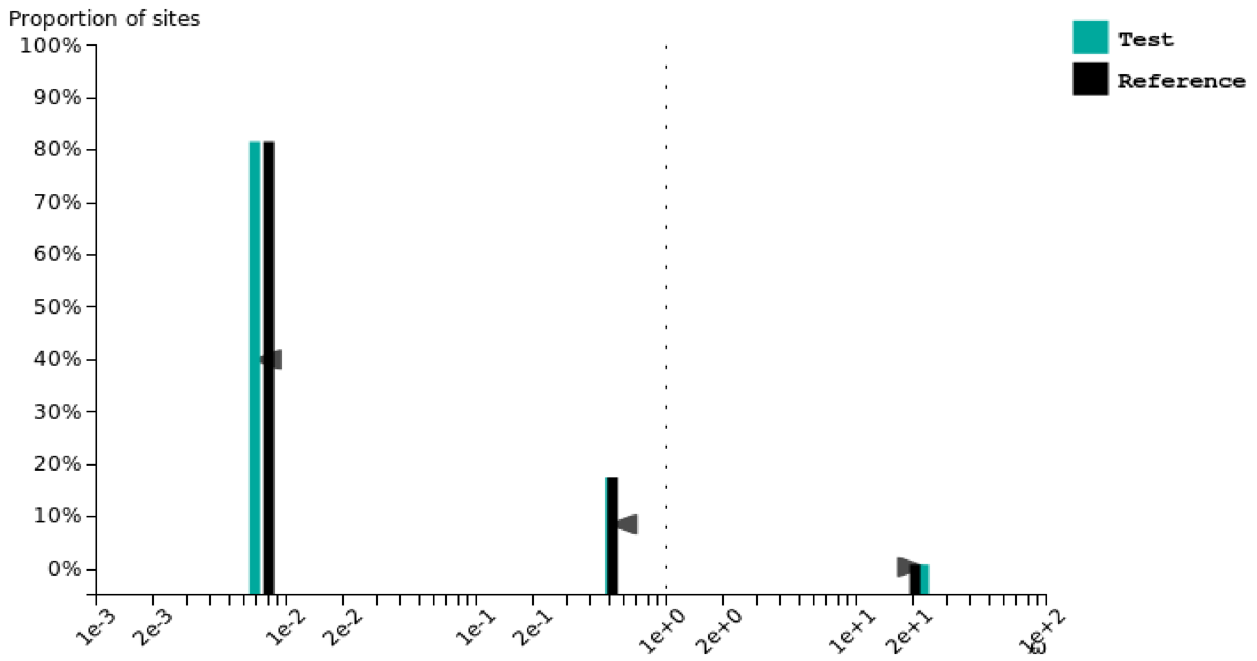
