## Supplementary fig. S15 for "Contrasted gene decay in subterranean vertebrates: insights from cavefishes and fossorial mammals"

### Astyanax mexicanus SF

Test for selection **relaxation** (K = **0.56**) was **not significant** (p = **0.078**, LR = **3.12**).

See here for more information about this method.
Please cite PMID 123456789 if you use this result in a publication, presentation, or other scientific work.

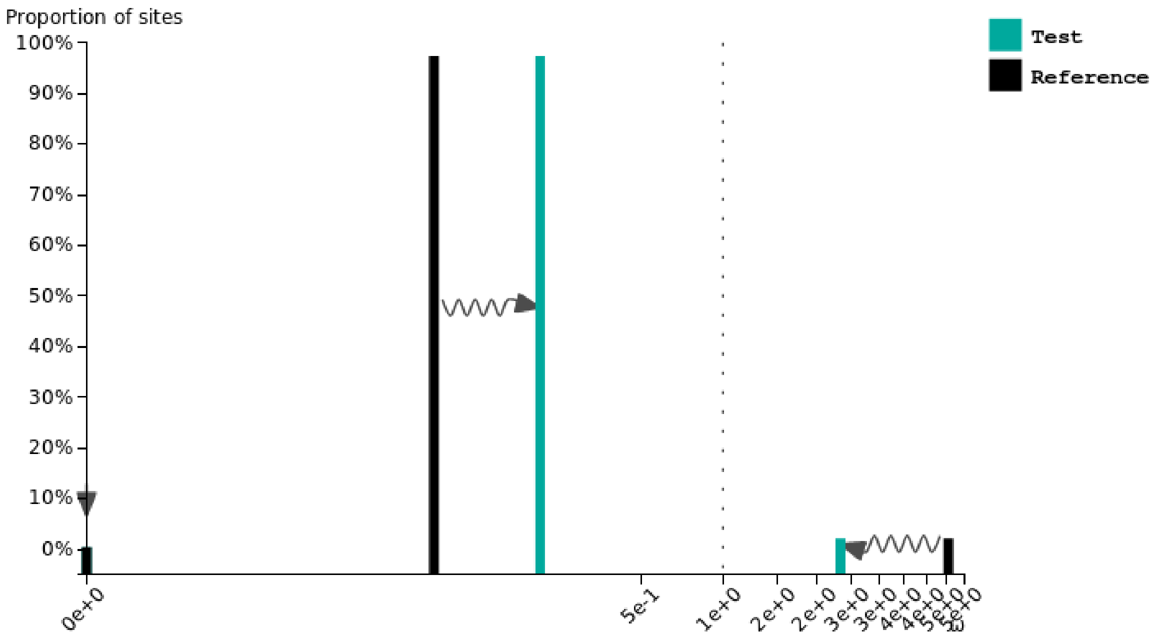


### Astyanax mexicanus CF

Test for selection **intensification** (K = **1.22**) was **not significant** (p = **0.195**, LR = **1.68**).

See here for more information about this method.
Please cite PMID 123456789 if you use this result in a publication, presentation, or other scientific work.

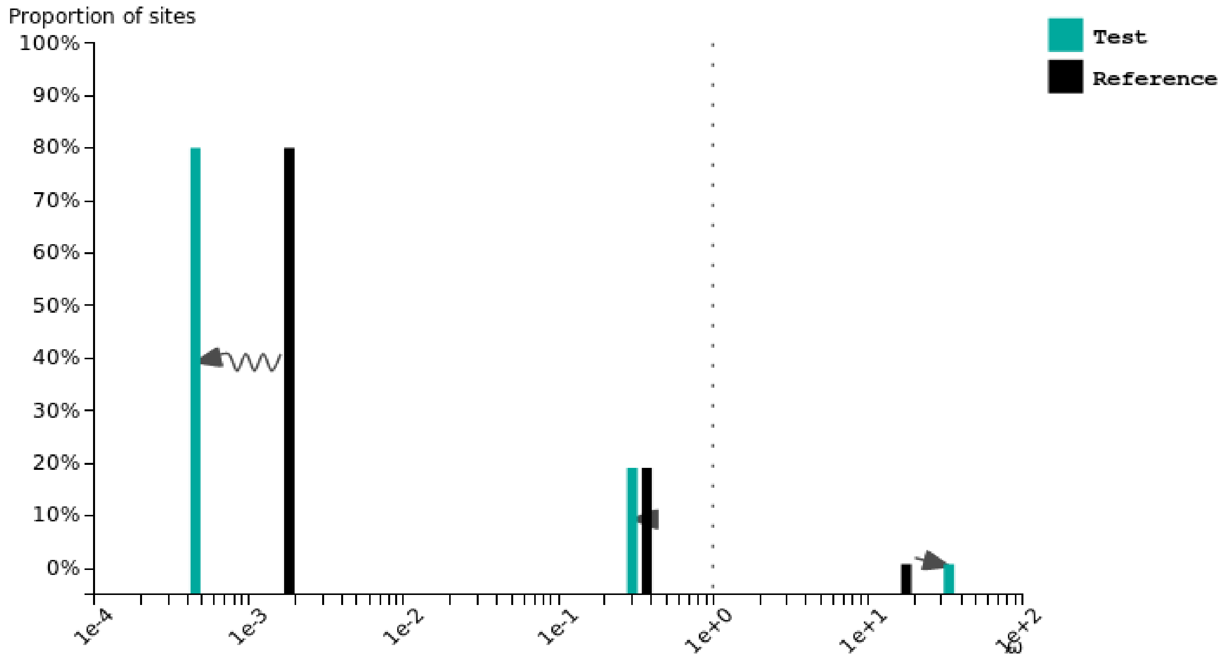

### Brotula barbata

Test for selection **relaxation** (K = **0.97**) was **not significant** (p = **0.234**, LR = **1.42**).

See here for more information about this method.
Please cite PMID 123456789 if you use this result in a publication, presentation, or other scientific work.

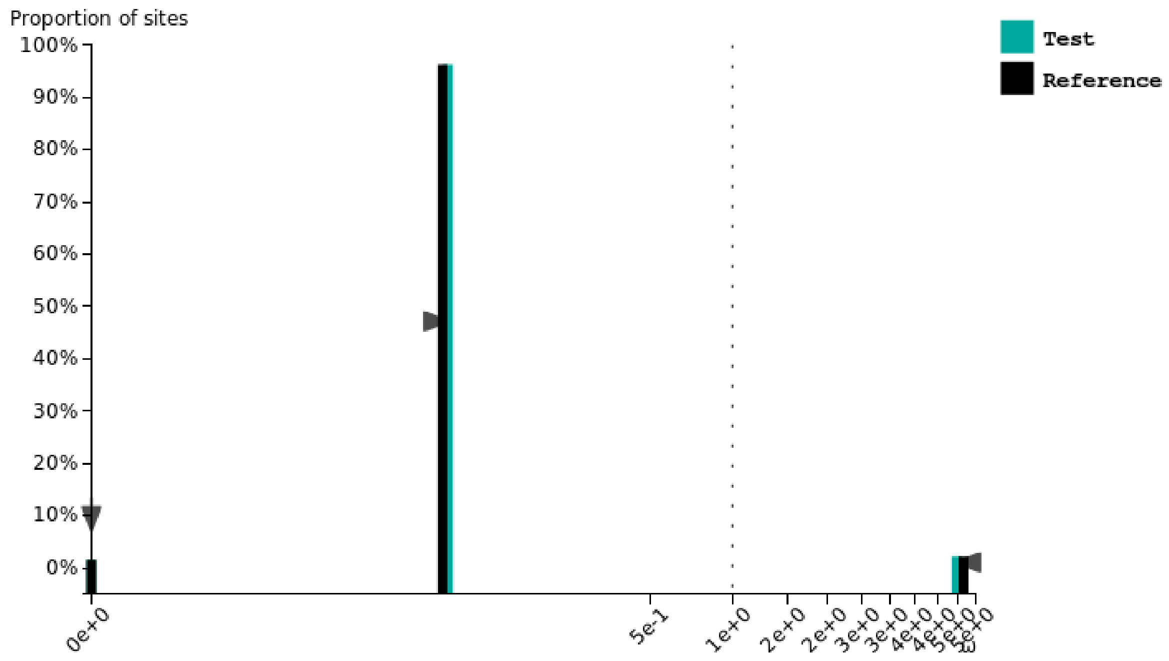


### Carapus acus

Test for selection **intensification** (K = **1.15**) was **significant** (p = **0.000**, LR = **34.13**).

See here for more information about this method.
Please cite PMID 123456789 if you use this result in a publication, presentation, or other scientific work.

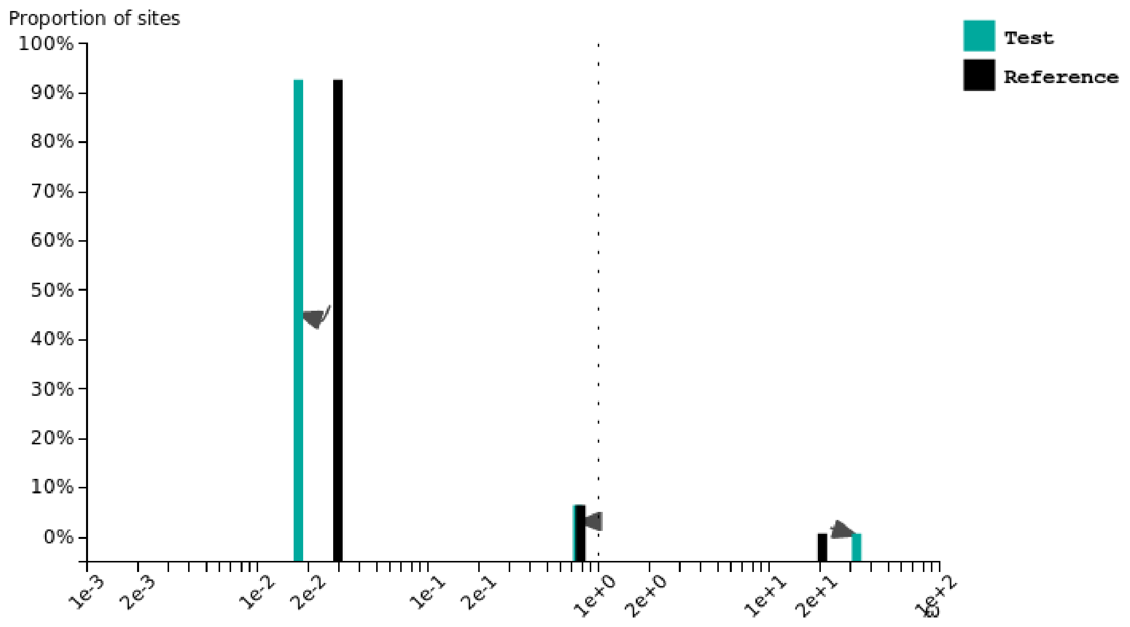

### Lamprogrammus exutus

Test for selection **intensification** (K = **1.03**) was **not significant** (p = **0.228**, LR = **1.45**).

See here for more information about this method.
Please cite PMID 123456789 if you use this result in a publication, presentation, or other scientific work.

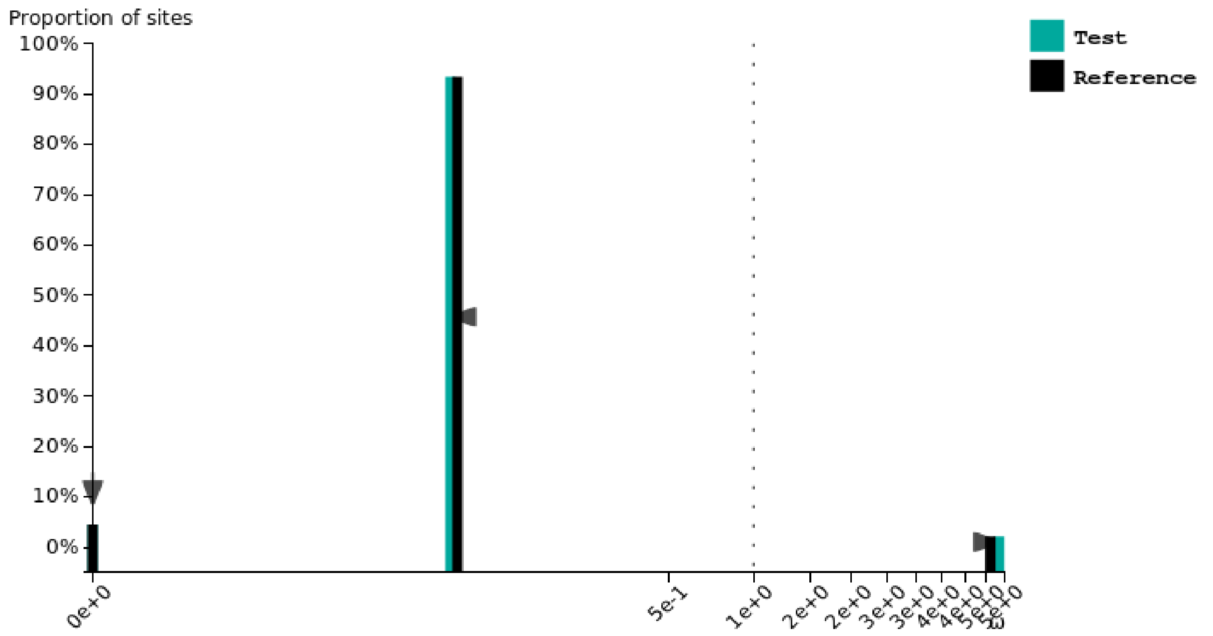


### Lucifuga gibarensis

Test for selection **intensification** (K = **1.00**) was **significant** (p = **0.001**, LR = **10.96**).

See here for more information about this method.
Please cite PMID 123456789 if you use this result in a publication, presentation, or other scientific work.

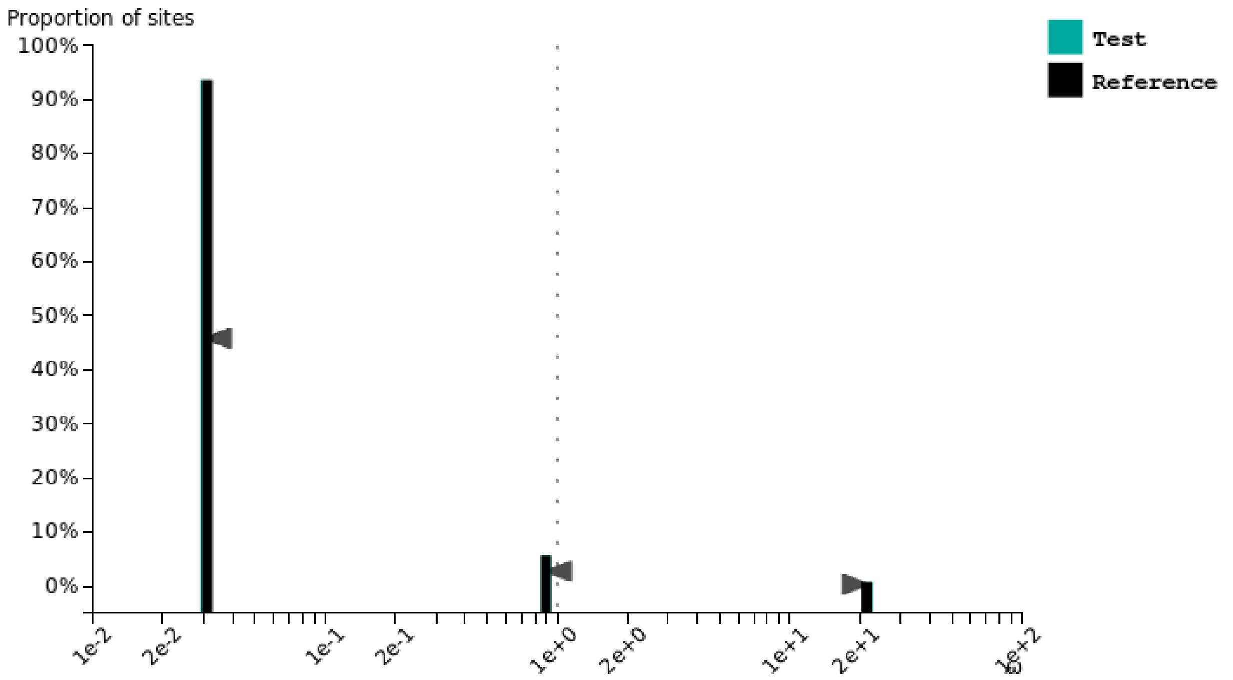

### Lucifuga dentata

Test for selection **intensification** (K = **1.07**) was **not significant** (p = **0.487**, LR = **0.48**).

See here for more information about this method.
Please cite PMID 123456789 if you use this result in a publication, presentation, or other scientific work.

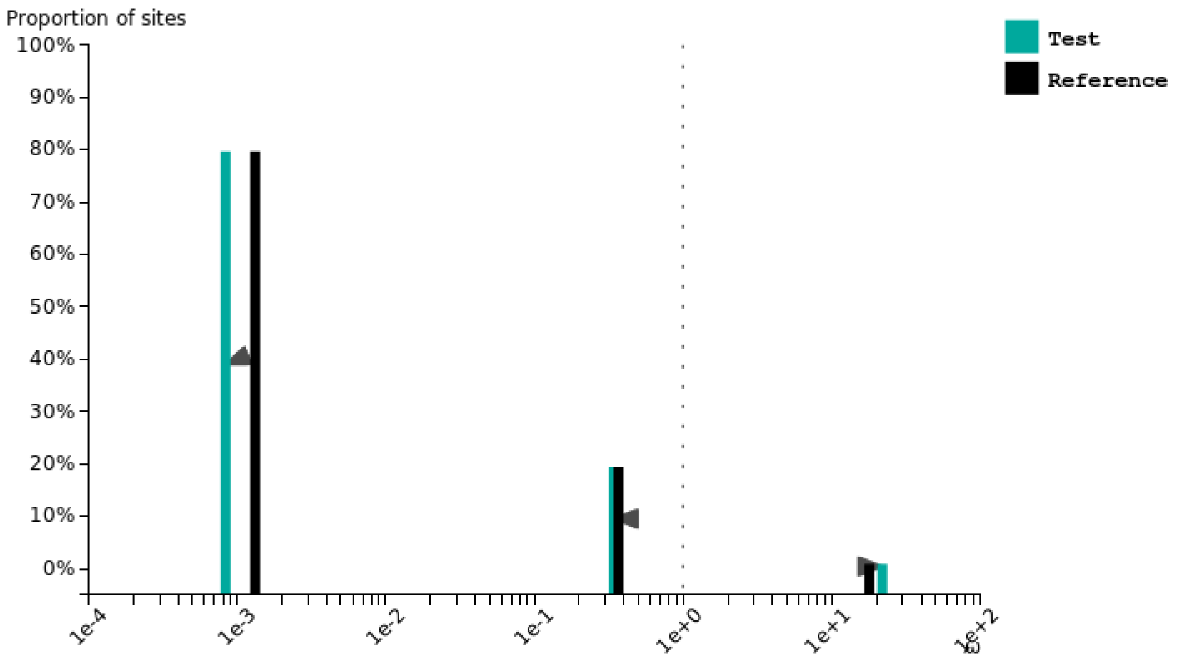


### Danio rerio

Test for selection **relaxation** (K = **0.98**) was **significant** (p = **0.001**, LR = **11.65**).

See here for more information about this method.
Please cite PMID 123456789 if you use this result in a publication, presentation, or other scientific work.

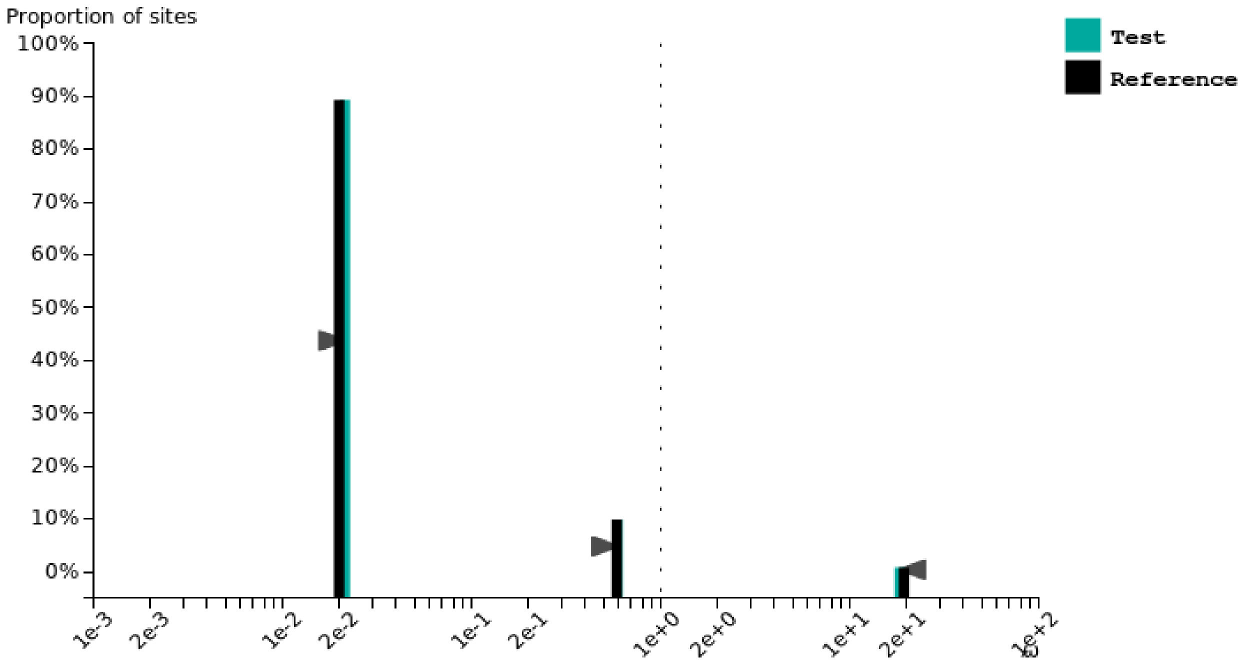

### Dichotomyctere nigroviridis

Test for selection **relaxation** (K = **1.00**) was **not significant** (p = **0.879**, LR = **0.02**).

See here for more information about this method.

Please cite PMID 123456789 if you use this result in a publication, presentation, or other scientific work.

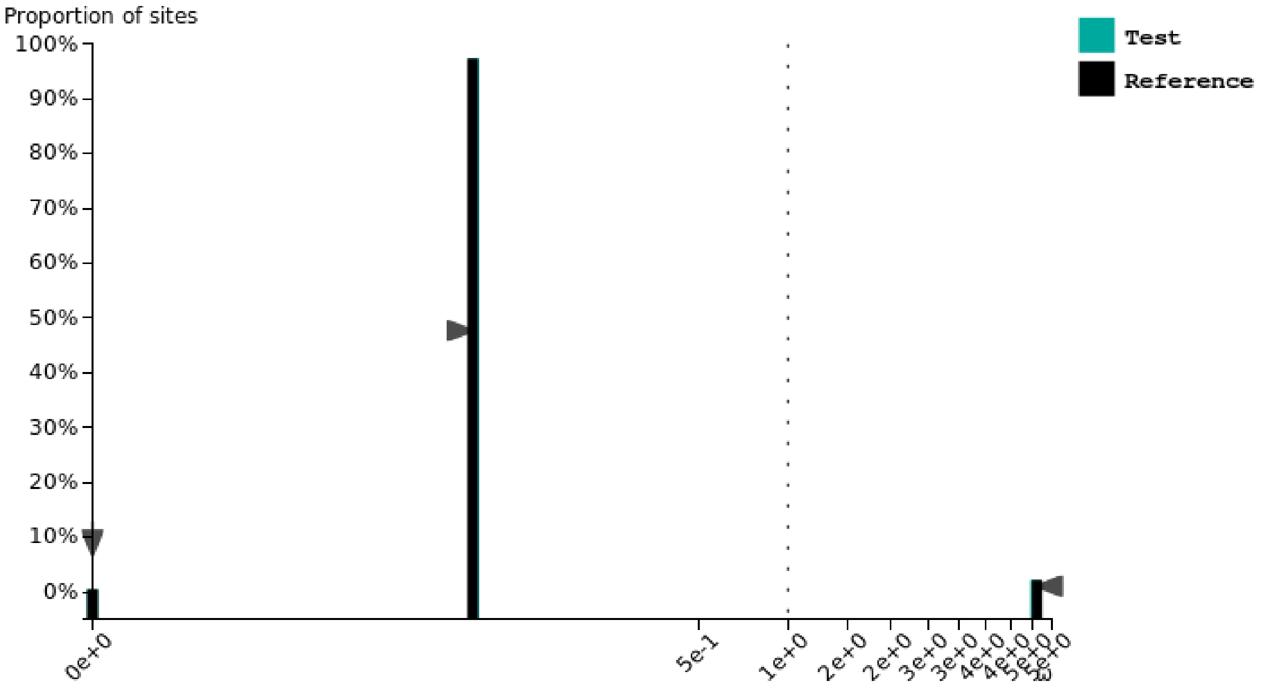


### Gadus morhua

Test for selection **relaxation** (K = **0.95**) was **significant** (p = **0.002**, LR = **9.98**).

See here for more information about this method.

Please cite PMID 123456789 if you use this result in a publication, presentation, or other scientific work.

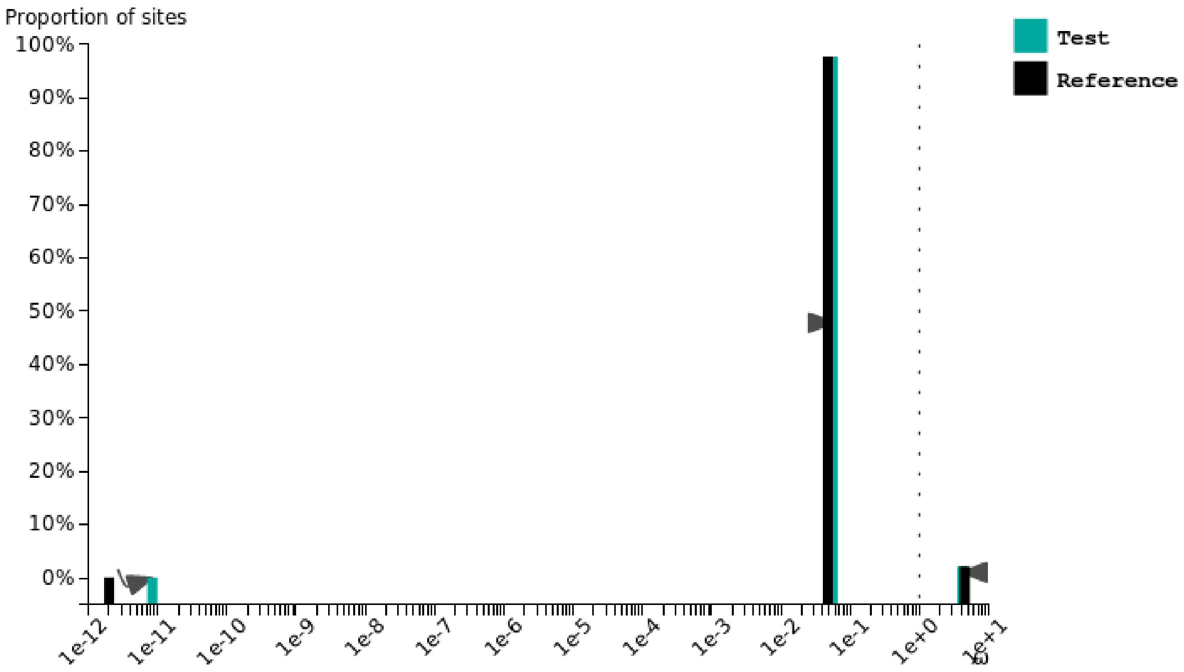

### Gasterosteus aculeatus

Test for selection **relaxation** (K = **0.90**) was **significant** (p = **0.000**, LR = **21.90**).

See here for more information about this method.

Please cite PMID 123456789 if you use this result in a publication, presentation, or other scientific work.

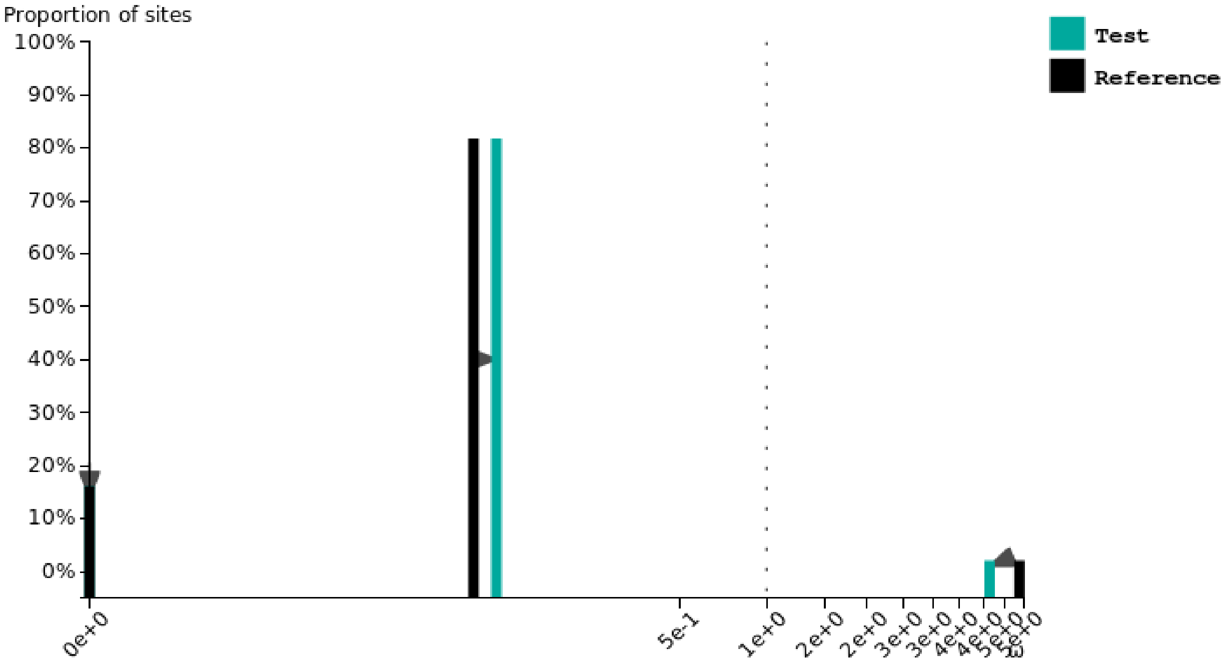


### Oreochromis niloticus

Test for selection **intensification** (K = **1.02**) was **not significant** (p = **0.334**, LR = **0.93**).

See here for more information about this method.

Please cite PMID 123456789 if you use this result in a publication, presentation, or other scientific work.

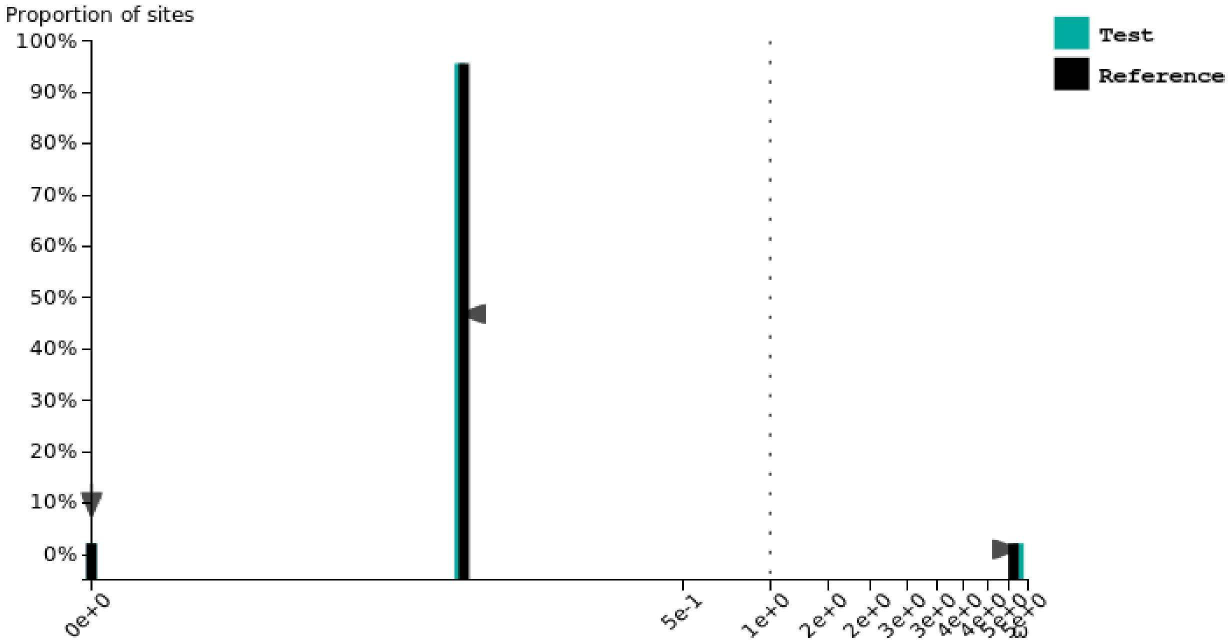

### Oryzias latipes

Test for selection **relaxation** (K = **0.91**) was **significant** (p = **0.000**, LR = **28.24**).

See here for more information about this method.
Please cite PMID 123456789 if you use this result in a publication, presentation, or other scientific work.

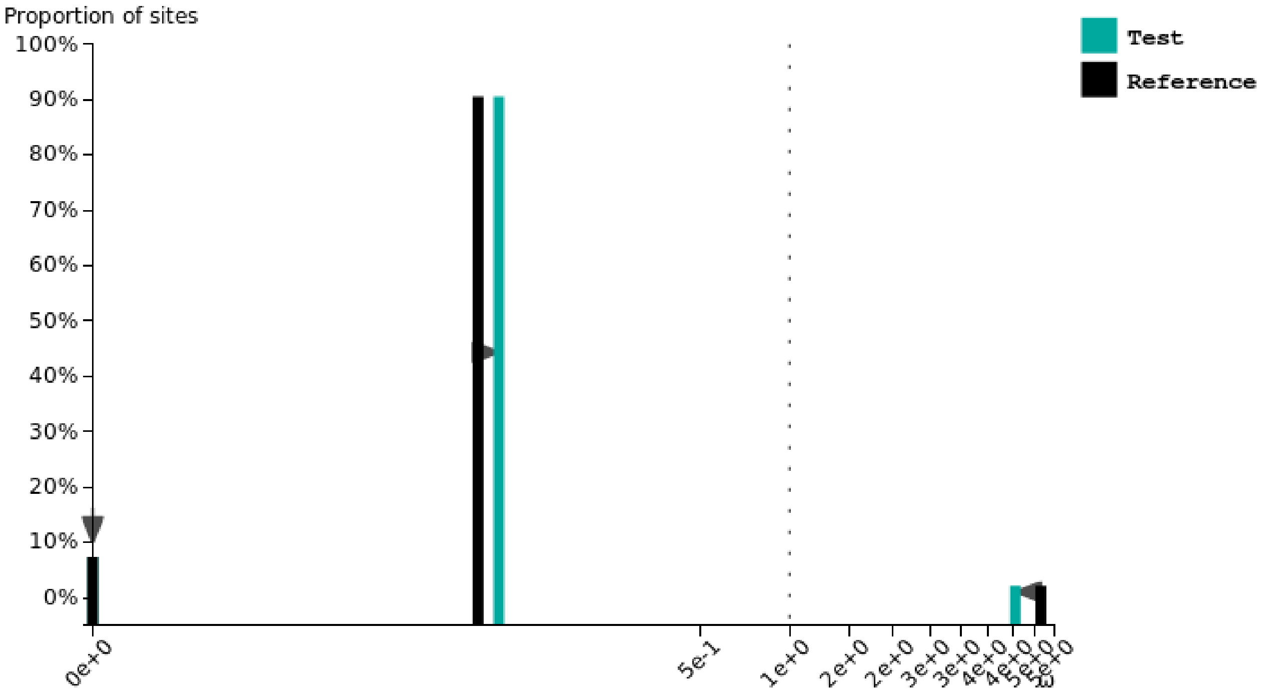


### Xiphophorus maculatus

Test for selection **relaxation** (K = **0.92**) was **significant** (p = **0.030**, LR = **4.69**).

See here for more information about this method.
Please cite PMID 123456789 if you use this result in a publication, presentation, or other scientific work.

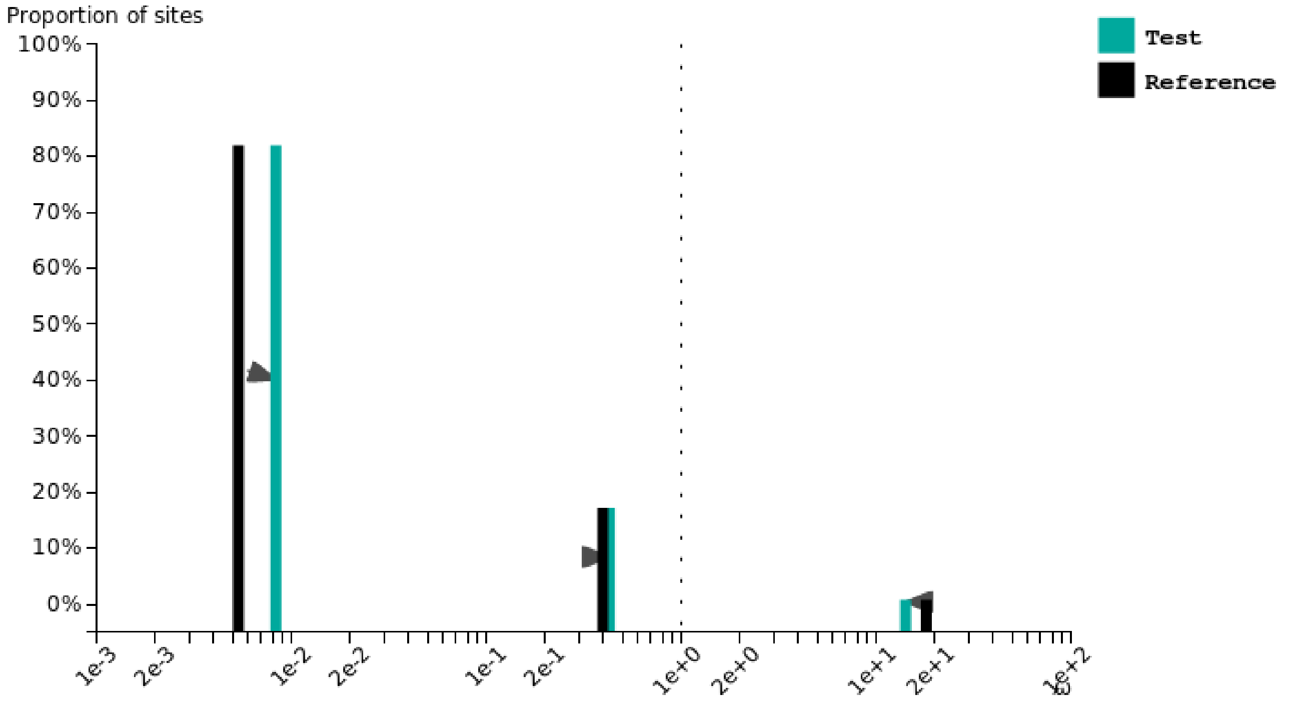

### Pygocentrus nattereri

Test for selection **intensification** (K = **1.10**) was **significant** (p = **0.000**, LR = **13.84**).

See here for more information about this method.

Please cite PMID 123456789 if you use this result in a publication, presentation, or other scientific work.

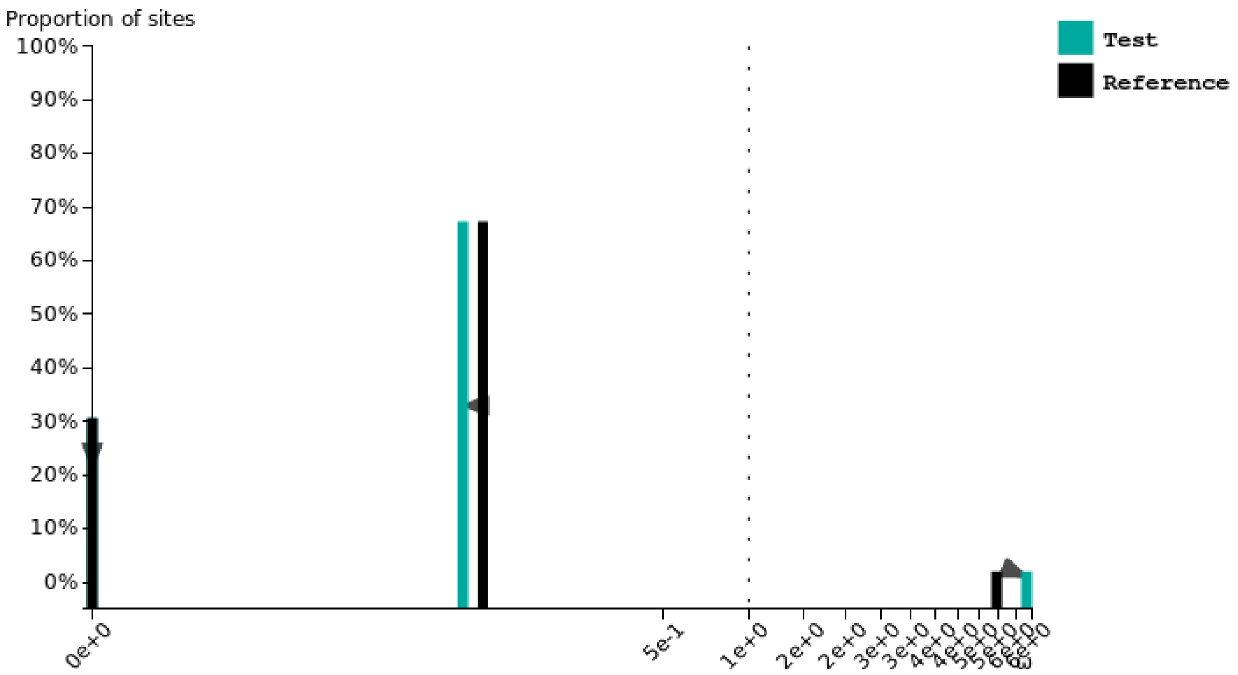
