## Supplementary fig. S16 for "Contrasted gene decay in subterranean vertebrates: insights from cavefishes and fossorial mammals"

### Astyanax mexicanus SF

Test for selection **relaxation** (K = **0.94**) was **not significant** (p = **0.081**, LR = **3.05**).

See here for more information about this method.

Please cite PMID 123456789 if you use this result in a publication, presentation, or other scientific work.

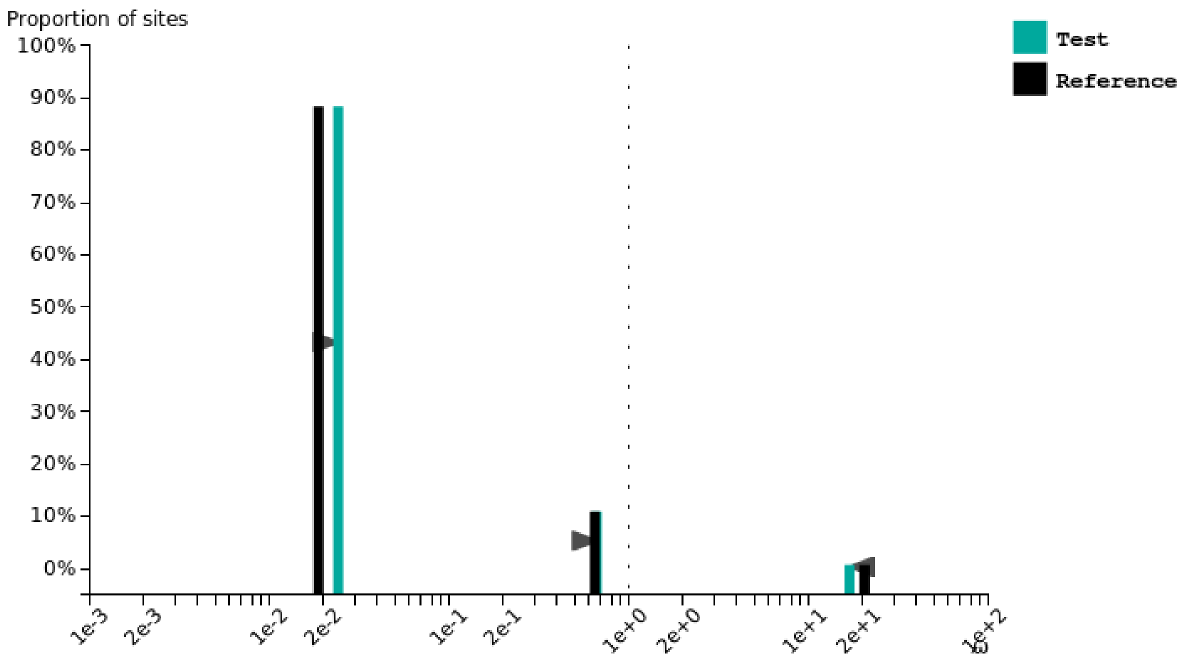


### Astyanax mexicanus CF

Test for selection **relaxation** (K = **1.00**) was **not significant** (p = **1.000**, LR = **0.00**).

See here for more information about this method.

Please cite PMID 123456789 if you use this result in a publication, presentation, or other scientific work.

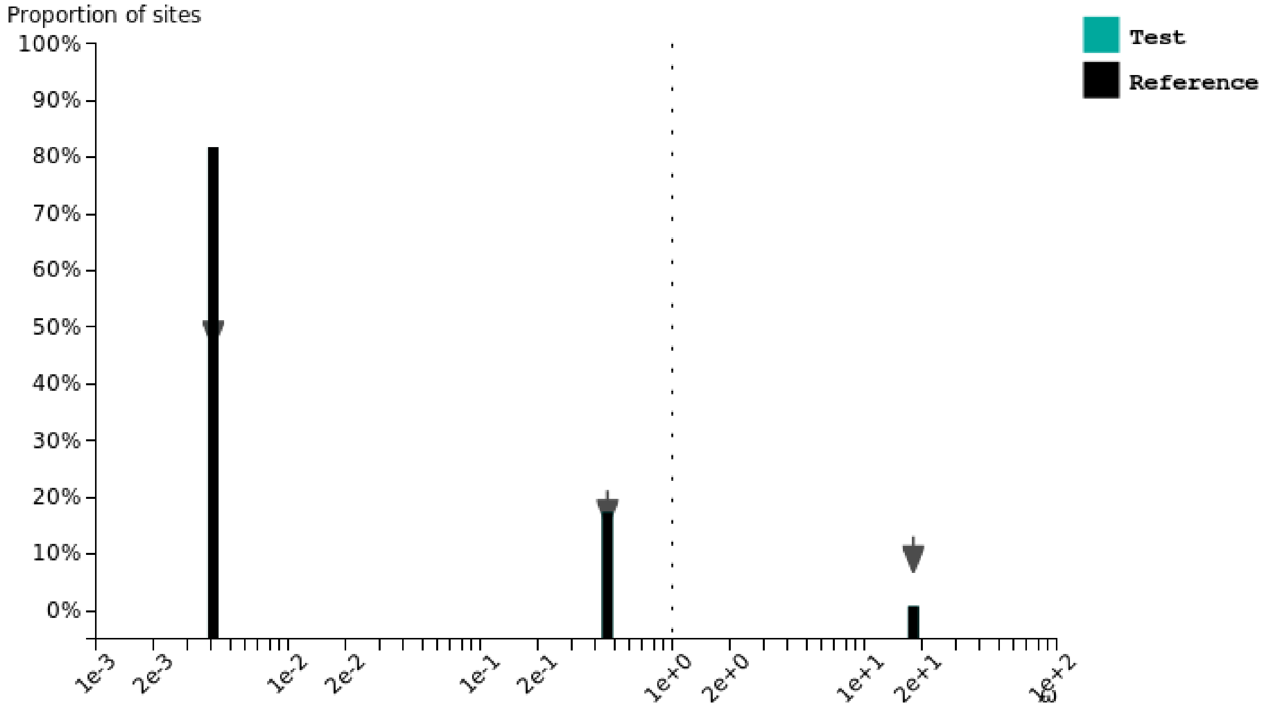

### Brotula barbata

Test for selection **relaxation** (K = **0.98**) was **not significant** (p = **0.304**, LR = **1.06**).

See here for more information about this method.

Please cite PMID 123456789 if you use this result in a publication, presentation, or other scientific work.

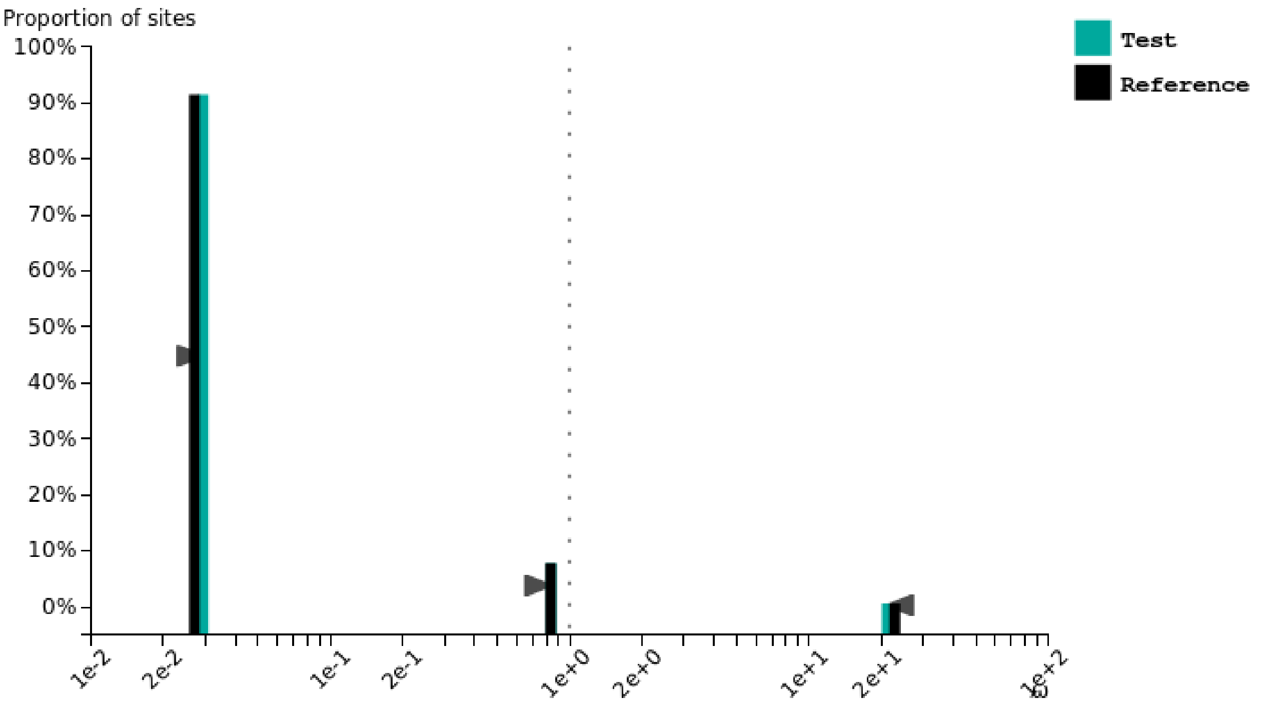


### Carapus acus

Test for selection **intensification** (K = **1.19**) was **significant** (p = **0.000**, LR = **228.78**).

See here for more information about this method.

Please cite PMID 123456789 if you use this result in a publication, presentation, or other scientific work.

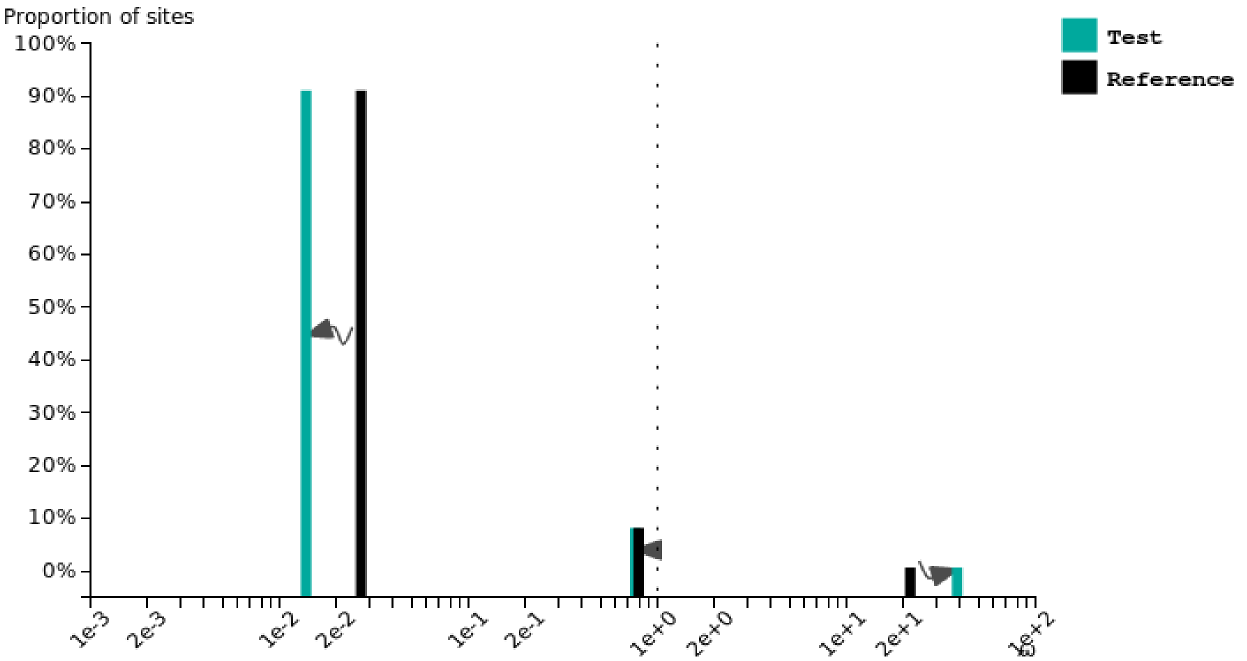

### Lamprogrammus exutus

Test for selection **intensification** (K = **1.10**) was **significant** (p = **0.003**, LR = **8.74**).

See here for more information about this method.

Please cite PMID 123456789 if you use this result in a publication, presentation, or other scientific work.

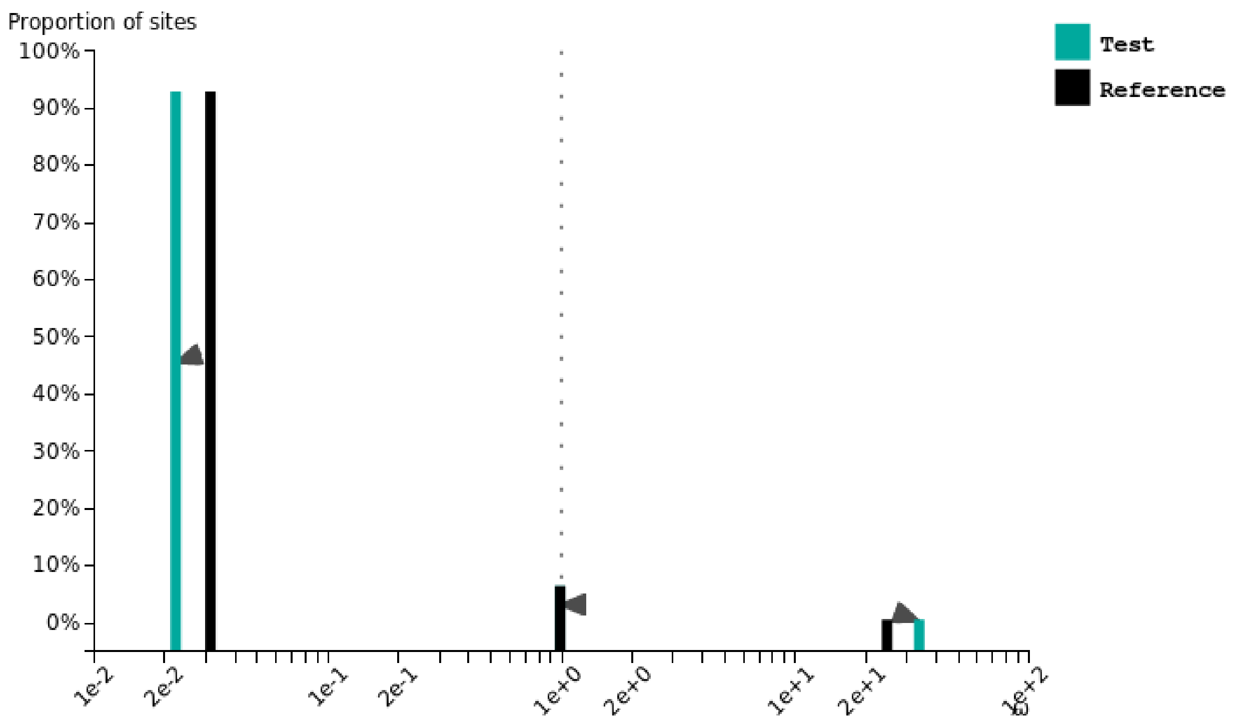


### Lucifuga gibarensis

Test for selection **relaxation** (K = **0.75**) was **significant** (p = **0.000**, LR = **33.52**).

See here for more information about this method.

Please cite PMID 123456789 if you use this result in a publication, presentation, or other scientific work.

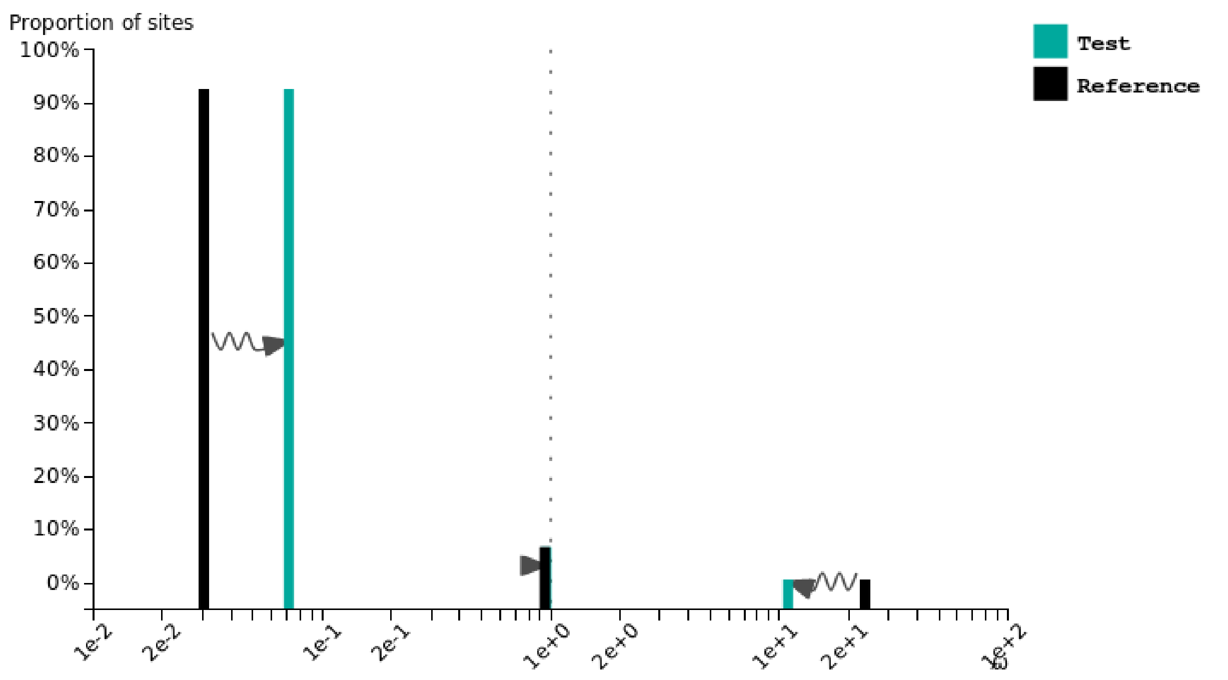

### Lucifuga dentata

Test for selection **relaxation** (K = **0.48**) was **significant** (p = **0.000**, LR = **42.53**).

See here for more information about this method.

Please cite PMID 123456789 if you use this result in a publication, presentation, or other scientific work.

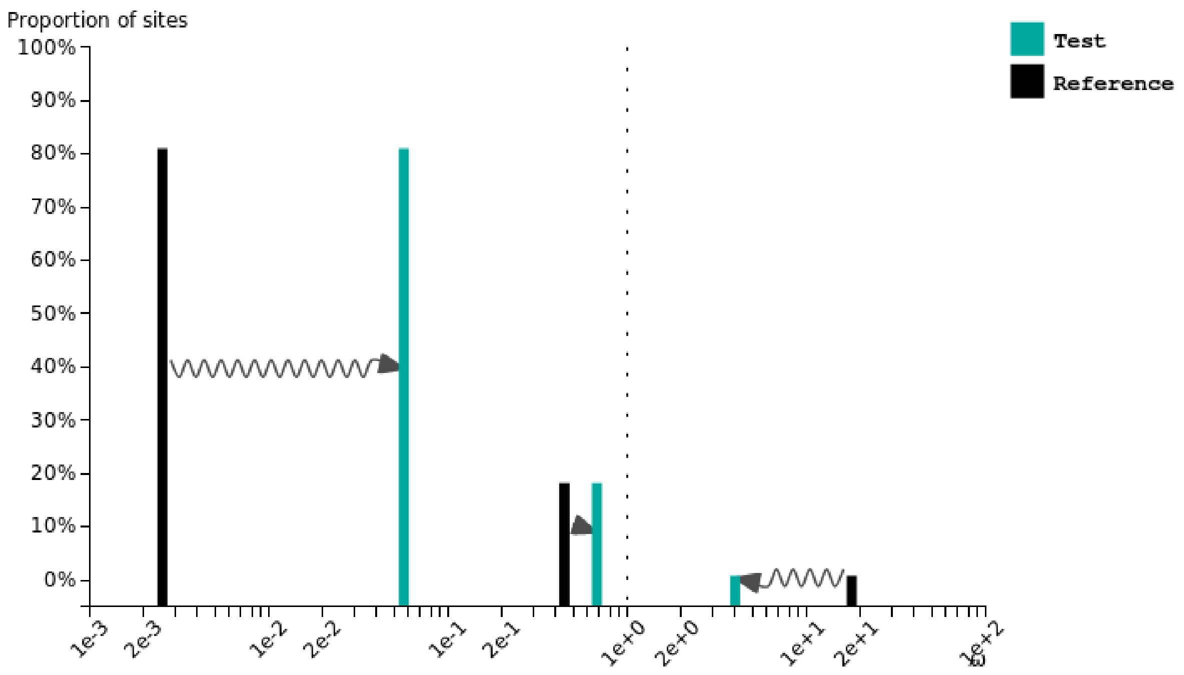


### Danio rerio

Test for selection **intensification** (K = **1.02**) was **significant** (p = **0.046**, LR = **3.99**).

See here for more information about this method.

Please cite PMID 123456789 if you use this result in a publication, presentation, or other scientific work.

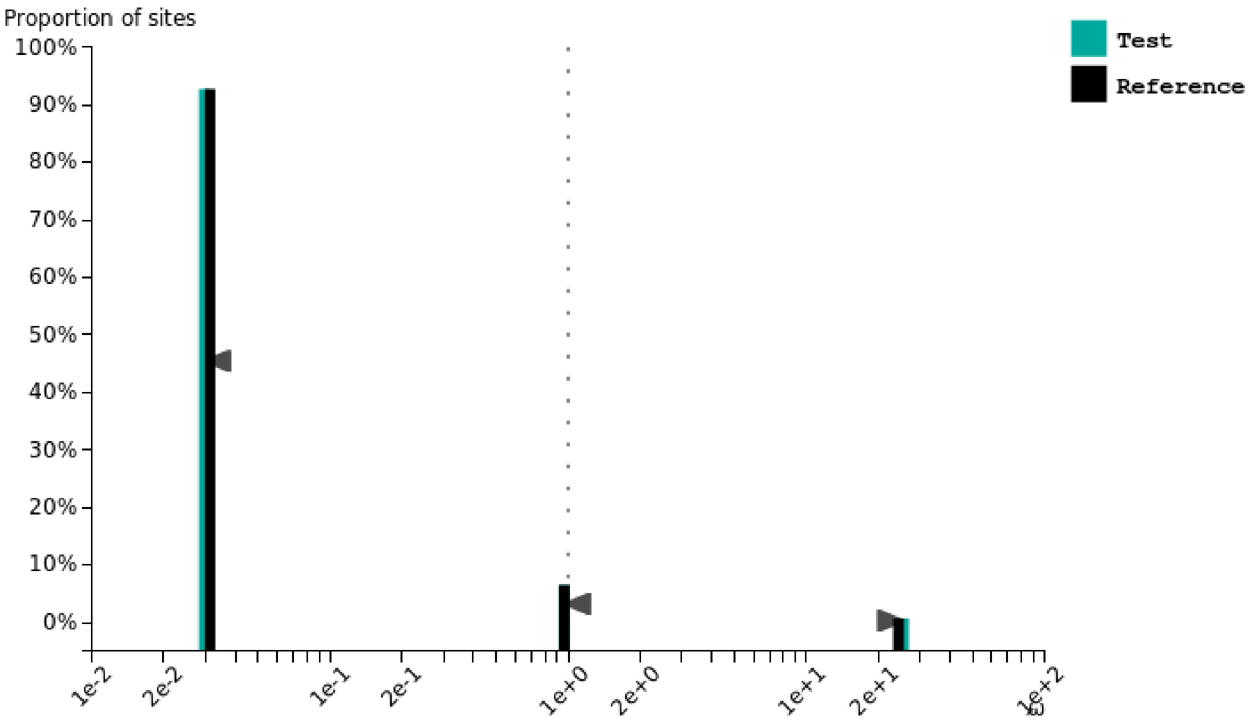

### Dichotomyctere nigroviridis

Test for selection **relaxation** (K = **0.76**) was **significant** (p = **0.000**, LR = **213.59**).

See here for more information about this method.

Please cite PMID 123456789 if you use this result in a publication, presentation, or other scientific work.

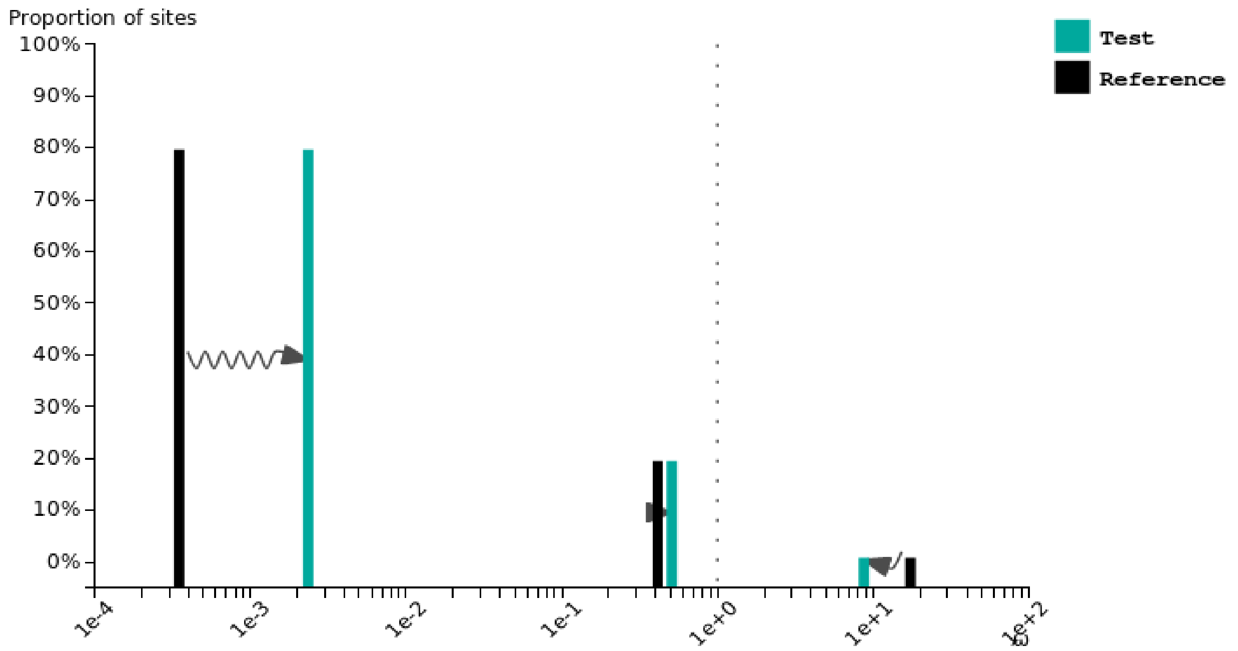


### Gadus morhua

Test for selection **relaxation** (K = **0.98**) was **not significant** (p = **0.249**, LR = **1.33**).

See here for more information about this method.

Please cite PMID 123456789 if you use this result in a publication, presentation, or other scientific work.

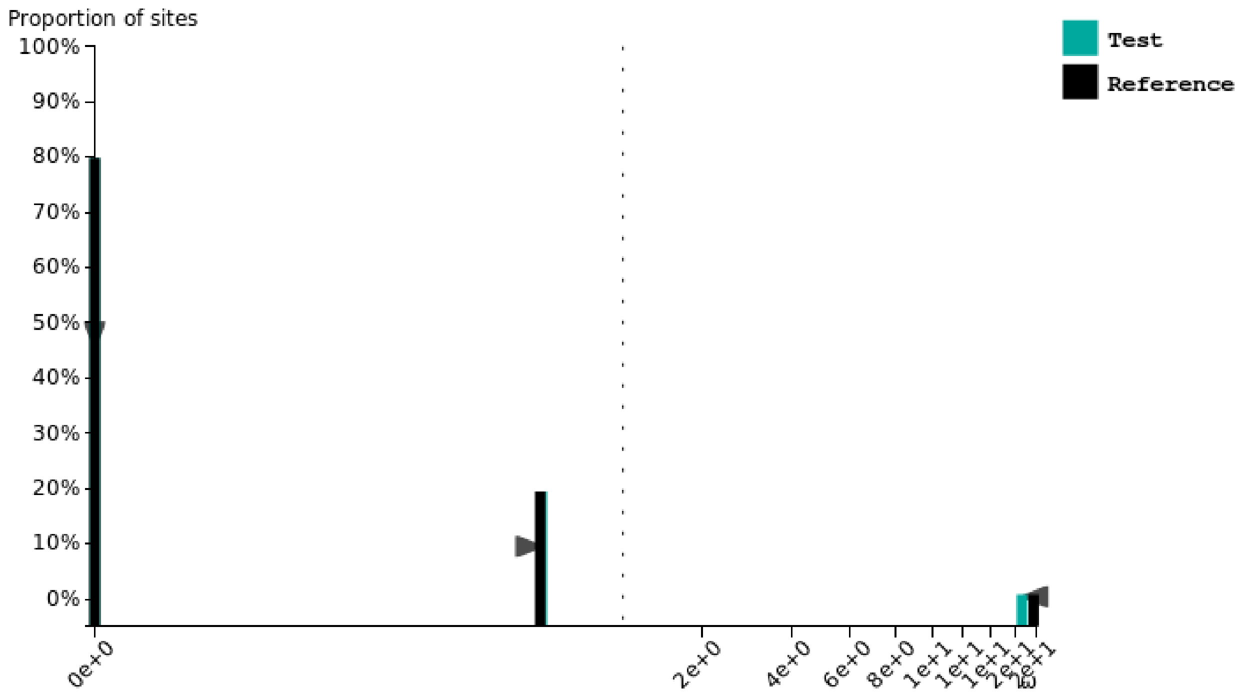

### Gasterosteus aculeatus

Test for selection **relaxation** (K = **0.89**) was **significant** (p = **0.000**, LR = **116.69**).

See here for more information about this method.

Please cite PMID 123456789 if you use this result in a publication, presentation, or other scientific work.

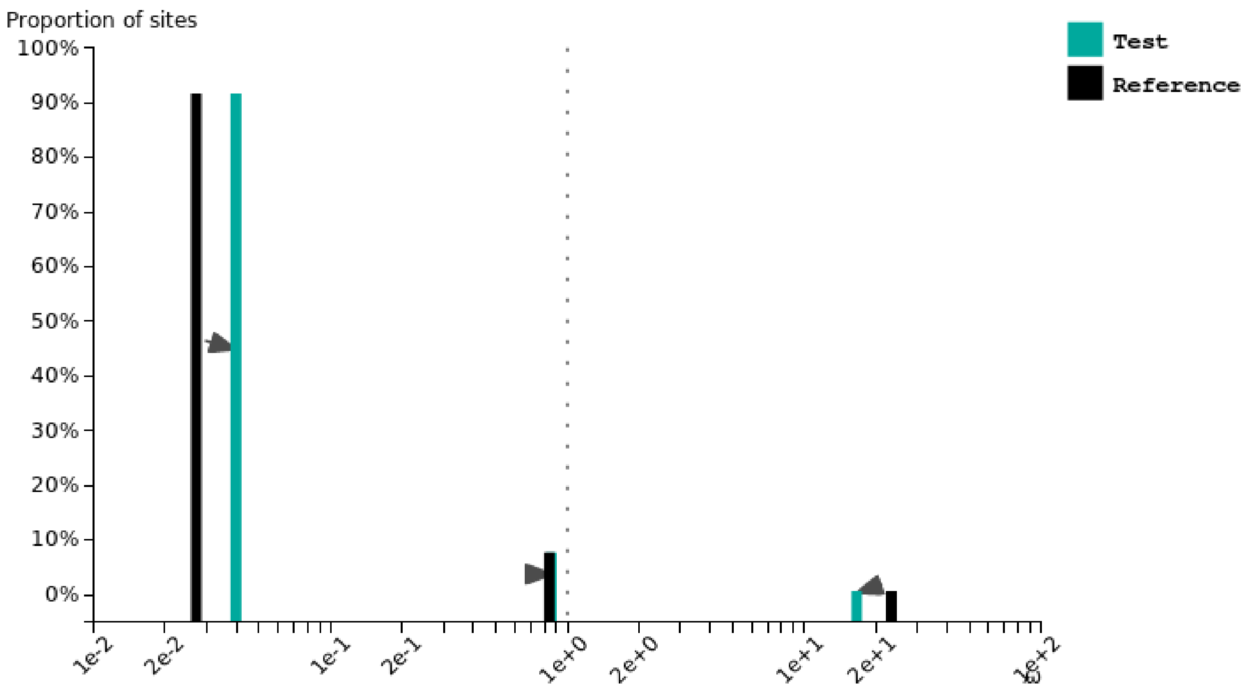


### Oreochromis niloticus

Test for selection **relaxation** (K = **0.96**) was **significant** (p = **0.000**, LR = **14.71**).

See here for more information about this method.

Please cite PMID 123456789 if you use this result in a publication, presentation, or other scientific work.

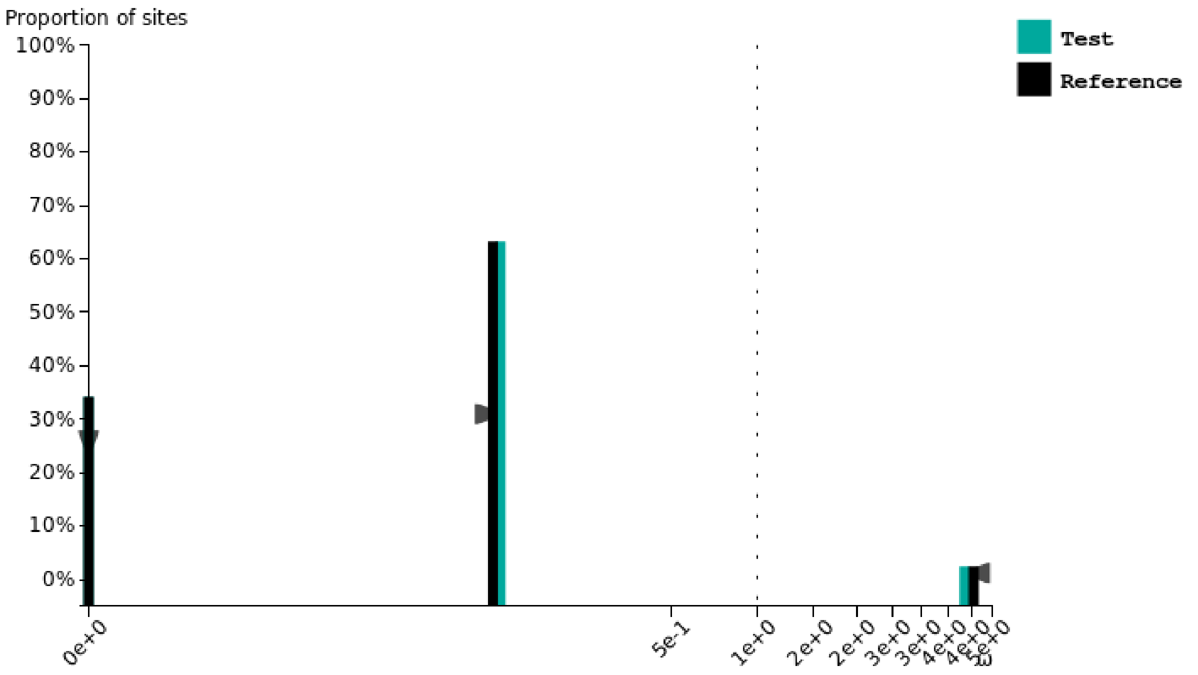

### Oryzias latipes

Test for selection **relaxation** (K = **0.93**) was **significant** (p = **0.000**, LR = **40.23**).

See here for more information about this method.

Please cite PMID 123456789 if you use this result in a publication, presentation, or other scientific work.

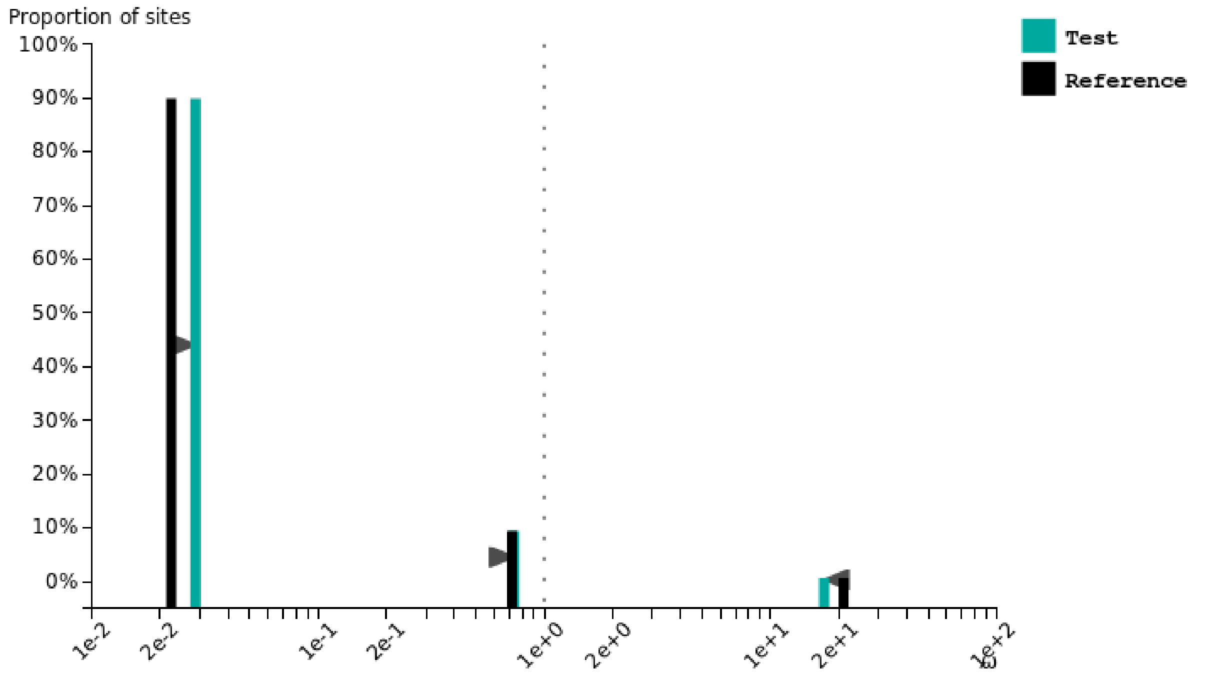


### Xiphophorus maculatus

Test for selection **relaxation** (K = **0.96**) was **significant** (p = **0.000**, LR = **12.37**).

See here for more information about this method.

Please cite PMID 123456789 if you use this result in a publication, presentation, or other scientific work.

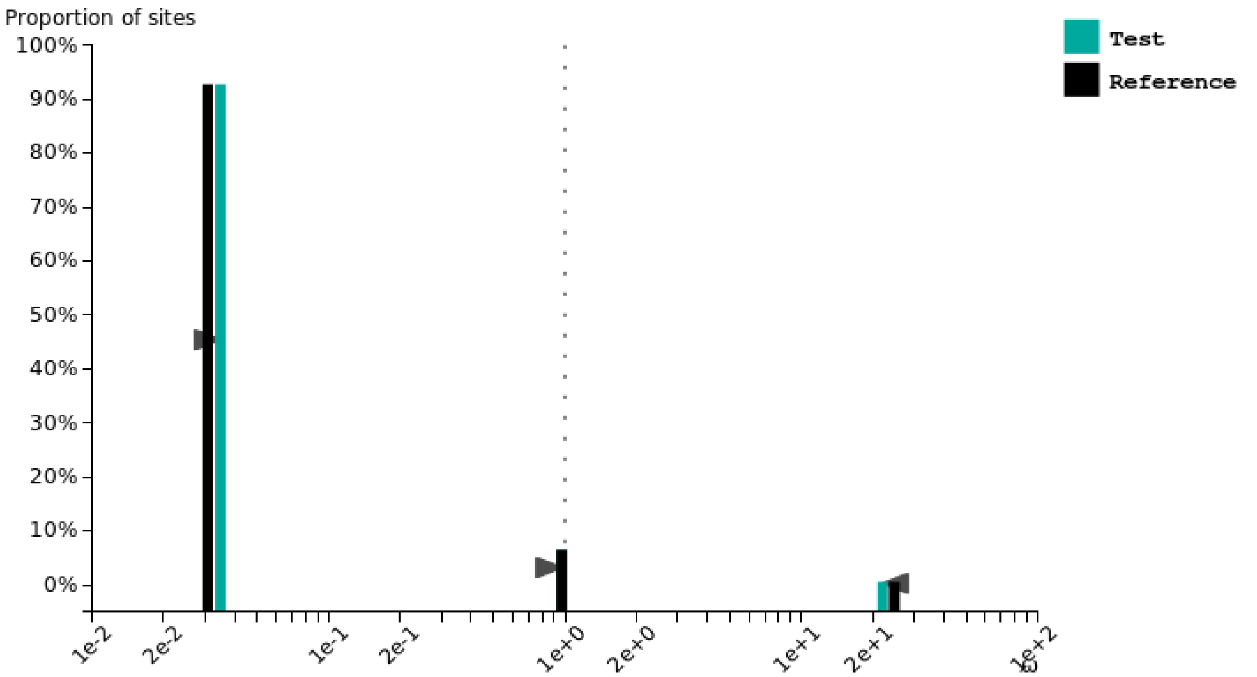

### Pygocentrus nattereri

Test for selection **intensification** (K = **1.33**) was **significant** (p = **0.000**, LR = **172.11**).

See here for more information about this method.

Please cite PMID 123456789 if you use this result in a publication, presentation, or other scientific work.

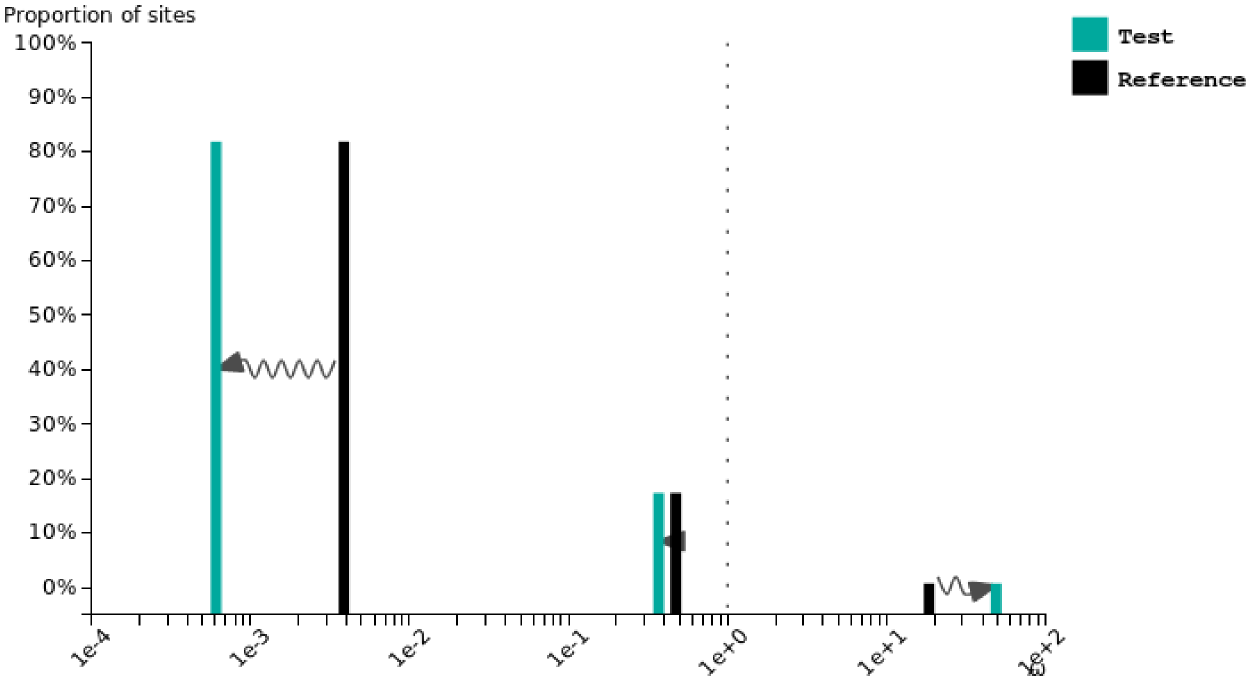
