## Supplementary fig. S17 for "Contrasted gene decay in subterranean vertebrates: insights from cavefishes and fossorial mammals"

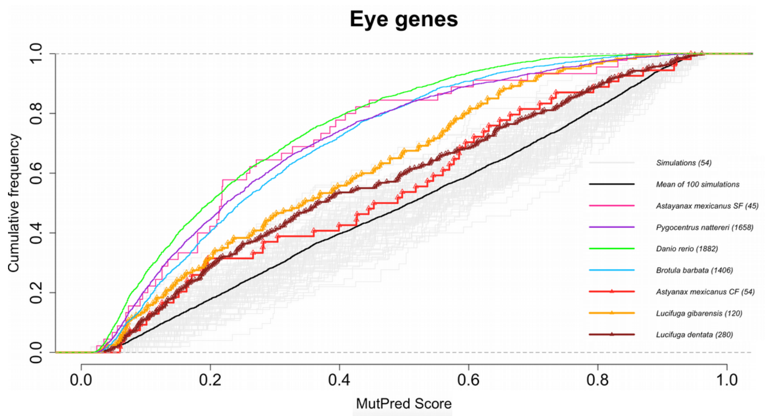


**Kolmogorov-Smirnov Tests p-values**

| | Simu | A.m SF | P.n | D.r | B.b | A.m CF | L.g | L.d |
|---|---|---|---|---|---|---|---|---|
| Simu | 1.0 | | | | | | | |
| A.m SF | $1e^{-6}$ | 1.0 | | | | | | |
| P.n | $3e^{-16}$ | 0.9 | 1.0 | | | | | |
| D.r | $3e^{-16}$ | 0.4 | $4e^{-3}$ | 1.0 | | | | |
| B.b | $3e^{-16}$ | 0.4 | $2e^{-2}$ | $2e^{-9}$ | 1.0 | | | |
| A.m CF | 0.2 | $9e^{-4}$ | $9e^{-6}$ | $3e^{-7}$ | $3e^{-5}$ | 1.0 | | |
| L.g | $5e^{-6}$ | $5e^{-2}$ | $5e^{-4}$ | $3e^{-6}$ | $1e^{-3}$ | 0.3 | 1.0 | |
| L.d | $2e^{-5}$ | $3e^{-3}$ | $9e^{-13}$ | $3e^{-16}$ | $2e^{-12}$ | 0.5 | 0.1 | 1.0 |

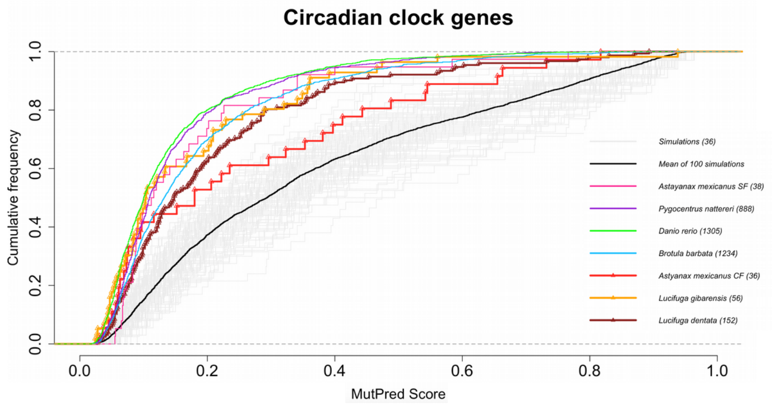


**Kolmogorov-Smirnov Tests p-values**

| | Simu | A.m SF | P.n | D.r | B.b | A.m CF | L.g | L.d |
|---|---|---|---|---|---|---|---|---|
| Simu | 1.0 | | | | | | | |
| A.m SF | $9e^{-5}$ | 1.0 | | | | | | |
| P.n | $3e^{-16}$ | 0.6 | 1.0 | | | | | |
| D.r | $3e^{-16}$ | 0.1 | $2e^{-4}$ | 1.0 | | | | |
| B.b | $3e^{-16}$ | 0.7 | $4e^{-7}$ | $5e^{-12}$ | 1.0 | | | |
| A.m CF | $2e^{-2}$ | 0.2 | $2e^{-2}$ | $5e^{-3}$ | 0.1 | 1.0 | | |
| L.g | $2e^{-6}$ | 0.3 | 0.3 | 0.2 | 0.2 | 0.3 | 1.0 | |
| L.d | $2e^{-9}$ | 0.4 | $1e^{-4}$ | $2e^{-5}$ | 0.3 | 0.2 | 0.1 | 1.0 |

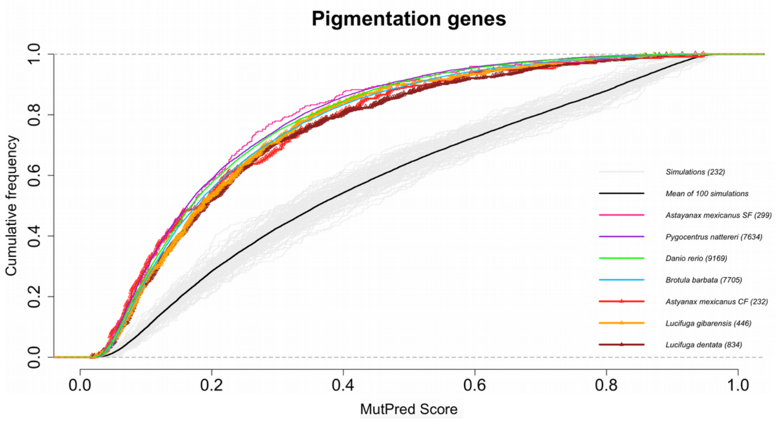


**Kolmogorov-Smirnov Tests p-values**

| | Simu | A.m SF | P.n | D.r | B.b | A.m CF | L.g | L.d |
|---|---|---|---|---|---|---|---|---|
| Simu | 1.0 | | | | | | | |
| A.m SF | $3e^{-16}$ | 1.0 | | | | | | |
| P.n | $3e^{-16}$ | 0.8 | 1.0 | | | | | |
| D.r | $3e^{-16}$ | 0.6 | $3e^{-3}$ | 1.0 | | | | |
| B.b | $3e^{-16}$ | 0.2 | $7e^{-6}$ | $3e^{-2}$ | 1.0 | | | |
| A.m CF | $3e^{-16}$ | 0.1 | 0.2 | 0.2 | 0.6 | 1.0 | | |
| L.g | $3e^{-16}$ | 0.2 | $3e^{-2}$ | 0.1 | 0.5 | 0.4 | 1.0 | |
| L.d | $3e^{-16}$ | 0.1 | $2e^{-4}$ | $6e^{-3}$ | 0.1 | 0.3 | 0.5 | 1.0 |
