## Supplementary fig. S21 for "Contrasted gene decay in subterranean vertebrates: insights from cavefishes and fossorial mammals"

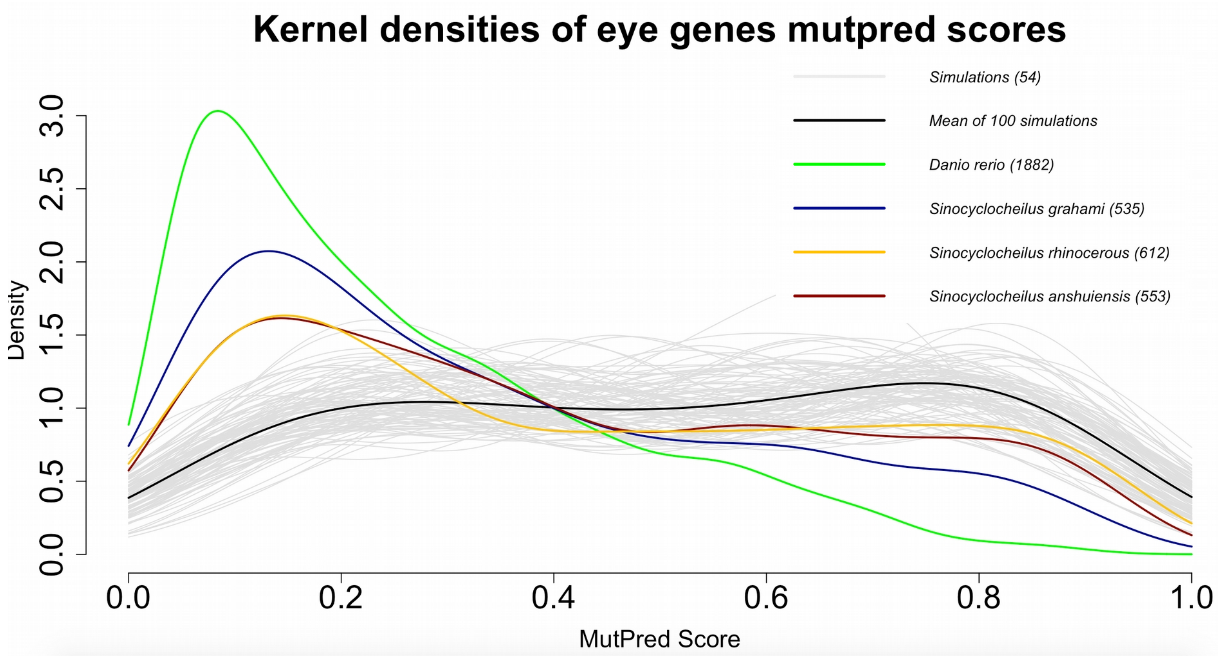

Kernel densities of eye genes mutpred scores
Simulations (54)
Mean of 100 simulations
Danio rerio (1882)
Sinocyclocheilus grahami (535)
Sinocyclocheilus rhinocerous (612)
Sinocyclocheilus anshuiensis (553)
Density
0.0 0.5 1.0 1.5 2.0 2.5 3.0
0.0 0.2 0.4 0.6 0.8 1.0
MutPred Score


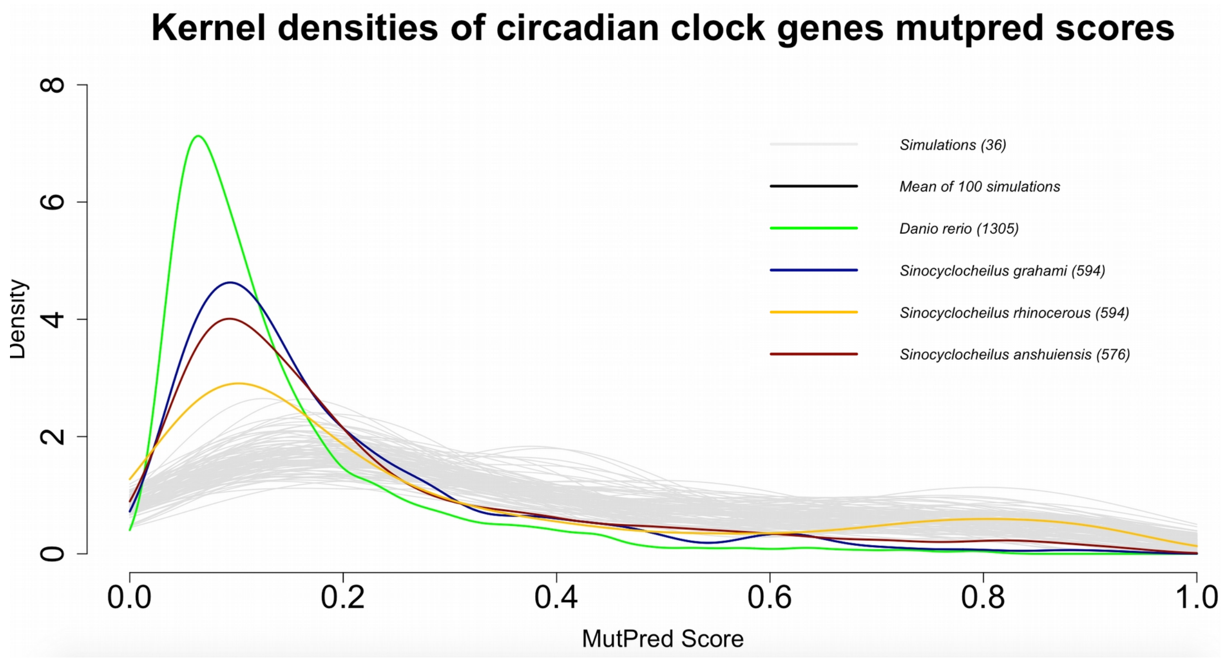

Kernel densities of circadian clock genes mutpred scores
Simulations (36)
Mean of 100 simulations
Danio rerio (1305)
Sinocyclocheilus grahami (594)
Sinocyclocheilus rhinocerous (594)
Sinocyclocheilus anshuiensis (576)
Density
0 2 4 6 8
0.0 0.2 0.4 0.6 0.8 1.0
MutPred Score


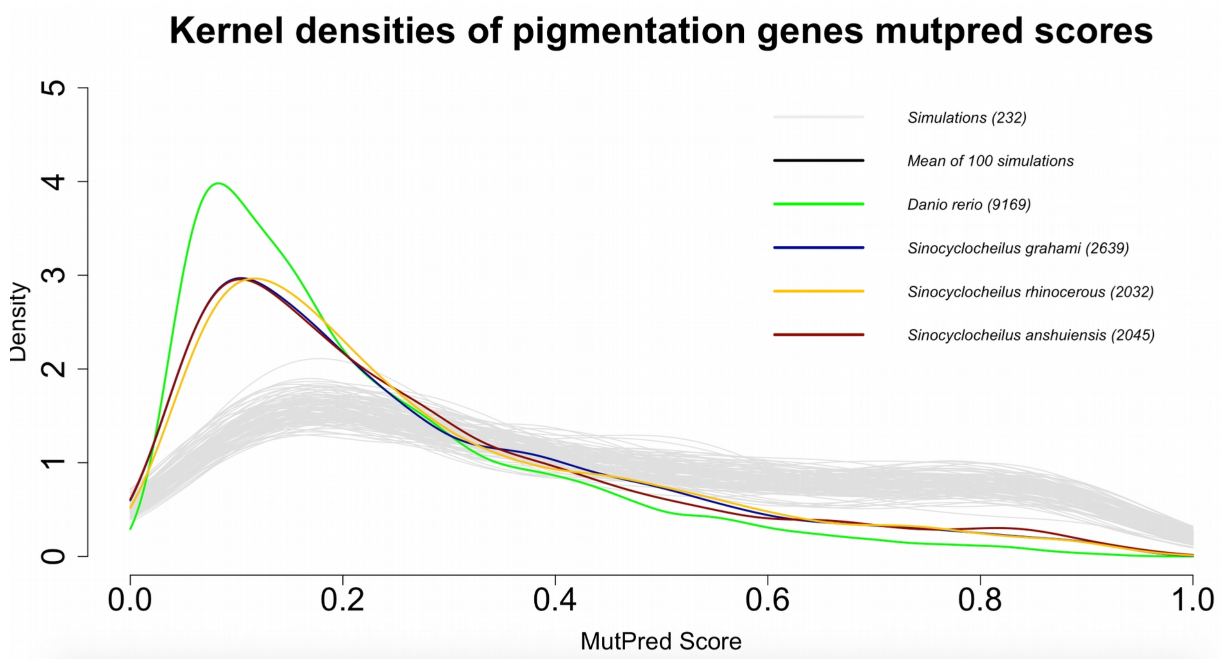

Kernel densities of pigmentation genes mutpred scores
Simulations (232)
Mean of 100 simulations
Danio rerio (9169)
Sinocyclocheilus grahami (2639)
Sinocyclocheilus rhinocerous (2032)
Sinocyclocheilus anshuiensis (2045)
Density
0 1 2 3 4 5
0.0 0.2 0.4 0.6 0.8 1.0
MutPred Score

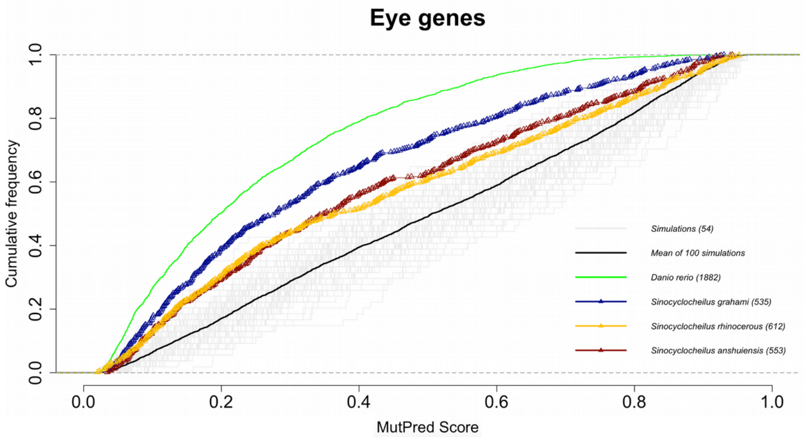


Kolmogorov-Smirnov Tests p-values

| | Simu | D.r | S.g | S.r | S.a |
|---|---|---|---|---|---|
| Simu | 1.0 | | | | |
| D.r | $3e^{-16}$ | 1.0 | | | |
| S.g | $3e^{-16}$ | $3e^{-8}$ | 1.0 | | |
| S.r | $5e^{-13}$ | $3e^{-16}$ | $3e^{-5}$ | 1.0 | |
| S.a | $6e^{-14}$ | $3e^{-16}$ | $2e^{-3}$ | 0.5 | 1.0 |

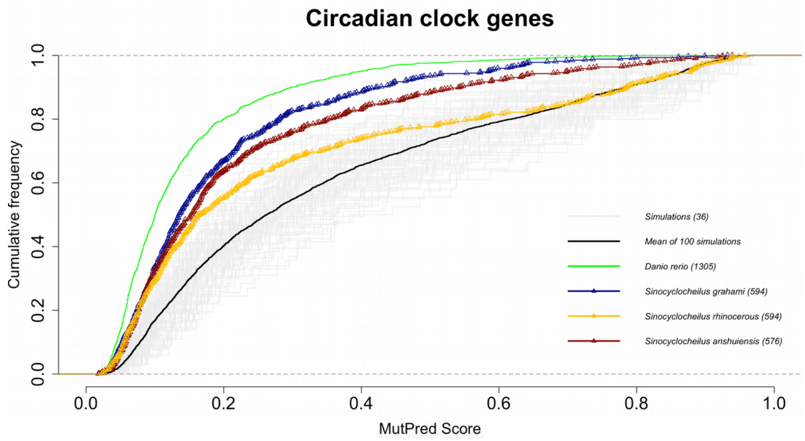


Kolmogorov-Smirnov Tests p-values

| | Simu | D.r | S.g | S.r | S.a |
|---|---|---|---|---|---|
| Simu | 1.0 | | | | |
| D.r | $3e^{-16}$ | 1.0 | | | |
| S.g | $3e^{-16}$ | $2e^{-11}$ | 1.0 | | |
| S.r | $2e^{-16}$ | $3e^{-16}$ | $4e^{-7}$ | 1.0 | |
| S.a | $3e^{-16}$ | $6e^{-16}$ | 0.1 | $7e^{-4}$ | 1.0 |

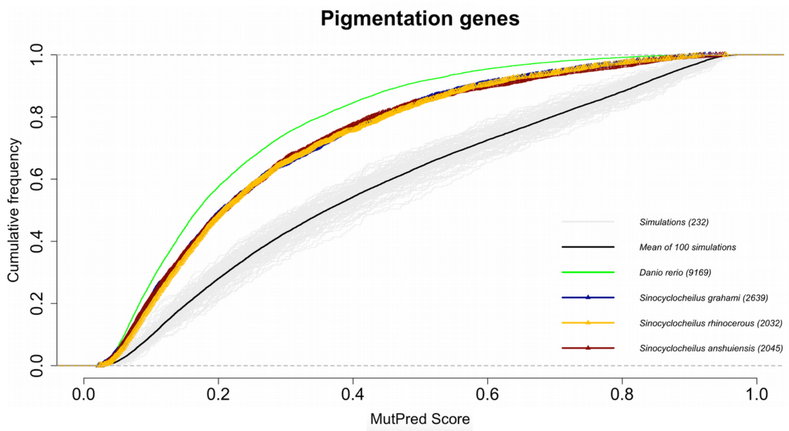


Kolmogorov-Smirnov Tests p-values

| | Simu | D.r | S.g | S.r | S.a |
|---|---|---|---|---|---|
| Simu | 1.0 | | | | |
| D.r | $3e^{-16}$ | 1.0 | | | |
| S.g | $3e^{-16}$ | $3e^{-16}$ | 1.0 | | |
| S.r | $3e^{-16}$ | $1e^{-15}$ | 0.3 | 1.0 | |
| S.a | $3e^{-16}$ | $8e^{-14}$ | 0.9 | 0.5 | 1.0 |
